# Shared Maturation Pathways of HIV-1 Envelope-reactive V3-glycan bnAb Lineages in Human and Rhesus Macaque

**DOI:** 10.64898/2026.01.29.702664

**Authors:** Matthew Clark, Hui Li, Mitchell Martin, Tyler D. Evangelous, Sophie M.-C. Gobeil, Madison Berry, Kshitij Wagh, Elena E. Giorgi, Michael P. Hogarty, Chengyan Zhao, Shuyi Wang, Lorie Marchitto, Wenge Ding, Younghoon Park, Juliette Rando, John Carey, Andrew J. Connell, Ashwin N. Skelly, Yue Chen, Seth Rohr, Chuancang Jiang, Sravani Venkatayogi, Katayoun Mansouri, Robert J. Edwards, Jared Lindenberger, Bhavna Hora, Nicole Doria-Rose, John Mascola, Marit J. van Gils, Rogier W. Sanders, Bette Korber, Beatrice Hahn, Kevin O. Saunders, Priyamvada Acharya, Kevin Wiehe, Barton F. Haynes, George M. Shaw, Wilton B. Williams

## Abstract

Understanding broadly neutralizing antibody (bnAb) lineage development in rhesus macaques (RMs) infected with Simian-HIV (SHIV) may inform HIV-1 vaccine designs. We analyzed HIV-1 envelope-antibody coevolution in 18 RMs infected by SHIV.BG505 (subtype A) and found highly conserved patterns of antibody recognition and Env escape, including in three animals that developed V3-glycan bnAbs. From one RM with V3-glycan targeted plasma breadth, we isolated 203 members of a single clonal Ab lineage designated DH1030. DH1030 Abs demonstrated striking genetic, functional and structural similarities with the human V3-glycan bnAb lineage DH270, which was isolated from an individual with subtype C HIV-1 CH848 infection. Notably, human-DH270 and macaque-DH1030 bnAbs shared early improbable mutations in HCDR2 that were critical for bnAb development. Thus, key improbable mutations and convergent patterns of antibody evolution and epitope recognition were shared across primate species and distinct HIV-1 subtypes, findings that may be leveraged in new HIV-1 vaccine designs.

**ONE SENTENCE SUMMARY:** Broadly neutralizing antibodies targeting HIV-1 envelope surface protein develop via a common maturation pathway across primate phylogeny.

## Introduction

A major goal of a vaccine against human immunodeficiency virus-type 1 (HIV-1) is the induction of broadly neutralizing antibodies (bnAbs) that target conserved regions on the envelope (Env) surface protein ^1^. Multiple bnAb epitopes have been described on HIV-1 Env as targets for HIV-1 vaccination ^2, 3^. The V3-glycan bnAb epitope supersite consists of high mannoses in the glycan-rich apex of Env in combination with a peptide motif (GDIR) at the base of the third variable loop (V3 loop). This epitope supersite is targeted by bnAbs that use different immunogenetics and structural angles of approach for antigen recognition ^4–8^. Moreover, potent V3-glycan bnAbs have been isolated from people living HIV-1 (PLWH) ^9–11^ and from SHIV-infected monkeys ^4, 12, 13^. Thus, V3-glycan bnAbs are a desired target for HIV-1 vaccines.

We previously isolated a V3-glycan bnAb lineage termed DH270 from an individual (CH848) with chronic HIV-1 infection ^8^. Isolation of DH270 lineage Abs enabled the inference and characterization of the DH270 unmutated common ancestor (UCA) and intermediate antibody (IA) lineage members to understand bnAb lineage maturation. DH270 lineage progression from the UCA to IA and mature Abs was associated with the acquisition of an arginine residue at position 57 (R57) in the heavy chain complementarity determining region-2 (HCDR2) that displaced the Env V1 loop while accessing the GDIR motif in the V3-glycan supersite ^14–16^. The emergence of R57 was a result of a key improbable mutation (G57R) associated with DH270 neutralization breadth ^8^. Moreover, Ab-Env co-evolution studies in the CH848 individual identified an autologous Env with V1 glycan deletions that was postulated to have primed the DH270 lineage ^8^, and subsequent immunogens based on this design successfully expanded DH270 lineage B cells in mouse models ^15, 17, 18^. Additionally, previous studies have reported HIV-1 Envs with V1 glycan deletions as candidate immunogens for eliciting V3-glycan-like B cells in animal models ^4, 19–21^.

A potential obstacle for vaccine-induction of V3-glycan bnAbs is the dearth of precursor B cells ^19, 22^. Nonetheless, HIV-1 Envs bearing V1-glycan deletions when used as B cell baits have shown promise for isolation of V3-glycan-like precursor B cells in HIV-1 naïve humans ^19,23^. Thus, understanding commonalities among V3-glycan bnAb lineages that may be exploited by different immunogens may inform vaccination strategies to elicit them. More recently, RMs infected with pathogenic SHIVs ^24^ induced HIV-1 Env bnAbs and recapitulated Env-Ab co-evolution events, including Env mutations that select for bnAb lineage induction, expansion and affinity maturation ^4, 12^. This macaque model of bnAb induction thus provides a molecular guide for bnAb induction via vaccination by employing similar Ab-Env coevolution studies as those performed in human HIV-1 infection ^8, 25^.

Germinal centers (GC) play a critical role in development of antigen-reactive B cell responses ^26–29^, but the cellular mechanisms of bnAb development in macaque GCs remain poorly defined. Antigen-activated GC precursor B cells establish the dark zone in which they undergo rounds of proliferation and somatic hypermutation (SHM), followed by migration to the light zone for positive selection as plasmablasts, memory precursor B cells, or undergo dark zone recirculation for further SHM ^30–33^. HIV-1 Env bnAbs are generally thought to require high levels of SHM, including improbable mutations to achieve optimal neutralization potency and breadth ^14^. However, the dynamics of bnAb B cells in GC dark and light zones during lineage maturation, including acquisition of functional improbable mutations associated with neutralization potency and breadth of HIV-1 Env bnAbs, are largely unknown.

Here we studied the induction and evolution of HIV-1 Env V3-glycan bnAbs in macaque SHIV infection compared to human HIV-1 infection to search for shared mechanisms of affinity maturation across primate phylogeny targeting different HIV-1 subtypes. In this study, human and macaque V3-glycan bnAbs were elicited by different Envs (subtype A BG505 in rhesus; subtype C CH848 in human). For the SHIV.BG505 infected macaque, we identified a subset of viruses containing deletions in V1 that circulated coincident with priming of the DH1030 bnAb lineage, similar to the scenario postulated for DH270 priming ^15, 34^ and for SHIV.5MUT induced bnAbs ^4^. Surprisingly, we then identified an early shared key functional improbable mutation in the HCDR2 of both rhesus and human bnAb lineages that conferred similar Ab structure and epitope recognition. To investigate how two widely divergent HIV-1 Envs infecting two different primate species could share common developmental pathways, we analyzed the maturation pathway of the macaque V3-glycan bnAb lineage by performing longitudinal B cell receptor (BCR) and transcriptome sequencing of B cell compartments in peripheral blood cells (PBMCs) and secondary lymphoid tissues, including lymph node (LN), spleen, and bone marrow (BM). Macaque V3-glycan bnAb BCR sequences clustered into different phylogenetic sub-clades distinguished by acquisition of the functional improbable mutation that appeared to be selected in LN GC—a desired feature of HIV-1 vaccine regimens to shape bnAb lineage development ^14,35^. Our data highlight the value of SHIV-infected macaques as an outbred model system to explore conserved molecular pathways of bnAb development following infection and vaccination.

## Results

### Convergent patterns of Env-Ab coevolution in SHIV.BG505 infected RMs

We inoculated 18 RMs with SHIV.BG505 in this study (**Table 1**). Fourteen RMs were inoculated with SHIV.BG505 bearing a potential N-linked glycosylation site (PNGS) at Env position 332 (SHIV.BG505.N332), two RMs with SHIV.BG505 lacking a PNGS at Env position 332 (SHIV.BG505.T332), and two RMs with a 50:50 mixture of these SHIVs. All animals became productively infected with peak plasma virus titers at day 14 of 10^6^-10^8^ vRNA molecules/ml (**Figure 1A)**. Plasma viral load setpoints at week 12 ranged from 10^2^-10^5^ vRNA molecules/ml. Two monkeys died of rapid disease progression to AIDS within the first six months of infection. Except for RM6447 that died of AIDS-like illness at week 9 post-infection, all monkeys developed autologous strain-specific nAb ID50 titers between 1:40 – 1:10,000 (**Figure 1A**). We defined neutralization breadth as ID_50_ titers of at least 1:20 against 4 or more heterologous viruses in a test panel of 16 tier-2 or difficult-to-neutralize strains. At 48 weeks post-SHIV infection, none of the RMs exhibited neutralization breadth reaching this threshold (**Table S1**). Between weeks 56 and 96, three RMs – 08N021, 10N011 and 6923 – reached this threshold eventually neutralizing 31-88% of the test panel (**Figure 1B; Table S1**). For the three monkeys 08N021, 10N011 and 6923, we mapped plasma neutralization breadth to the canonical V3-glycan bnAb epitope motifs N332 and ^324^GDIR_327_ in heterologous Env backbones (**Figure 1C**). Thus, we found that 3 of 18 monkeys infected by SHIV.BG505 developed V3-glycan bnAbs, but only after more than one year of SHIV infection.

**Figure 1:**
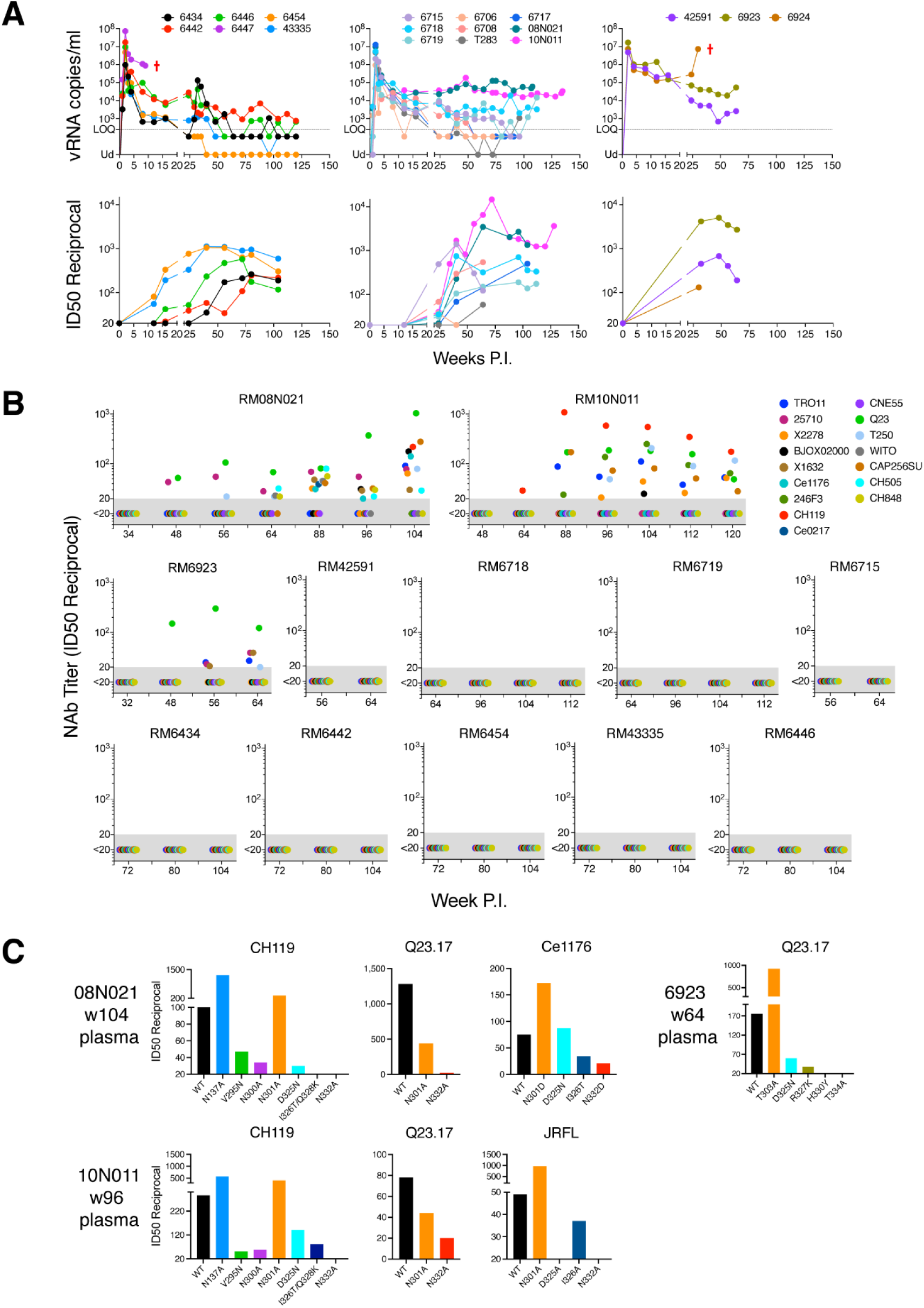
Viral load kinetics and neutralizing antibody responses of SHIV BG505-infected RMs. **(A)** Top panel shows the viral loads. The dashed line indicates the limit of quantification (LOQ) of vRNA detection (LOQ = 250 vRNA copies/mL). Animals with AIDS are indicated by red cross signs (†). Bottom panel shows the ID50 titer (reciprocal) of the autologous neutralizing antibody responses. **(B)** Plasma neutralization against a panel of 16 tier-2 heterologous viruses. **(C)** Mapping of the plasma neutralizing response against mutants lacking key V3-glycan epitope residues in different heterologous backbones. The Y-axes represent reciprocal of ID50.

**TABLE 1.**
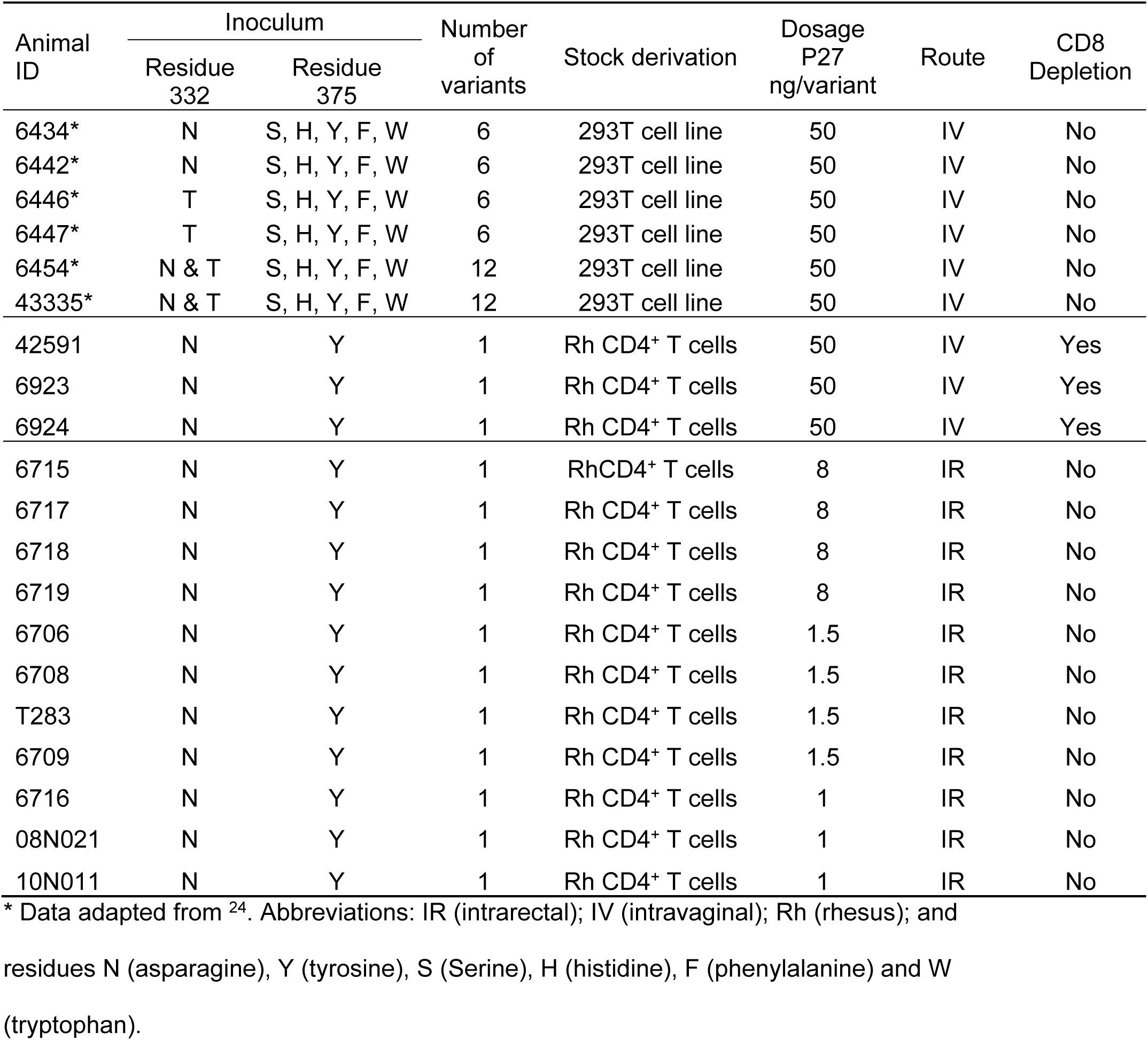

Because BG505 is a prototypic strain used widely in HIV-1 vaccine research, we sought to understand the molecular basis for its early strain-specific immunogenicity ^36^ and how the Env evolved to elicit bnAbs. Circulating virus has a lifespan of less than an hour, and the cells producing it have a lifespan of about a day. This makes the viral quasi-species in plasma a sensitive and dynamic measure of viral fitness and selection pressures, including that exerted by nAbs, cytotoxic T cells and antiretroviral drugs ^37–41^. Single genome sequencing (SGS) provides a proportional representation of viral sequences in plasma and maintains sequence linkage across complete *env* genes, thereby enabling sites of antibody recognition to be mapped across Env ^42, 43^. We thus compared the patterns of Env sequence evolution in 17 of 18 macaques where samples were available and vRNA titers were sufficient to identify signature sites of recognition and escape from autologous strain-specific antibodies and bnAbs targeting the canonical V3-glycan epitope. **Figure S1A** depicts Env gp140 sequence evolution over the course of infection in all monkeys compared with the human subject BG505. We used our previously developed algorithm LASSIE ^44^ to detect Env sites under selection based on loss of transmitted-founder (TF) sequence in longitudinal SGS sequences and found evidence of strong positive selection across all infected monkeys and the human (**Figures 2A-2B, and S2-S4**). As seen previously for macaques infected with SHIVs bearing the same TF Env isolated from a human participant ^12^, there was remarkable convergence in Env evolution with the same or similar amino acid mutations identified under selection across the macaque hosts and the human (**Figures 2A-2C**). Of the 84 total LASSIE selected sites across BG505 and 17 RMs, 36 (43%) were selected in at least 2 human/rhesus hosts (**Figure S4**). These mutations also included recurrent glycan acquisition or deletion mutations that shaped the glycan shield evolution, such as the addition of glycans at sites 241 and 289, which cover an immunodominant “glycan hole” epitope ^45, 46^, and the gain of N332 glycan in BG505 and 2 out of 3 macaques infected with wildtype BG505 that lacks this glycan. When mapped on the trimer structure (**Figure 2C**), the most recurrently selected sites fell in regions of the 241-289 glycan hole, the C3-V4 and V3 regions, in agreement with the presence of strain-specific mAbs in SHIV BG505-infected RMs ^36^. Remarkably, 6 Env sites were selected in 11-13 hosts out of 17 RMs and the BG505 human; each of these sites was either selected in BG505 or showed mutations at lower frequency. Three such sites (347, 356, 358) were in the C3 region, two in the N241-N289 glycan hole (241 and 289) and one at the tip of V3 (R308H) that was selected for in 12 out of 17 RMs. Of note, the two RMs with the highest plasma breadth (RM08N021 and RM10N011) showed the highest overlap with selected Env sites in BG505 (7 and 9, respectively, out of 13 sites in BG505) (**Figures 2A-2C**). Although hypervariable loops evolve rapidly by mutations and indels even in within-host longitudinal Envs, we found several patterns of common indels shared across multiple RMs **(Figure S4**), including a 4-amino acid (aa) deletion in V1 loop.

**Figure 2:**
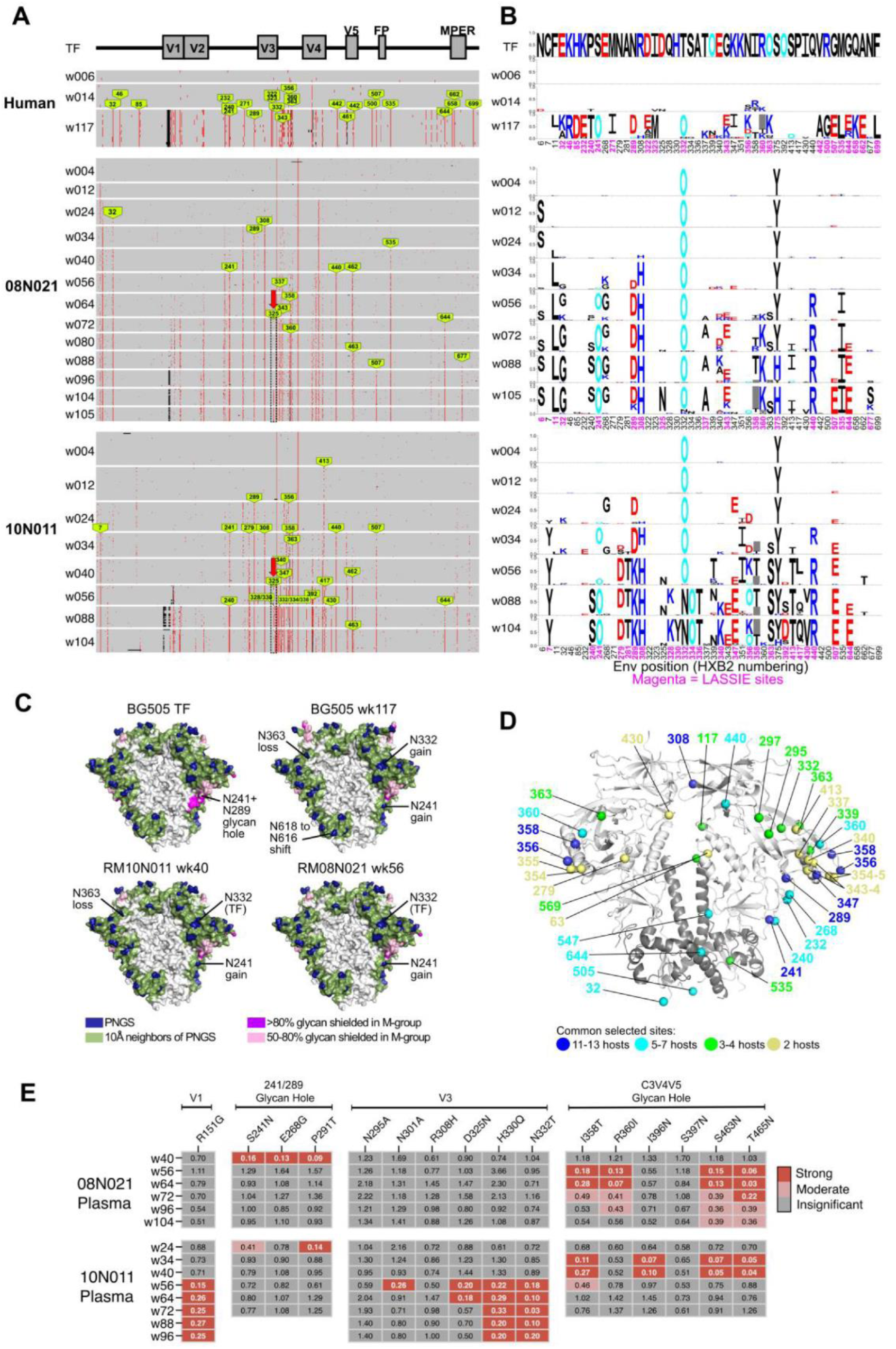
Recapitulation of Env evolution. **(A)** Pixel plots of gp140 SGS of human BG505 and SHIV-BG505 infected RMs 08N021 and 10N011. Green arrows – conserved mutation sites with >50% TF loss and present in ≥ 2 monkeys. Black dotted boxes – V3-glycan bNAb epitope region. **(B)** LASSIE sites (>80% TF loss at ≥1 time point) in each host linked to panel A are labeled in magenta. Mutations (logos); height is proportional to its frequency at a given time point. TF residues are blanked out for longitudinal time points. Wk: week post infection. O: PNGS. **(C)** Common patterns of glycan shield evolution between BG505 and the two RM hosts. **(D)** Structural mapping of Env sites under selection across ≥2 hosts. Selected sites shown as spheres on Env trimer structure PDB:5FYL and are color-coded according to number of hosts they were identified to be under selection. **(E)** Plasma from w24-w104 were tested against V1, 241/289 glycan-hole, V3 and C3V4V5 glycan-hole mutants. The change in ID50 of the mutant relative to the BG505.T332N parental reflects the degree of escape from autologous responses; relative ID50 (RID50s = ID50mutant/ ID50parent). Strong (dark red), moderate (light red) or insignificant (grey) changes in neutralization resistance are indicated.

For many positively selected residues in gp140 in the first year of infection, we could phenotypically map an autologous, strain-specific nAb response (**Figure S1B**), including the previously characterized N241+N289 glycan hole and C3-V4-V5 epitopes from animal immunizations with BG505 Env trimer ^45^. In two of the three RMs with V3-glycan bnAbs (08N021 and 10N011), we could identify selected mutations in the V3-glycan and/or ^324^GDIR_327_ bnAb epitope supersite (**Figures 2A-2B and S5**) and could confirm phenotypically that bnAbs in these animals targeted these epitopes (**Figure 1C**). For RM 6923, while D325N was observed at 60% at the last SGS time point of 48 weeks post infection (**Figure S1A**), this did not meet our prespecified criterion on >80% TF-loss for LASSIE selection. However, this could indicate a late V3-glycan bnAb response since this macaque also showed heterologous nAbs mapping to V3-glycan bnAb epitope at week 64 (**Figure 1C**). Additionally, BG505 Env variants in the human and the macaques that developed V3-glycan bnAbs evolved with common glycan shield patterns shaped by nAb selection pressure (**Figure 2C**). Positively selected sites of mutation across all infected RMs are depicted on a surface projection of the trimer (**Figure 2D**) with neutralization phenotype mapping summarized in **Figures 2D** and **S1B**.

RMs 08N021 and 10N011 exhibited the greatest neutralization breadth and thus we focused on these two monkeys for further analysis. RM 10N011 was previously found to have developed a Fab-dimerized bnAb that bound high-mannose glycans on HIV-1 Env ^47^. In the current study, we sought to isolate and characterize the V3 glycan bnAbs present in RM 08N021 to elucidate their biophysical properties and ontogeny.

### Isolation of V3-glycan bnAbs from RM08N021

From RM08N021, we isolated four PBMC-derived members of a V3-glycan bnAb lineage (DH1030.1, DH1030.2, DH1030.3, and DH1030.4) using an antigen-specific B cell sort approach ^47, 48^. We isolated two additional members of this lineage, DH1030.5 and DH1030.6, from BMA-derived mononuclear cells using an antigen-unbiased plasmablast/ plasma cell sort strategy ^49^. Notably, we found that DH1030.1-DH1030.5 encoded an improbable mutation from glycine to arginine at position 56 (G56R) in the HCDR2 similar to previously identified G57R functional improbable mutation in the human DH270 V3-glycan bnAb lineage (**Figure S6A**) ^8^.

Moreover, this mutation was also critical for heterologous HIV-1 neutralization capacity of DH1030.1 when expressed as a recombinant monoclonal (m) Ab (**Figure S6B**) as similarly described for DH270 V3-glycan bnAb lineage development ^8^. To determine whether the macaque DH1030 and human DH270 V3-glycan bnAbs have similar mechanisms of action shaped by convergent evolution in the HCDR2, we next structurally analyzed the representative DH1030.1 mAb for comparison with DH270.

To visualize the Env epitope of DH1030.1 at high resolution, we determined two cryo-EM structures of the DH1030.1 antigen binding fragment (Fab) bound to BG505 SOSIP or CH848 SOSIP. The cryo-EM maps showed three DH1030.1 Fab molecules bound symmetrically per Env trimer. Applying 3-fold symmetry resulted in a 3.5 Å resolution map for the BG505-DH1030.1 complex and a 3.7 Å resolution map for the CH848-DH1030.1 complex (**Figures 3A, S7 and S8; Tables S2 and S3**). We determined the x-ray crystal structure of the DH1030.1 Fab at (1.79 Å resolution) (**Table S2**). DH1030.1 bound at the Env V3-glycan supersite, burying a total of 1207 A^2^ and 1327 A^2^ at its interface with Env for the CH848-bound and BG505-bound complexes, respectively, and making contacts with protein surfaces and glycans, including the N332 glycan (**Figures 3B-3C; Table S4**). DH1030.1 contacts with Env were dominated by its heavy chain with all the CDR loops involved in contacts with Env. By contrast, the light chain DH1030.1 contacts with Env were limited to glycans, with R51, Y54, S61 making contact with the N332 glycan. We found that the conformations of the CDR loops in the free Fab were overall similar to the CDR conformations of the DH1030.1 Fab bound to Env, with a maximum ∼2.4 Å Cα shift of the CDRH3 region spanning residues 101-109 being sufficient to accommodate binding to Env (**Figure S9**). Similar to known HIV-1 bnAbs targeting the V3-glycan supersite ^1, 35, 50^, DH1030.1 makes close contact with Env N332 glycan, with both its heavy and light chains involved in contacting this glycan. The 17-aa residue long DH1030.1 HCDR3 loop extends into the binding site to contact protein residues Ile322, Gly324, Asp325, Arg327 and Gln328 in the V3 loop, while the Env N332 glycan stacks against the extended DH1030.1 HCDR3 loop (**Figure 3D**). Approximately 60% of the contact surface of N332-glycan with the DH1030.1 Fab comes from the HCDR3 loop with a terminal mannose nestled within a pocket made by the heavy and light chains.

**Figure 3.**
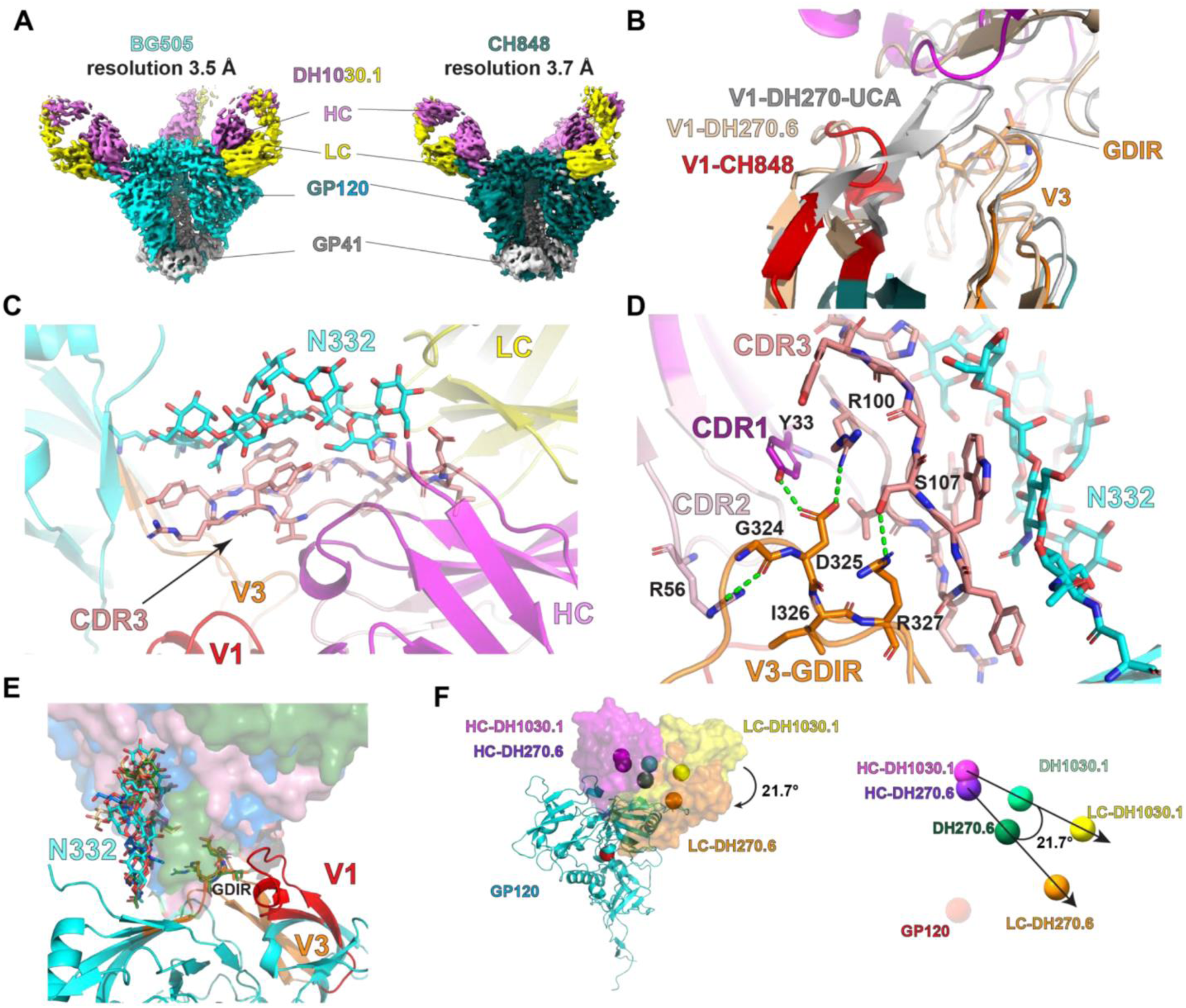
Cryo-EM structures of DH1030.1 complexes. **(A)** CryoEM map of DH1030.1 Fab in complex with BG505.T332N SOSIP and CH848 10.17 DS.SOSIP_DT; 3.5 Å and 3.7 Å C3 symmetry, respectively. **(B)** The V1 loop (red) adopts a similar orientation as observed for the complex between DH270.6 (colored in wheat) and CH848 Env contrasting with the orientation of the V1 loop observed in the complex with DH270-UCA (grey). **(C-D)** Atomic visualization of Env N332 glycan and GDIR interactions; (C) DH1030.1 HCDR3 loop makes extensive contacts with N332 glycan, and (D) DH1030.1 HCDR2 G56R mutation contact the Env V3 GDIR motif. **(E)** PGT121 (dark green, PDB:5CEZ), 1010-74 (wheat, PDB:5T3Z) and DH270.6 (dark blue) (shown as surfaces) bnAbs make extensive contacts with N332 glycan (shown as sticks) as seen for DH1030.1. **(F)** Differences in angles of DH270.6 and DH1030.1 bound to CH848 Env. Spheres: the center of mass (CM) of CH848 gp120 (red), heavy chains of DH1030.1 (magenta) and DH270.6 (purple), CM of the heavy chains of DH1030.1 (light green) and DH270.6 (dark green) and the light chains of DH1030.1 (yellow) and DH270.6 (orange). Cryo-EM structures PDB IDs, 6UM6 and 6UM5, were used as the model for DH270.6 and DH270-UCA, respectively.

When compared to DH270 V3-glycan bnAb isolated from human HIV-1 infection ^8, 15^, we found that DH1030 shared similar binding footprint (**Figure 3E**), consistent with the remarkable similarity in the evolution of a glycine to arginine mutation at an almost identical HCDR2 site in the human DH270 and macaque DH1030 V3-glycan bnAbs (**Figures 3F and S6A**). DH1030 mAbs bearing a revertant mutation at position 56 (R56G) in the HCDR2 of the heavy chain variable region (V_H_) had dramatic reduction or abrogation of HIV-1 neutralization against a panel of heterologous viruses (**Figure S6B**), indicating the emergence of V_H_ G56R as a key mutation associated with acquisition of DH1030 heterologous HIV-1 neutralization capacity as previously described for human DH270 V3-glycan bnAb where V_H_ G57R was associated with acquisition of neutralization breath ^8^. DH1030.1 and DH270.6 bound Env with overlapping footprints, although an ∼22° rotation of the Fab oriented the DH1030.1 light chain away from Env, resulting in a heavy chain dominated recognition mode of DH1030.1, in contrast to DH270.6 where both heavy and light chains make extensive contacts with Env (**Figure 3F**). Structural analyses of human V3-glycan bnAbs in complex with HIV-1 Envs (**Figure S6C**) indicated a role for suitably positioned arginines in Ab heavy and light chain variable regions in mediating contact with the ^324^GDIR_327_ motif in Env. This was supported by site directed mutagenesis studies for human as confirmed via site directed mutagenesis studies for human DH270 ^8^ and macaque DH1030 V3-glycan bnAbs (**Figure S6B**).

Consistent with the close contact observed between DH1030.1 and the N332 glycan, DH1030 mAbs demonstrated N332 glycan-dependent HIV-1 neutralization (**Figures 4 and S6B**). DH1030 mAbs neutralized up to 29% of a panel of 208 HIV-1 isolates, and up to 42% of 146 heterologous HIV-1 isolates bearing the N332 glycan in the 208-virus panel (**Figure 4A**). Like DH270, DH1030 mAbs did not neutralize any of the 46 heterologous HIV-1 strain among the 208-virus panel that had N334 instead of N332 or did not encode either residue (**Figure 4B**). Testing the statistical similarity of neutralization profiles between DH1030.5 and other human V3-glycan bnAbs, we found that DH1030.5 showed statistically significant correlations in IC50 titers against heterologous pseudoviruses, and significant overlap between sensitive and resistant viruses shared across bnAbs. While comparison to each human V3-glycan bnAb was significant, the highest similarity was found with DH270.1 with Pearson r2 = 0.22 between IC50 titers (p=0.0002) and 84% overlap in mutually sensitive or mutually resistant (Fisher’s exact test, p = 3.1x10^-25^). Thus, these data demonstrate that DH1030 Abs had characteristics of V3-glycan bnAbs with remarkable similarity to human V3-glycan bnAbs, especially DH270.

**Figure 4.**
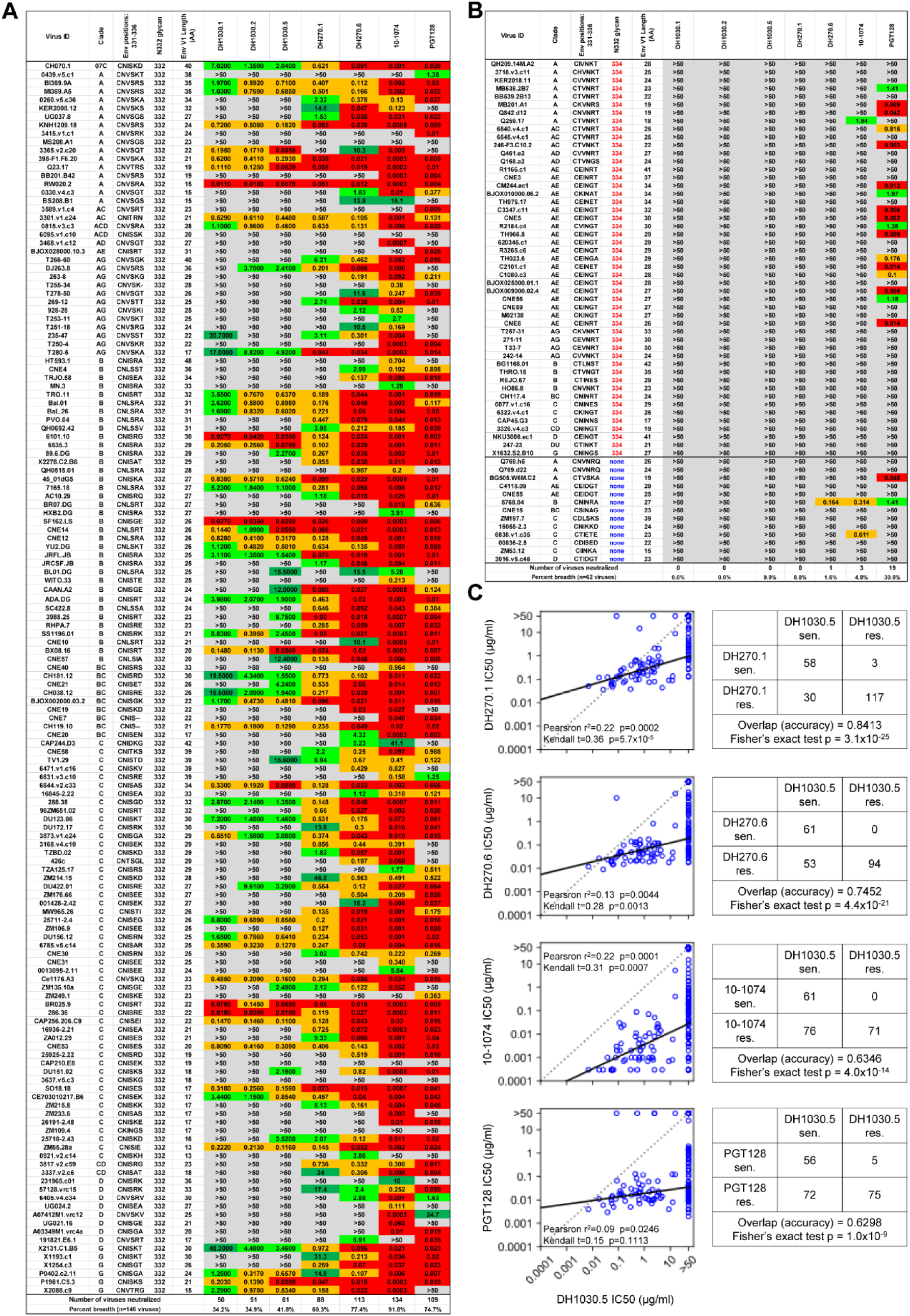
Neutralization profile of V3-glycan bnAbs, including DH1030. Neutralization titers of representative DH1030 Abs and known human V3-glycan bnAbs against a panel of 208 heterologous HIV-1 strains that encoded N332-glycan **(A)**, encoded N334-glycan instead of N332 or encoded neither N332 nor N334 **(B**). Neutralization was measured in TZM-bl cells and reported in IC50 (µg/ml). Neutralization titer is color coded for potency. (**C**) The left panels in each row show IC50 titers for DH1030.5 and other V3-glycan bnAbs for heterologous viruses from the 208-virus panel. Correlation coefficients and p-values from Pearson and Kendall tau rank tests, and the linear regression line were calculated using data points with IC50 < 50µg/ml for both bnAbs. The contingency tables on the right show the overlap in the sensitive/resistant viruses between DH1030.5 and other V3 glycan bnAbs. The overlap (accuracy) in number of mutually sensitive (sen) or resistant (res) viruses and the Fisher’s exact test p-values are shown below each contingency table.

### Genetic signatures of DH1030 bnAb lineage maturation

To fully evaluate the B cell repertoire of DH1030 Abs in peripheral blood at week 105 post-SHIV infection in RM08N021, we used an antigen-specific high-throughput approach termed LIBRA-seq ^51^ to identify 10 new clonally-related DH1030 Abs (D1030.7-DH1030.16) and an antigen-unbiased high-throughput single cell immune profiling assay (10x Genomics, Pleasanton, CA) ^47^ to identify 46 additional clonally-related DH1030 Abs (DH1030.17-DH1030.62). Phylogenetic analysis of the first 62 DH1030 Abs isolated (traditional sorting, n=6; LIBRA-seq, n=10; 10x Genomics single cell immune profiling, n=46) showed that they clustered into three clades; majority of clade 1 Abs had V_H_ G56 signature, whereas all clade 2 Abs had V_H_ G56 residue, and all clade 3 Abs had V_H_ R56 residue (**Figure S10A**). Like in the human DH270 bnAb lineage ^8^, V_H_ R56 was fixed in DH1030 Abs following the G56R mutation as seen in clade 3 of the DH1030 phylogram. We expressed representative mAbs from all three clades to determine the functional profiles of DH1030 Abs encoding glycine, arginine or another residue at HCDR2 position 56. Of six mAbs from clade 1, three (DH1030.22, DH1030.20 and DH1030.27_53) neutralized the autologous BG505.T332N or heterologous CH848 viruses that were tested, but none of them neutralized heterologous tier 2 HIV-1 strains from the global panel of reference strains ^52^ (**Figure S10B**). Interestingly, the three clade 1 mAbs with limited neutralization capacity had V_H_ G56 or S56 residues, and one additional clade 1 mAb (DH1030.34_46) that had V_H_ R56 was non-neutralizing (**Figure S10A-S10B**). Additionally, none of three mAbs from clade 2 that encoded V_H_ G56 neutralized HIV-1 strains tested (**Figure S10B**). In contrast to results from clades 1 and 2, all nine mAbs tested from clade 3 with V_H_ R56 neutralized up to 6 of 9 heterologous tier-2 HIV-1 strains from the global panel of reference HIV-1 strains (**Figure S10B**). Interestingly, DH1030 nAbs from clades 1 and 3 potently neutralized a heterologous tier-2 HIV-1 strain (CH848.10.17_N133DN138T) ^15^ with PNGS deletions in Env V1 region (V1-glycan deletions) (**Figure S10B**), indicative of the sensitivity of DH1030 Abs to viruses with V1-glycan deletions. However, of the clade 1 and 3 nAbs, only the clade 3 nAbs were able to neutralize the CH848.10.17 strain with the V1-glycans restored ^8, 15^ (**Figure S10C**). These data demonstrated that the clade 3 nAbs had matured to accommodate V1-glycans on Env, which is characteristic of development towards bnAb status, and were not strain-specific nAbs. All the clade 1 and 3 nAbs demonstrated N332-dependence via ELISA (**Figure S10D-S10E**). Thus, even within a bnAb B cell lineage like DH1030, only a subset of B cells acquired the appropriate mutations that conferred neutralization breadth towards development of bnAb status.

To identify additional mutations associated with acquisition of HIV-1 neutralization breadth, we analyzed heavy chain sequences for DH1030 Abs along the path to clade 3 bnAbs and found a serine-to-proline mutation at position 114 (S114P) in the HCDR3 that was present only in clade 3 bnAbs, except DH1030.4 and DH1030.5 (**Figure S11A**). Reverting V_H_ P114S into DH1030.1 bnAb had minimal reduction in binding but substantial reduction in neutralization breadth (**Figures S11B-S11C**). Computational modeling of V_H_ S114P on DH1030.1-Env complex suggested that V_H_ P114 residue in the HCDR3 introduced a bend in the heavy chain to bring neighboring aspartic acid residue at position 113 in the HCDR3 (D113) within H-bonding distance with the N332 glycan (**Figure S11D**). Thus, S114P mutation in the HCDR3 may serve as a compensatory mutation to G56R in the HCDR2 for the development of some DH1030 bnAbs.

### Maturation pathways of DH1030 V3-glycan bnAb lineage

Given the maturation of HIV-1 Env-reactive bnAbs via germinal center (GC) reactions in immune tissues ^26^, we next sought to interrogate DH1030 GC B cells in LN and spleen. Here, we modified the Barcode Enabled Antigen Mapping (BEAM)-Ab assay ^53^ to study both antigen-reactive and non-antigen-reactive B cells in different B compartments. Here, barcode-labeled antigens (see methods) were used to identify antigen-reactive B cells from PBMC, LN, or spleen although we sorted different B cell subsets in these samples based on cell surface phenotypes with designed Ab panels capable of detecting naïve, germinal center, and memory B cell populations (**Figure S12**). Post-sort analysis with barcode-labeled antigens allowed deconvolution of the predicted antigen-reactive B cells among the compartments studied as previously described for BEAM-Ab assay ^53^. Thus, we established a sequencing-based assay to assess phenotyping and BCR-antigen reactivity (PheBAR-seq) of single B cells in blood and secondary lymphoid tissues.

Using our PheBAR-seq assay, we studied eight B cell subsets from PBMCs, LNs and spleen at week 105 post-SHIV infection, and recovered 270,746 cells with paired immunoglobulin heavy (IgH) and light (IgL) chain genes, of which were 141 new clonally-related DH1030 members (**Table S5; Figure 5A**). The estimated frequency of DH1030 lineage B cells ranged from 300-600 per million B cells in peripheral blood and LN/spleen tissues (**Table S6**). DH1030 Abs were detected predominantly in the memory subsets of PBMCs, LN and spleen as well as the GC subset in the LN and spleen (**Figure 5A**). Following PheBAR-seq, we isolated a new total of 203 DH1030 Abs that used macaque V_H_3-al encoding 17-aa long HCDR3 and paired with V_K_2-g gene (**Table S5**), and phylogenetic analyses of these Abs indicated that they clustered into different clades based on genetic signatures, in particular the amino acid residue at VH_56_ position, rather than the B cell compartments (**Figure S13**).

**Figure 5.**
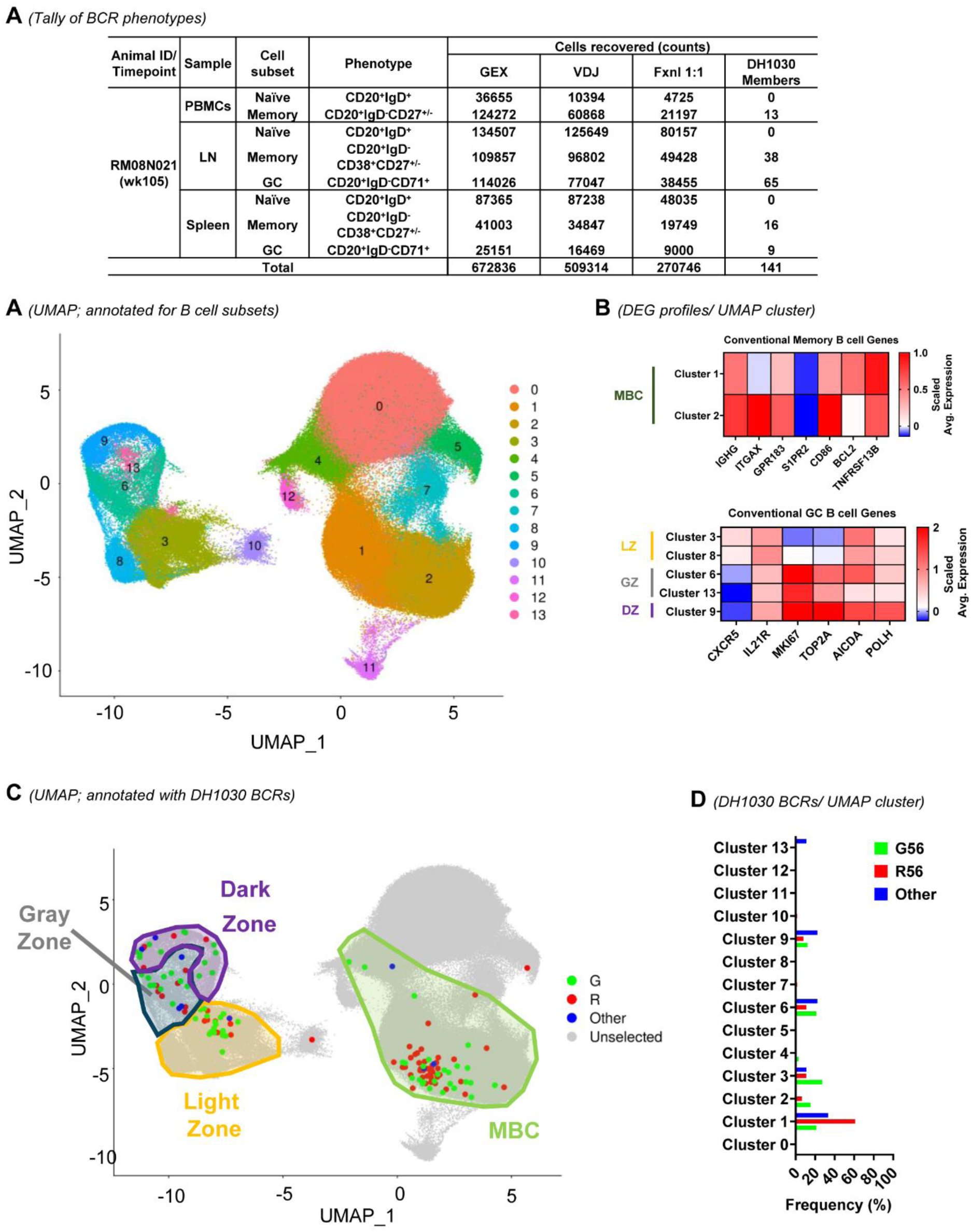
Transcriptional profiling of clonally-related DH1030 B cells. **(A)** Table listing the cell surface markers for B cell subsets sorted via flow cytometry, and number of cells recovered for analyses of BCRs with 1:1 functional heavy-light chain pairing and gene expression (GEX) profiling. The number of clonally-related DH1030 BCRs based on similar VD_H_J_H_ rearrangements is also shown; see table S5 for predicted antigen-reactivity of DH1030 B cells. **(B)** Integrated UMAP of B cells with BCRs identified in 14 unique clusters. UMAP reflects integrated LN, spleen, and PBMC cells that were sorted based on surface phenotype markers. **(C)** GEX profiling by scaled average expression for distinct lineages that make up the UMAP clusters in panel B. Clusters indicative of light, gray and dark zones in the GC, and memory B cells (MBC), are shown in the shaded regions on the UMAP. Each lineage was defined by canonical memory and germinal center B cell genes. **(D)** Integrated UMAP of B cells showing DH1030 lineage members with different genotypes based on amino acid residue at HCDR2 position 56; G (green), R (red), or Other (blue) amino acid. **(E)** Frequency of DH1030 lineage members in each UMAP cluster indicated in panel B.

The gene expression (GEX) profiling of the 270,746 B cells with paired IgH+IgL genes studied via PheBAR-seq was in agreement with our sort strategy for naïve, GC, or memory phenotypes from the eight cell subsets studied; B cells predominantly clustered based on phenotypes for GC, memory and naïve cell subsets (**Figure S14A-S14B**). Moreover, genes conventionally associated with GC, memory and naïve B cell subsets were enriched in the respective B cell subsets (**Figure S14B**). The total B cells across all three subsets clustered into 14 transcriptionally unique clusters using the Seurat R package (v5.3.0) for graph-based cell clustering, dimensionality reduction and data visualization ^54–56^ (**Figure 5B**). Clusters 1 and 2 demonstrated GEX profile consistent with memory B cells (**Figure 5C**). Cluster 1 was primarily isolated from LN memory cells and showed a GEX profile indicative of memory precursor B cells ^30–33^, confirmed by increased EBI2 (GPR183) expression required for light zone egress in tandem with downregulated S1PR2 which promotes GC retention. Additionally, expression of CD86, BCL2, and TNFRSF13B (TACI) are associated with recent egress of memory precursor B cells ^30, 31, 57, 58^. Cluster 2 was of memory PBMC origin and displayed a more mature memory B cell phenotype showing heightened expression of conventional switched memory B cell genes IgG, ITGAX (CD11c), and CD86 relative to precursor memory B cells identified in cluster 1 ^59^. Clusters 3, 8, 6, 13 and 9 demonstrated GEX profiles consistent with GC B cells; light zone (LZ; clusters 3 and 8), transitional or gray (clusters 6 and 13), and dark zone (DZ; cluster 9) GC B cell subsets (**Figure 5C**). GC B cell clusters 3 and 8 showed enriched expression of CXCR5, required for LZ trafficking, and IL21R, which is integral for B cell positive selection, in addition to low level expression of MKI67 and TOP2A indicative of G1 phase of the cell cycle consistent with LZ B cells (28-31, 42-43). Clusters 6 and 13 of GC origin downregulated CXCR5 while upregulating MKI67 and displayed a modest increase of TOP2A suggestive of S phase transitional B cells (28-31, 42-43). GC B cell cluster 9 also showed downregulation of CXCR5 with robust enrichment of MKI67, TOP2A, AICDA, and POLH indicating progression to G2/M phase and increased AICDA and POLH mediated somatic hypermutation (SHM) events typically found in DZ B cells (28-31, 42-43). Additionally, GEX of cell cycle genes revealed distinct subsets of GC B cells in G1, S, and G2/M phases (**Figure S14C-S14D**) in agreement with previous studies that established LZ B cells in the G1 phase, transitional cells in the S phase, and DZ cells in the G2/M phases ^60, 61^. Thus, our analysis was able to clearly define light and dark zone GC B cell compartments.

Next, we interrogated the B cell subsets studied via PheBAR-seq in search of DH1030 clonal lineage members. We found DH1030 clonal lineage members primarily in the GC and memory B cell subsets (**Figure 5D**). However, the highest frequency of DH1030 V_H_ R56-contaning B cells was present in the memory B cells of LNs (cluster 1) whereas G56-containing lineage members clustered in various compartments of LN and spleen GCs (clusters 3, 6, and 9) (**Figures 5D-5E**). DH1030 Abs with V_H_ G56 residue seemed to be predominantly retained in the GC; 61.54% of LN-derived GC DH1030 B cells having V_H_ G56 residue and 23.53% of LN-derived memory DH1030 B cells having V_H_ G56 residue (**Figure 6A**). That 30.77% of LN-derived GC DH1030 B cells had V_H_ G56R improbable mutation and 70.59% of LN-derived memory DH1030 B cells had V_H_ G56R mutation suggested that DH1030 V_H_ G56R mature lineage members were able to progress from GC to a memory phenotype (**Figure 6A**).

**Figure 6:**
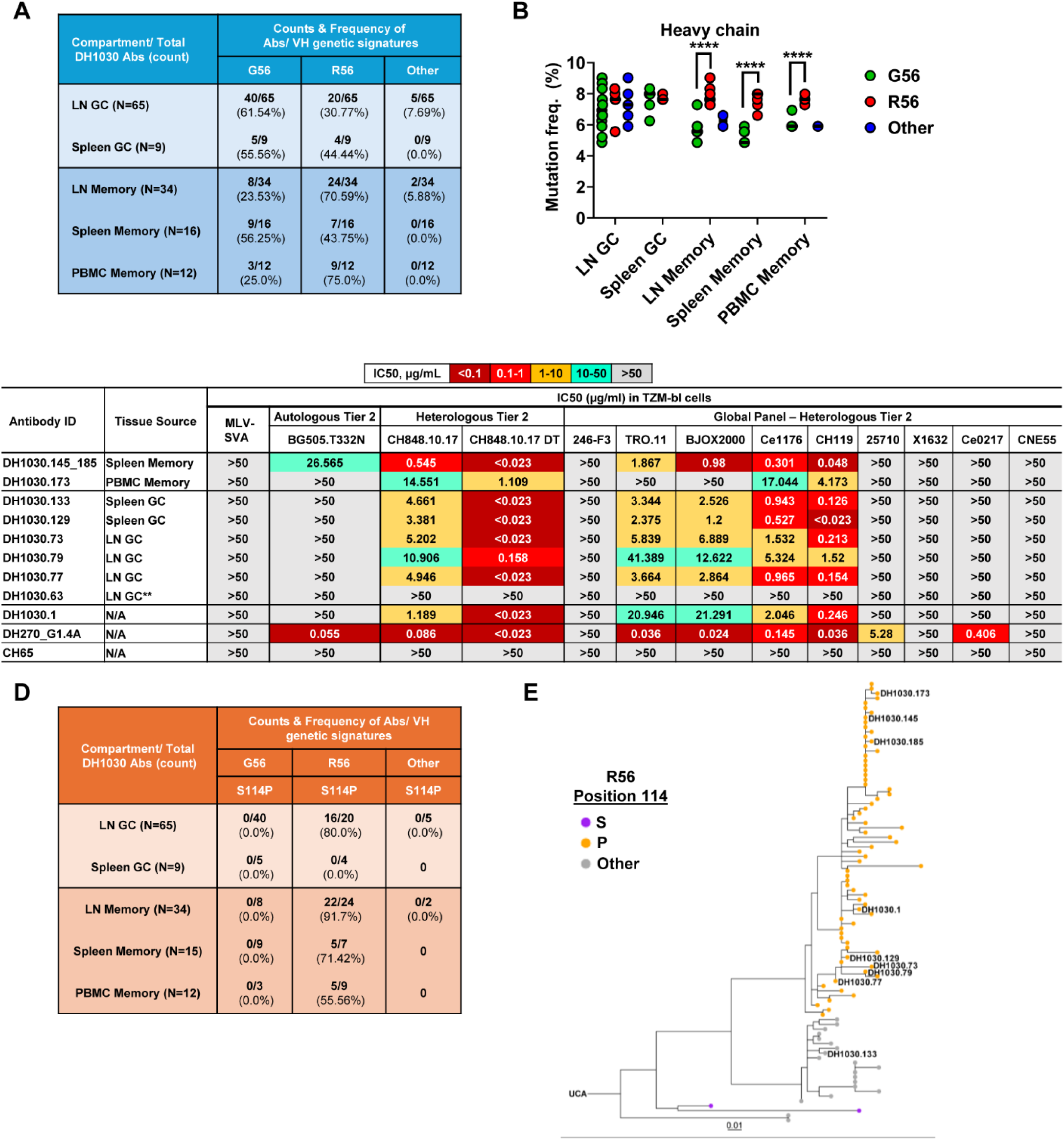
Tissue-specific maturation of DH1030 clonally-related B cells. **(A)** A table listing tissue origin and phenotype of DH1030 B cells bearing glycine (G), arginine (R) or other residues at HCDR2 position 56 that were studied via PheBar-seq. **(B)** Heavy chain somatic hypermutation (SHM) frequency in DH1030 B cells from different compartments; ****p-value <0.0001 (Two-way ANOVA, GraphPad Prism v10.1.2). **(C)** Neutralization profile of representative DH1030 Abs against autologous and heterologous HIV-1 strains. Neutralization titers reported as IC50 in µg/ml. All mAbs tested had arginine at position 56 in the HCDR2, except DH1030.63* that encoded S56 residue. **(D)** Table detailing tissue origin and phenotype of DH1030 B cells encoding S114P mutation in the HCDR3 in addition to either a glycine, arginine or other residues at HCDR2 position 56. Shown are the numbers and frequencies (%) for DH1030 BCRs with these phenotypes in different compartments. **(E)** Phylogenetic tree of clonally-related DH1030 VH sequences harboring G56R mutation and whether they evolve with either serine (purple), proline (yellow), or other (gray) residues at position 114 in the HCDR3. DH1030 Abs indicated on the phylogram were generated as mAbs and tested for neutralization in panel C, except DH1030.63 encoding S56 instead of R56 signature.

However, the enhanced progression of DH1030 V_H_ R56 lineage members from GC to memory phenotype coincided with increased SHM frequencies observed in DH1030 V_H_ R56 lineage members having a memory phenotype compared to DH1030 V_H_ G56 lineage members (**Figure 6B**). We next expressed and functionally characterized representative DH1030 mAbs that were detected in memory B cells present in PBMCs (DH1030.173), LNs (DH1030.145) and spleen (DH1030.185); and GC B cells present in LNs (DH1030.73, DH1030.77 and DH1030.79) and spleen (DH1030.129 and DH1030.133) (**Figure 6C**). The representative mAbs were among the DH1030 Abs containing the V_H_ G56R mutation with the highest prediction BEAM scores for antigen-reactivity and SHM frequencies in the memory and GC compartments of blood, spleen and LN. LN GC-derived DH1030.63 which encoded V_H_ S56 residue, a mutation from G56 in the germline gene that is unrelated to R56, was tested for comparison. We found that DH1030 mAbs with V_H_ R56 residue from different B cell compartments had similar potency and breadth of HIV-1 strains tested whereas DH1030.63 did not neutralize any strains tested (**Figure 6C**), thus reiterating the significance of the G56R improbable mutation for maturation of DH1030 Abs.

As shown in **Figure S11**, S114P mutation in the HCDR3 of DH1030 bnAbs was an additional mutation to HCDR2 G56R that was associated with maturation for neutralization capacity. Interestingly, in the B cell subsets studied via PheBAR-seq, we found that only DH1030 V_H_ R56-containing B cells encoded V_H_ P114 residue and these B cells were found only in LN-derived GC (**Figures 6D**). Moreover, DH1030 bnAbs encoding V_H_ R56 evolved with V_H_ P114 following the acquisition of this mutation associated with neutralization breadth (**Figures 6E and S11C**).

Taken together, these data demonstrated that DH1030 V_H_ G56R improbable mutation appears to be a pivotal catalyst for DH1030 lineage transition into the memory phenotype, and compensatory mutations including V_H_ S114P further promote maximal neutralization potency and breadth. These data also raise the hypothesis that compensatory mutations to V_H_ G56R play key roles in increased affinity to support bnAb lineage maturation—a focus of an independent investigation. Overall, our findings implicate a GC-driven maturation pathway in LNs wherein DH1030 key improbable mutations acquired during GC maturation established bnAb epitope recognition and neutralization profile necessary for transition into memory phenotype. In addition, these findings provide insights into the cellular context for the importance of improbable mutations during bnAb lineage maturation.

### DH1030 bnAb lineage induction

#### DH1030 lineage evolution

To interrogate the kinetics of DH1030 lineage development, we used Illumina next generation deep-sequencing (NGS) to trace clonally-related DH1030 BCR sequences with similar VD_H_J_H_ rearrangements in the heavy chain and VJ_K_ rearrangement in the light chain. We analyzed over 24 million BCR sequences derived from PBMC samples from weeks 12-96 post-SHIV infection and found clonal members in samples from weeks 80, 88 and 96 but not in earlier samples taken at weeks 12, 24, 64 or 72 (**Figure S15**). From these data, we inferred the sequences of the UCA (DH1030.UCA.v2) with high confidence using the SONAR pipeline ^41, 62^ (**Figures S15D-S15E**). To achieve this, we created RM 08N021-specific germline references for the heavy and light chain variable regions using IgDiscover ^63^ and identified animal-specific VD_H_J_H_ and VJ_K_ alleles for DH1030 bnAb UCA. As expected, SHM increased over time and DH1030 lineage sequences from week 80 post-SHIV infection showed SHM from the germline VD_H_J_H_ allele between 0.5% and 2.1% (mean = 1.1%). Thus, these data suggested that the DH1030 germline precursor was primed between week 72 and 80 post-SHIV infection. This timing coincided closely with the first detection of mutations in the ^324^GDIR_327_ epitope at residue 325 (**Figures 2A-2B**). Surprisingly, this estimation for timing of bnAb lineage priming conflicted with the first appearance of neutralization breadth in plasma, which was evident by week 40 post-SHIV infection. We reconciled this discrepancy by showing that plasma breadth at weeks 40-64 did not map to the V3-glycan epitope signature sites (^325^GDIR_328_ and N332 or N301) whereas later plasma samples from weeks 88-104 did (**Figures 1C and Table S1**). We could not identify the epitope target of this early bNAb response detected in week 40-64 samples. Altogether, our findings indicated that the DH1030 lineage was primed and expanded during late stage disease beginning between weeks 72-80 post-SHIV infection, which is similar to the late priming event of the human DH270 V3-glycan bnAb lineage ^8^.

#### Viral Envs associated with DH1030 lineage induction.

Next, we functionally characterized recombinant DH1030.UCA.v2 mAb to gain insights into the BG505 SHIV Env variant responsible for inducing the DH1030 lineage. Having narrowed the timing of DH1030 bnAb lineage priming to week 72-80 post SHIV infection, we examined closely SGS sequences near this time to identify candidate priming immunogens (**Figure 2A**). We were mindful of the recent demonstration that V1 deletions in the SHIV.BG505.5MUT Env led to rapid and consistent induction of V3-glycan bnAbs in macaques ^4^. **Figure 7A** shows an alignment of the V1 region of Env sequences cloned from SGS sequences from weeks 64 and 72 post-SHIV infection. Interestingly, one of these sequences (w072.25) contained a 4-aa deletion in an identical position as the del4 and del8 deletions as reported by Skelly et al. to be responsible for V3-glycan bnAb induction in SHIV.BG505.5MUT-infected monkeys ^4^. When assayed for neutralization sensitivity to the DH1030 UCA.v2 mAb, this w072.25 Env-pseudotyped virus was uniquely sensitive compared with the other forty-three RM08N021 Envs tested (**Figures 7A-7B**). Of note, this 4-aa deletion in V1 was also found in RM 10N011 that also developed V3-glycan bnAbs, and in 3 other macaques (**Figure S4C**). In a similar fashion, the human DH270 V3-glycan bnAb UCA did not neutralize the CH848TF virus but potently neutralized a variant with shorter V1 length (CH848.10.17) and lacking glycans at V1 residues 133 and 138 (CH848.10.17.DT) ^8, 15^. These data suggested that Envs with shorter V1 sequences lacking glycans in the Env V1 region may promote priming and expansion of V3-glycan bnAb B cell lineages in macaques and humans.

**Figure 7:**
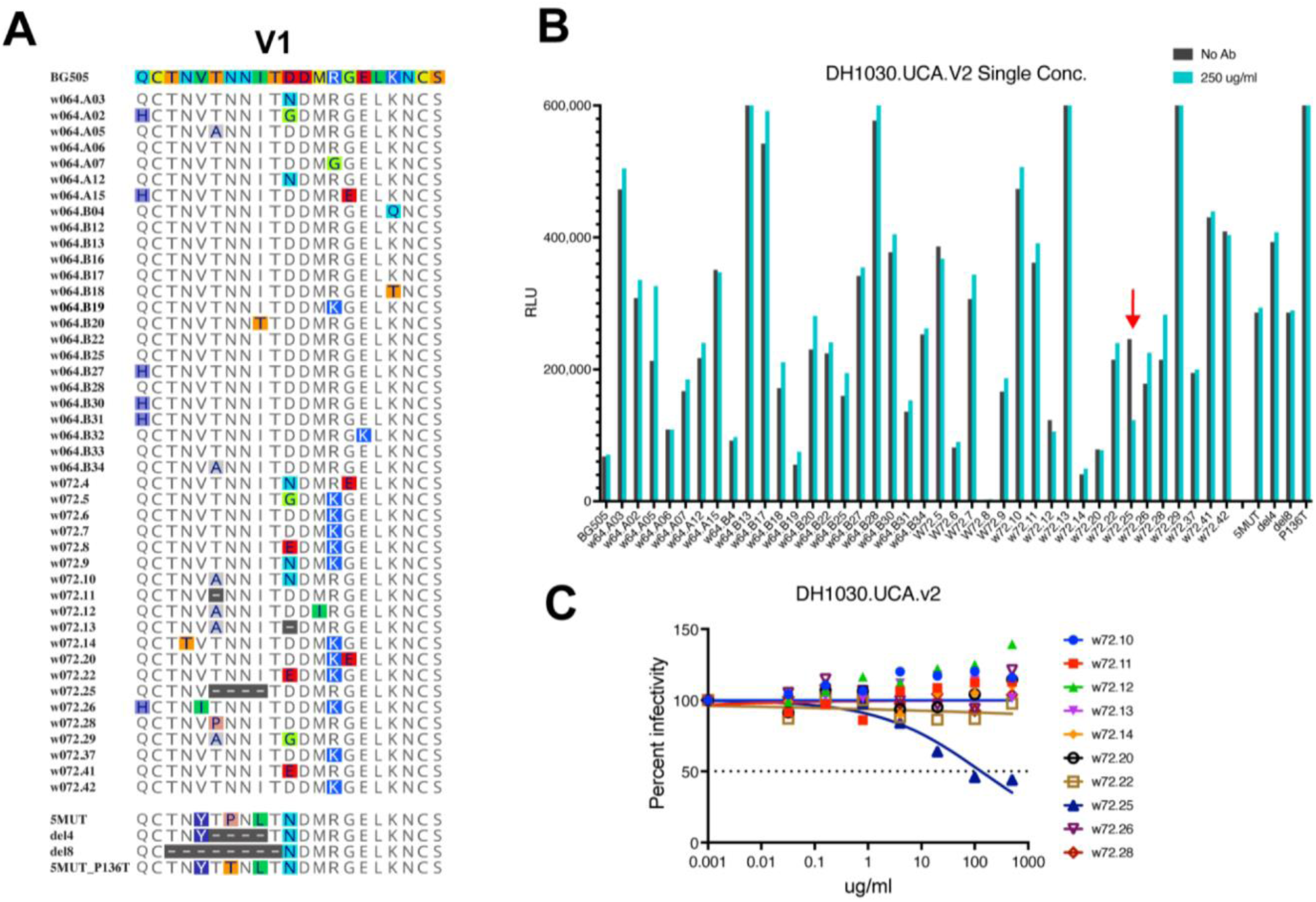
Neutralization sensitivity of DH1030 UCA to autologous Envs at weeks 64 and 72 of RM 08N021. **(A)** V1 sequence evolution in monkey 08N021 at weeks 64 and 72. **(B)** A single concentration of 250 μg/ml was used to screen positive binding Envs by neutralization assay. **(C)** Neutralization assays were performed by serial dilutions to obtain the IC50 of DH1030.UCA against w72 envs.

## Discussion

We infected a large number of RMs (N=18) with a pathogenic SHIV bearing the TF Env from a prototype BG505 strain used widely in HIV-1 vaccine research to understand the molecular basis for its early strain-specific immunogenicity and how the Env evolved to elicit bnAbs. Strikingly, we found consistent patterns of Env evolution and immune escape in all animals in the first year of infection, and then a distinct pattern of escape focused on ^324^GDIR_327_ and N332 in a subset of animals that developed V3 glycan bnAbs. By studying sequence evolution over the course of infection in all SHIV infected monkeys compared with the human subject BG505, we found remarkable convergence in Env evolution with the same or similar amino acid mutations identified under selection across the macaque hosts and the human.

Additionally, we isolated a large V3-glycan bnAb clonal lineage consisting of 203 members, termed the DH1030 lineage, from a representative SHIV BG505 infected RM 08N021 with V3-glycan targeted plasma breadth. Remarkably, DH1030 Abs elicited by subtype A BG505 macaque infection demonstrated striking genetic, functional and structural similarities with the human V3-glycan bnAb lineage DH270 that was elicited by a subtype C infection: a demonstration of convergent evolution with DH1030 being the only Ab found to date in both primates and non-human primates to mimic human DH270 at the genetic, functional and structural levels. Having 203 DH1030 B cell clonal lineage members facilitated extensive evaluation of the maturation pathway of DH1030 in circulating and tissue B cell compartments, and we now show that improbable mutations associated with maturation of macaque DH1030, as previously observed for human DH270, were predominantly found in memory compared to germinal center (GC) B cell subsets of LN and spleen. Finally, we demonstrated that an improbable mutation associated with maturation of DH1030 and DH270 V3-glycan bnAb lineages appeared to be a pivotal catalyst for lineage transition into the memory phenotype following a GC reaction. These findings implicate a trajectory for vaccines to replicate when selecting bnAb lineage members with improbable mutations that drive development towards bnAb status.

Our data highlight the value of SHIV-infected macaques as an outbred model system to explore conserved molecular pathways of bnAb development following infection and vaccination. Plasma V3-glycan bnAb induction in SHIV BG505 infected macaques was associated with the evolution of Env variants with deletion of PNGS in the V1 region of Env—a concept that is currently being tested in the clinic in different Env strains to elicit V3-glycan bnAbs ^64^. We previously postulated that V1 PNGS deleted Envs likely expanded the DH270 V3-glycan bnAb lineage in human HIV-1 infection ^8, 15^, and V1-glycan deleted Envs have been shown to be effective at isolating putative V3-glycan bnAb precursors in HIV-1 naïve humans ^19, 21, 23^. Enhanced neutralization of autologous and heterologous Envs with V1-glycan deletions by DH1030 mAbs, and neutralization sensitivity of DH1030.UCA.v2 mAb to an autologous Env bearing V1 glycan deletions, suggest that macaque DH1030 lineage was induced by a V1 glycan-deleted Env, consistent with recent observations by Skelly et al. in SHIV.BG505.5MUT macaque infection ^4^. Unique to the current study is the identification of an Env priming immunogen for a macaque DH1030 V3-glycan lineage that was isolated from SHIV BG505 infected RM 08N021; the Env immunogen had V1-glycan deletions in agreement with concepts postulated for eliciting V3-glcyan bnAbs. Altogether, these data highlight a common maturation pathway of human and macaque V3-glycan bnAb lineages that may be exploited for vaccine development.

V3-glycan bnAbs have been reported to use varied angles of approach to mediate Env recognition of the V3-glycan supersite ^5, 7, 8, 15, 65, 66^. Previously, Wang *et al.* isolated and characterized a macaque V3-glycan bnAb (Ab1485) that was elicited by SHIV AD8 infection, but the authors did not study the ontogeny and maturation events of this V3-glycan bnAb lineage ^66^. Here, we found that macaque and human V3-glycan bnAbs used different Ig gene pairs but encoded a common motif that facilitated Ab-Env recognition in a bnAb lineage. For example, an improbable mutation in the HCDR2 of both macaque (V_H_ G56R) and human (V_H_ G57R) V3-glycan bnAbs was associated with the acquisition of neutralization breadth, thus demonstrating remarkable similarity in bnAb B cell evolution in humans and TF Env SHIV-infected RMs ^12, 24^. Having found multiple clonally-related DH1030 members, we found that compensatory mutations to V_H_ G56R likely drove the maturation of the DH1030 lineage from precursor to bnAb status. Phylogenetic analyses of the DH1030 clone demonstrated that within the DH1030 V3-glycan bnAb lineage only a subset of the Abs acquired the appropriate mutations associated with acquisition of tier 2 heterologous HIV-1 neutralization activity; a characteristic of bnAbs. Furthermore, our transcriptomic analysis of DH1030 maturation revealed that acquisition of V_H_ G56R improbable mutation appeared to be key for both: 1) exit of DH1030 GC B cells into the memory population and 2) driving mutations that further enhance neutralization potency and breadth, for example, V_H_ S114P. These data imply that priming and boosting vaccine regimens to elicit V3-glycan bnAbs must select the appropriate mutations associated with acquisition of neutralization breadth in the BCRs of Abs at different stages of bnAb lineage development. Additionally, our data indicated that mutations associated with DH1030 V3-glycan bnAb lineage development were prevalent in BCRs of Abs predominantly found in the LN GC B cell subsets suggesting that DH1030 V3-glycan bnAb lineage matured via a GC-dependent pathway, consistent with HIV-1 vaccine regimens designed to engage GC reactions for bnAb development ^35, 67, 68^.

To investigate how two widely divergent HIV-1 Envs infecting two different primate species could share common developmental pathways, we analyzed the maturation pathway of the DH1030 macaque V3-glycan bnAb lineage by performing longitudinal BCR and transcriptome sequencing of B cell compartments in PBMCs and secondary lymphoid tissues, including LN, spleen, and BM. Here we established a high-throughput single cell approach to define the BCR and transcriptome sequences of total B cells in peripheral blood and tissues, including the antigen-specific B cells; we termed this assay PheBAR-seq. Here we demonstrated the use of innovative single cell technologies to study B cell immunology in RMs—a key resource for translational HIV research. In particular, we used our modified reagents and analytical pipeline using 10x Genomics technologies to identify macaque DH1030 BCRs in different peripheral blood and tissue B cell subsets that were defined based on gene expression profiling. Our methods have wide applicability in the field of infectious diseases that use non-human primates like RMs.

Previous studies have shown that macaque HIV-1 BG505 SOSIP trimer immunizations and SHIV BG505 infections predominantly generated autologous nAbs with strain-specific binding to glycan-hole epitopes ^69–72^. Consistent with those studies, we find that SHIV BG505 (+/-N332 glycan) infected macaques typically induce similar nAbs that select for Env evolutionary pathways that are remarkably recurrent across multiple macaques and the BG505 participant. Despite this propensity for nAbs targeting autologous epitopes, three RMs induced V3-glycan bnAbs after more than one year of infection. Our data support other findings that variants of the BG505 Env that evolve during SHIV infection can develop features that are favorable for the induction of V3-glycan bnAbs, particularly shortening and hypoglycosylation of V1 ^4^. These data raise the possibility that modified BG505 Env that mimics naturally evolved BG505 variants in infection might elicit HIV-1 Env V3-glycan targeted bnAb lineages in humans. This is consistent with recent observations that different versions of BG505 Env with V1 modifications showed promise in preclinical animal studies by inducing bnAbs ^4^ or bnAb precursors ^73^, and different forms of BG505 and other strains with V1 deletions are currently being evaluated in human clinical trials for the induction of V3 glycan bnAb lineages ^74, 75^. Thus, the significance of this paper is that there is a common maturation pathway of human and macaque V3-glycan bnAb lineages that may be exploited for vaccine development. Moreover, the maturation pathway for DH1030 provides a benchmark for the mechanisms of V3-glycan bnAb lineage induction in the human clinical trial that is studying DH270-like V3-glcan bnAb lineage induction (NCT05903339).

## Materials and Methods

### Study Design

All RMs were maintained in accordance with the Association for Assessment and Accreditation of Laboratory Animals. Research was conducted in compliance with the Animal Welfare Act and other federal statutes and regulations relating to animals and experiments involving animals and adheres to principles stated in the Guide for the Care and Use of Laboratory Animals, NRC Publication, 2011 edition. SHIV infection of Indian RMs were performed at (Bioqual Inc., Rockville, MD) and previously described ^12^. HIV-1 transmitted-founder (TF) Envs were derived from viruses that established infection in humans, thus TF Env-bearing SHIVs ^24^ provided a novel model for studying HIV-1 infection as described ^12^. Blood and plasma samples were collected for binding and neutralization assays. HIV/SHIV bearing BG505.T332N Env was referred to as an autologous virus strain.

### Viral RNA sequencing

We employed a previously described single-genome sequencing approach to amplify a total of 3,086 SHIV 3′ half-genomes from serial plasma samples of 17 animals ^43^. We didn’t sequence RM6447 because it died at wk9 post infection. In brief, viral RNA was extracted from plasma using the QIAamp Viral RNA Extraction Kit (Qiagen), targeting approximately 20,000 viral RNA copies. Reverse transcription was performed with SuperScript III Reverse Transcriptase (Invitrogen). The resulting cDNA was then subjected to endpoint dilution, followed by nested PCR amplification using previously reported primers and conditions ^24^. Single-genome template PCR products were sequenced via the Illumina MiSeq platform. Sequence alignment and analysis were conducted using Geneious Prime software, and the sequences were visualized with the Pixel Plot (https://www.hiv.lanl.gov/content/sequence/pixel/pixel.html).

### B cell isolation strategies

#### Conventional flow cytometry sorting of B cells

Conventional flow sorting of antigen-specific B cells using SOSIP trimer antigens as baits was previously described ^76, 77^. Briefly, cryopreserved PBMCs were thawed and counted and cells labeled with optimized concentrations of fluorochrome-mAb conjugates and fluorophore-labeled SOSIP trimer antigens as B cell baits. Here, we sorted B cells that bound wild-type Env CH119 SOSIP.v8.2 as previously described ^78^ that was conjugated to Alexa Fluor 647 (AF647) but did not bind the mutant Env CH119 SOSIP.v8.2 N332T conjugated to VioBright 515 (VB515). The single cell sorted B cells were then studied for heavy and light chain gene amplification using PCR-based approaches as described ^47, 48, 76, 77^.

Our approach for flow cytometry sorting of antigen-unbiased plasmablast or plasma cells in cryopreserved cells of BMAs was previously described ^49^. Here, we sorted plasmablast or plasma cells based on the following gating hierarchy: CD14-CD16-CD3-CD20-HLA-DR+CD11c-CD123-CD80+IgG+/-. We amplified the heavy and light chain genes from individually sorted cells as previously described ^47, 48^.

#### Modified 10X Genomics single cell immune profiling assay for isolation of antigen-naïve B cells

B cell enrichment and high throughput single cell profiling was performed on peripheral blood cells using a modified single cell immune profiling assay as previously described to facilitate macaque VDJ sequencing ^47^. Briefly, B cells were isolated from PBMCs using negative enrichment (Stemcell Technologies) and loaded onto a 5’ v1.1 chip across all 8 lanes, targeting 10,000 cells per lane. Gene expression libraries (GEX) and VDJ libraries were generated as previously described and sequenced on a NovaSeq 6000 using read lengths R1: 26, R2: 91, I1:8, I2: 0 for GEX; and R1: 26, R2: 150, I1: 8, I2: 0 for VDJ. Read depths targeted from GEX were 20,000 reads/cell for the GEX libraries and 2,000 reads/cell for the VDJ libraries ^47^. For VDJ amplification in the single cell immune profiling assay, we used primers designed to target the constant regions of macaque immunoglobulin genes as described ^47^.

#### LIBRA-seq isolation of antigen-reactive B cells

Antigens used for single B cell isolation via flow cytometry were biotinylated and then conjugated to a fluorophore for antigen-specific B cell sorting. Here, we conjugated the biotinylated antigens possessing a barcode label to a streptavidin-fluorophore-DNA barcode (BioLegend TotalSeq-C). All antigens were conjugated to a PE fluorophore and also had a unique DNA barcode. We sorted all PE positive B cells, and used the unique DNA barcode for each antigen during analysis to identify candidate antigen-reactive B cells. Briefly, BG505.T332N SOSIP, CH848.10.17_DT (N133DN138T) SOSIP, and CH505TF SOSIP were conjugated to their unique TotalSeq C-PE-DNA barcode using the same ratios for conjugation as described above for flow sorting with antigens not linked to a DNA barcoded fluorophore. B cells were then negatively enriched as described above, incubated with the three barcoded antigens, and sorted into 20μl of PBS + 0.04% BSA. To maximize cell recovery, cells were not recounted before loading onto the 10X chip. Libraries were generated using the 10X 5’ VDJ and Feature Barcode kits from 10X genomics.

### Pipeline for PheBAR-seq

#### Production of BEAM-Ab Assemblies for B cell Sorting

To isolate antigen-reactive B cells, antigen assemblies were generated comprised of antigen and a BEAM Conjugate that contains 10X barcoded streptavidin molecule linked with a PE fluorescence marker. The BEAM Assemblies served as B cell baits or hooks to capture antigen-reactive B cells, and were produced according to the 10X Genomics Barcode Enabled Antigen Mapping (BEAM) Workbook for BEAM-Ab Assembly (Doc CG000597 Rev A). The biotinylated proteins used to make BEAM Conjugates were HIV-1 Env antigens: BG505.MUT11B SOSIP, CH848 10.17 SOSIP, and CH848 10.17 DT SOSIP and a non-relevant HIV antigen as Coronavirus N Protein N-Terminal Domain (NTD). Two BEAM Assemblies were produced for each HIV-reactive antigen using unique barcodes to improve reliability of B cell capture and increase confidence of subsequent BEAM scoring. One assembly contained no antigen and was prepared as a negative control for background subtraction (see BEAM scoring). Antibody-coated beads (negative bead control Influenza HA-reactive mAb CH65; positive control beads: HIV-1 Env reactive mAbs PGT151 and NTD-reactive mAb DH1051 ^79^ were used to confirm expected binding and fluorescent profiles of BEAM Assemblies by flow cytometry ^53^.

#### Cell staining and sorting

Cryopreserved PBMC, Spleen, and LN samples from RM08N021 were thawed at 37°C and slowly diluted in 1mL RPMI with 10% FBS (complete RPMI, cRPMI) supplemented with Benzonase at 25 U/mL. After 1 minute, the cell suspension was further diluted with 13mL cRPMI. Cells were washed via centrifugation for 5 minutes at 400rcf, then resuspended in cRPMI with 5 μM Chk2 inhibitor (Millipore Sigma; Catalog #220486) at 2X10^6^ cells/mL final concentration based on the frozen cell count. After cell resuspension, an aliquot (10 µL) was taken for cell count verification using the Countess 3 Automated Cell Counter (Thermo Fisher). Following centrifugation, the cells were resuspended in FACS buffer (1% BSA in 1X PBS) with 5μM Chk2 inhibitor and CD4 blocking antibody, Clone SK3 (Biolegend; Catalog #344602; 1:100 dilution) on ice for 30 minutes at a final concentration of 1X10^6^ cells/100µL. An aliquot of 0.4-0.5x10^6^ cells was set aside as a negative control without BEAM antigens to determine antigen-specific sort gating, but stained with the rest of the panel. The remaining cells were stained with surface antibodies for flow sorting and BEAM Assemblies for 30 minutes on ice. Cells were stained with the antibody staining panel in FACS buffer at a final concentration of 1X10^6^ cells/100µL. Following cell staining, cells were washed with 1X PBS, then resuspended in 1X PBS with 5 μM Chk2 inhibitor and Aqua vital dye (Thermo Fisher; Catalog #L34957; 1:1000) for 30 minutes on ice at a final concentration of 1X10^6^ cells/100µL. Cells were then washed with FACS buffer and, finally, resuspended in 100µL per 1X10^6^ cells in FACS buffer with 5 μM Chk2 inhibitor and kept on ice. IgD+ naïve, IgD-CD71+ GC, and IgD-CD71-CD38+CD27+ memory B cells were sorted from PBMC, Spleen, or LN tissues into 500µL cRPMI with 5 μM Chk2 inhibitor in a FACS tube using a 100 μm nozzle on a BD FACSymphony S6 (DHVI Flow Cytometry core). Sorted cells were kept on ice for no more than 1 hour and then centrifuged at 400 xg for 5 minutes. The supernatant was slowly aspirated and discarded while leaving behind ∼50-200µL of residual cells in buffer to target 400-2000 cells/μL as per 10X Genomics recommendations. The resuspended cells were then loaded into single cell GEMs using the Chromium Controller microfluidics chip and reverse transcription kit (10X Genomics).

#### Gene library prep and sequencing

From RM08N021, sorted B cells populations were used to generate VDJ, gene expression, and BEAM barcode libraries using Chromium Next GEM single Cell 5’ HT Reagent Kits v2 (10X Genomics; Catalog #PN-1000356) with Feature Barcode technology for BEAM per the manufacturer’s recommendations (10X Genomics, Document #CG000591 Rev A). One modification was the use of rhesus Ig constant region primers to generate the VDJ libraries that we established in an independent study ^47^. For single cell encapsulation, we input all the cells recovered as quantified above into 2-8 lanes of the 10X Genomics CHIP, thus allowing us to target 20,000 cells per lane. After preparation of the gene libraries, we performed library QC was performed using TapeStation 4200 with the high sensitivity D1000 or D5000 ScreenTape (Agilent Technologies, Catalog #5067-5582) and Qubit Flex fluorometer (Thermo Fisher Scientific). The qualified VDJ, gene expression and BEAM libraries were pooled at 1:10:1 ratio, and sequenced with Novaseq X Plus 10B 100 cycles kit (Illumina) to achieve the sequencing depth of ≥5,000 read pairs/cell for VDJ libraries, ≥50,000 read pairs/cell for 5’ gene expression libraries, and ≥5,000 read pairs/cell for BEAM libraries following the manufacturer’s protocol (Illumina, Document #200027171 v02). For Illumina NGS, we loaded concentration of libraries at 150pM and 1% Phix (Illumina, Catalog #FC-110-3001).

#### BEAM-Ab scoring prediction analysis

The BEAM-Ab antigen-library will be used to computationally generate a BEAM Score, using an automated Cell Ranger software (10X Genomics), which is a measure of antigen-reactivity to identify HIV-1 Env-reactive BCRs. The generation of BEAM Scores use the target antigen unique molecular index (UMI) barcodes and the control no-antigen barcodes; a positive score is an indication of a higher ratio of antigen binding relative to the control no-antigen conjugate, and is scaled per antigen on a range of 0-100 (lowest to highest Ag-reactivity). Two BEAM Assemblies were produced for each HIV-reactive antigen using unique barcodes to improve reliability of B cell capture and increase confidence of subsequent BEAM scoring. UMI barcodes counts for dual barcode antigens were summed to be used in BEAM scoring calculations. BEAM scoring uses beta distribution modeling that computes 100 times the probability that a BCR is binding to an antigen with 92.5% confidence, thus 100 is the maximal BEAM score for the best binding BCRs and as the BEAM score decreases so does the likelihood for BCR binding. DH1030 clonal lineage members isolated by BEAM-Ab were identified based on BCR sequence similarity to DH1030.1. Contigs assembled by Cell Ranger were re-annotated and analyzed using Cloanalyst (https://www.bu.edu/computationalimmunology/research/software/) with the default rhesus Ig library to determine B cell immunogenetics and clonality as previously described ^80^. Previous BEAM experiments have informed our working criteria for selection of BCRs from BEAM-Ab for Ab expression ^53^. Our criteria for IgG Abs are as follows: heavy/light chain UMIs ≥5, BEAM score >50 for HIV-1 antigens, BEAM Score <20 for non-relevant NTD protein. All isolated DH1030 Abs were IgG (**Table S5**). Using these criteria we selected representative DH1030 BCRs that also had maximal highest mutation frequencies in different B cell compartments mAb for expression and functional characterization.

We hypothesized that BEAM+ IgM B cells bound due to avidity and is a possible explanation for BEAM+ naïve B cells I different compartments (**Figure S12**). These potentially high-avidity binding naïve B cells lose the binding profile when expressed as a recombinant monoclonal antibody in an IgG backbone from our experiences with this assay. However, using the BEAM conjugate lacking an antigen as a negative control, we were able to subtract background or non-specific binding due to these avid IgM responses, thus our output for BEAM+ IgG B cells are predominantly high affinity binders. Thus, our studies focus mainly on IgG BEAM+ B cells for recombinant mAb expression and functions.

#### GEX analysis: Log_2_FC and scaled average expression

We used the default Seurat function ScaleData() for the scaled expression values which subtract the average expression and then divides by the standard deviation for each gene. The log_2_FC for differentially expressed genes is calculated using the FindMarkers() function in Seurat which is the log fold change of the average expression between the groupings studied.

### NGS and analysis of Ab genes

To infer the sequence of the DH1030 lineage UCA we obtained BCR sequences from PBMCs sampled at weeks 12, 24, 64, 72, 80, 88, and 96 post infection. PBMCs were washed with PBS and total RNA was extracted using RNAzol according to the manufacturer’s guidelines (Molecular Research Center, #RN190). We then performed cDNA synthesis, amplified heavy and light chains with nested PCRs, and generated NGS libraries for IgA/D/E/G/M isotypes using a human BCR sequencing kit (Takara, #634777) with the human primers replaced with proprietary rhesus primers generously provided by the manufacturer for pre-commercial testing. Libraries were sequenced on the Nextseq2000 using 2x300 cycle paired read P2 kit (Illumina). Raw reads were demultiplexed using bcl-convert (Illumina, version 4.3.13), paired and filtered for quality control with vsearch (v2.30.1, PMID: 27781170) and DH1030 lineage members were identified using the SONAR pipeline ^41, 62^. Briefly, the sequence of the DH1030 UCA was constructed by assigning VDJ germline genes from a custom RM08N021 germline reference database generated from naïve IgM+/IgD+ B cell sequencing as previously described ^41^. Then, non-templated regions of the UCA were inferred using the consensus of early lineage members with the fewest mutations from assigned germline genes as well as IgPhyML included with SONAR to generate a phylogenetic reconstruction of the lineage evolution.

The phylograms for DH1030 sequence analyses were initially generated using DH103UCA.2 that was inferred via Cloanalyst using the BCR sequences of the isolated DH1030 B cells from multiple assays. B cell clonality was determined based on similar heavy chain rearrangements and CDR3 length as described ^81^. For Abs that were inferred as being clonally-related by Cloanalyst, and had the same V_H_ sequence but different alleles, we manually inspected the V_H_ sequence for similar residues in the HCDR3, including the non-templated nucleotides. Immunogenetics information of rhesus antibody sequences were assigned by Cloanalyst using Cloanalyst’s default libraries of rhesus immunoglobulin genes (https://www.bu.edu/computationalimmunology/research/software/). Phylogenetic analysis was performed using the Dowser R package, which is part of the Immcantation framework. The paired heavy and light chain tree was built using the raxml method and default parameters. The ANARCI tool was used to number amino acid sequences using the Kabat numbering scheme.

#### High-resolution single cell VDJ, feature barcode, and transcriptome sequencing and analysis

RM VDJ libraries were generated following the manufacturers protocol (Chromium Next GEM Single Cell V(D)J protocol (v1.1, Rev E)) with three modifications as previously described, including primer designs to amplify the BCR sequences from RMs ^47^. Primers were designed using the previously published macaque Ig libraries ^82^ and Cloanalyst default rhesus Ig library (https://www.bu.edu/computationalimmunology/research/software/). Immune profiling was performed using the Cell Ranger Single Cell Software Suite (versions 4.0.0, 7.2.0). Samples were demultiplexed, assembled, filtered, and aligned to a custom RM VDJ reference with RM Ig genes from three reference libraries [the Cloanalyst default Ig library – see link above, IMGT, and Ramesh *et al.* ^82^] as previously described ^47^. A text file specifying the sequences of the RM inner enrichment primers was also used as described in the documentation for running customized libraries in Cell Ranger provided by 10X Genomics. Assembled contigs from Cell Ranger and their chain annotations were then used as input VDJ sequences into the Cloanalyst software package. Immunogenetics information of RM antibody sequences were assigned using the default RM Cloanalyst library and clonal partitioning was performed using a custom pipeline.RM GEX libraries were generated following the manufacturers protocol (Chromium Next GEM Single Cell GEX protocol (Chromium Next GEM Single Cell 5’ Reagent Kits v2 (CG000591, Rev B)). Transcriptomic analysis of macaque single B cells was performed using the Cell Ranger Single Cell Software Suite (versions 4.0.0, 7.2.0). Samples were demultiplexed, assembled, filtered, and aligned to the Mmul10 reference (accession NCBI:GCA_003339765.3) followed by UMI counting.

Gene count matrices generated by Cell Ranger were analyzed using the Seurat R package (v5.3.0) for graph-based cell clustering, dimensionality reduction and data visualization ^54, 55, 83^. Differentially expressed genes between cell clusters or groups were determined using Seurat by the Wilcoxon Rank Sum test ^84^. Graphs and plots were generated using the Seurat and ggplot2 R packages and Graphpad Prism version 8.

LIBRA-seq analysis was performed using the Cell Ranger Software Suite (version 4.0.0). Demultiplexed FASTQ files were generated with the mkfastq command and processed using the count command, along with a feature reference file specifying the barcodes of oligonucleotide-tagged antigens. The resulting feature barcode count matrix was then used to calculate LIBRA-seq scores as previously described (39). BEAM analysis was performed using a similar procedure in Cell Ranger (version 7.2.0) and the feature barcode count matrix was used to calculate BEAM scores as described above.

#### Expression of recombinant mAbs

For generation of large quantities of recombinant mAbs, commercially-obtained (GeneScript, Piscataway, NJ) plasmids with antibody heavy and light chain genes were used to transfect suspension Expi 293i cells using ExpiFectamine 293 transfection reagents (Life Technologies, GIBCO; Cat#A14524) as described ^47^. Purified recombinant mAbs were dialyzed against PBS, analyzed, and stored at 4°C. All recombinant mAbs were expressed from plasmids encoding a human or macaque IgG constant region, and were QC’ed in Western Blot for appropriate heavy and light chain protein expression.

#### Site-directed mutagenesis

In order to generate constructs of representative mAbs, site-directed mutagenesis was performed using the QuikChange Lightning Multi Site-Directed Mutagenesis Kit (Agilent) using manufacturer’s instructions. Primers containing the desired mutations were designed and synthesized. The mutagenic primers were used in a PCR reaction with the respective heavy and light chain constructs as plasmid DNA template. Following PCR amplification, the product was treated with DpnI to digest the parental DNA template. The resulting mutant plasmid DNA was then transformed into competent cells and colonies were screened for the presence of the desired mutations by sequencing and further used for antibody production.

#### Binding ELISA

Envelope binding to recombinant mAbs were tested in ELISA as previously described ^47^. Binding was assessed using recombinant HIV-1 Env stabilized chimeric SOSIP trimers. In general, biotinylated antigens were captured using streptavidin that was coated to Nunc-absorb (ThermoFisher) plates overnight at 4°C. Unbound proteins were washed away and the plates were blocked with goat serum-based (SuperBlock) blocking medua for 1 hour. Serial dilution of mAbs were added to the plate for 60 minutes. Binding Abs were detected with rhesus-specific HRP-labeled anti-IgG Fc Abs using 20μL per reaction with 1 hour incubation at room temperature. HRP detection was subsequently quantified with 3,3’,5,5’-tetramethylbezidine (TMB). ELISA binding levels were measured at an optical density of 450nm (OD450nm) and binding titers were analyzed as area-under-curve of the log-transformed concentrations (Log AUC).

#### Neutralization Assays

Monoclonal antibody neutralizing activity was assessed in TZM-bl cells as described ^85^. Env-pseudotyped HIV-1 bearing Envs used in SHIVs for infection were referred to as autologous viruses, whereas viruses bearing Envs from global panels of geographically diverse multi-clade strains were referred to as heterologous viruses. For neutralization assays, a mixture of CH01+CH31 bnAbs is used as a positive control for neutralization of all HIV-1 strains, and murine leukemia virus (MLV) was used as a negative retrovirus control. For neutralization assays, a positive for neutralizing antibody activity in a sample is based on the criterion of > 3X the observed background against the MLV negative control pseudovirus. The neutralization panel of 208 viruses was previously described ^8^. In **Figure 2E**, we demonstrated the dynamic changes of autologous responses in RM 08N021 and RM 10N011 plasma. RM08N021 w24 plasma was not tested because no autologous NAb response was detected (reciprocal ID50 <20). RM10N011 w88 and w96 plasma were tested for only the V1 and V3 mutants since the autologous NAb activities against the 241/289 and C3V4V5 glycan holes had been negative for at least 4 months (weeks 56 to 72).

### Structural analyses

#### Cryo-Electron Microscopy (cryo-EM)

Purified HIV-1 Env SOSIP was diluted to a final concentration of about 1 mg/mL in 2 mM Tris pH 8.0, 200 mM NaCl and 0.02% sodium azide, were mixed with 6-fold molar excess of DH1030.1 Fab and incubated for 2 hours at room temperature. 2.5 µL of protein was deposited on a Quantifoil 1.2/1.3 holey carbon grid that had been glow discharged for 30 seconds in a PELCO easiGlow Glow Discharge Cleaning System. After a 30 second incubation in > 95% humidity, excess protein was blotted away for 2.5 seconds before the grid was plunge frozen into liquid ethane using a Leica EM GP2 plunge freezer (Leica Microsystems). Cryo-EM data were collected on a FEI Titan Krios microscope (Thermo Fisher Scientific) operated at 300 kV. Data were acquired with a Gatan K3 detector operated in counting mode. Data processing was performed within cryoSPARC ^86^ including particle picking, multiple rounds of 2D classification, ab initio reconstruction, heterogeneous and homogeneous map refinements, particle subtraction, local refinement, and non-uniform map refinements. ChimeraX ^87^, Coot ^88^, and Phenix ^89^ were used for model-building and refinement.

#### X-ray crystallography

Crystals of DH1030.1 Fab were grown in 30% (v/v) MPD 100 mM Sodium acetate/Acetic acid pH 4.5 25% (w/v) PEG 1500 at 22°C in a sitting drop vapor diffusion setting using a drop ratio of 0.3 µL protein: 0.3 µL reservoir solution. Large UV-active plate shaped crystals were observed. A single crystal was cryopreserved directly from the drop. Diffraction data was collected at the Advanced Photon Source using sector 22ID beamline. The collected diffraction images were indexed, integrated, and scaled using HKL2000 ^90^. Initial phases were calculated by molecular replacement using Phenix.PHASER ^91, 92^ and the Fab from PDB 6PDU (Vaccine-elicited NHP FP-targeting antibody 13N024-a.01 in complex with HIV fusion peptide (residue 512-519)) as a search model. Iterative rounds of manual model building using Coot ^88^ and automatic refinement in PHENIX ^89, 93^ were performed. Data collection and refinement statistics are summarized in **Table S2**.

ChimeraX, ePISA (Protein interfaces, surfaces and assemblies service at the European Bioinformatics Institute; http://www.ebi.ac.uk/pdbe/prot_int/pistart.html) 94, and PyMOL (The PyMOL Molecular Graphics System, Version 3.0 Schrödinger, LLC) were used for structural analysis and visualization.

#### Negative stain electron microscopy (NSEM)

10μg of BG505gp140SOSIP.T332N was incubated with 36μg of DH1030.27_53 Fab in HBSO4-150 buffer (90.7μL for a total of 100μL complex-forming solution) overnight at 4°C (15:1 molar ratio of Fab to SOSIP) (data not shown). The complex was then cross-linked using 8mM Glutaraldehyde-HBSO4-150 and then concentrated using 2 mL Amicon 100kDa MWCO filter tubes (spun at 4k rpm for 10 minutes).

This resulted in a nominal SOSIP concentration of 5.44 mg/mL. Two techniques were used for staining. For the first one, a dilution buffer of 0.02% RR-HBS was used to dilute the complex to a SOSIP concentration of 0.136 mg/mL. After 2-5 minutes, the solution was added to the staining grid for 8-10 seconds, then rinsed with 2 drops of 1/20x HBS, then stained with 2% UF. For the second technique, the Dilution buffer was prepared using HBSO4-150 to dilute the complex to a SOSIP concentration of 0.272 mg/mL. The solution was immediately applied to the grid for 10-12 seconds and then stained with 2% UF. For both techniques, the stain lasted for 1 minute. For data analysis, three datasets were initially run separately for particle picking, 2D classification, and selection.

### Env evolution analyses

Env evolutionary analyses were performed as previously ^12^. Briefly, LASSIE ^44^ was used to identify amino acids and glycan mutations under selection using the threshold of loss of >80% of TF sequence at any time point in each host. Glycan shield prediction was performed using the Glycan Shield Mapping webtool on the Los Alamos HIV Databases (https://www.hiv.lanl.gov/content/sequence/GLYSHIELDMAP/glyshieldmap.html) ^46^ using glycans at >50% frequency in Env sequences at each time point from each host. Recurrent hypervariable loop insertion/deletion (indel) analyses were performed by analyzing Env sequences from the first year from each host (after this time point typically indels and mutations on top of indels can confound identification of repeated patterns) and by manual alignment and inspection of the hypervariable loop sequences. All sites under selection in either of the 3 hosts (human BG505, and RMs 08N021 and 10N011) are shown in **Figure 2B**, where only representative time points are shown for RM08N021 and RM10N011; the full data are shown in **Figures S2-S3**.

### Reagent Authentication

Expi293F and TZM-bl cell lines were provided with a certificate of analysis from their sources. Cell identity is verified by morphology or fluorescent markers expressed. Positive control abs were used in the TZM-bl neutralization assay to distinguish the cell lines and for comparison between lots when new vials of cells are cultured.

### Statistical Analysis

Linear regression as implemented in SciPy (www.scipy.org) was used to compare Log10 transformed IC50 titers for DH1030.5 to other V3-glycan bnAbs using only those viruses that were neutralized by both bnAbs being compared (**Figure 4**). Fisher’s exact test was used to assess the statistical significance of the overlap between sensitive and resistant viruses between DH1030.5 and other V3-glycan bnAbs. For comparisons of heavy chain SHM frequencies in DH1030 B cells from different compartments, we performed a two-way ANOVA test using GraphPad Prism v10.1.2 (**Figure 5**).

## ACKNOWLEDGEMENTS

We thank the DHVI Program Management and Finance teams for their support. The following individuals provided technical assistance: Sommer Holmes, Roman Fenner, Erik Svenson, Esther Lee, Rebecca M. Williams, Mary Tess Overton, Savanna Toure, Wenge Ding, Juliette Rando, Neha Chohan, John Carey and Yu Ding. We thank Garnett Kelsoe for his intellectual contributions via discussions on B cell biology. We thank M. Louder, R. Carroll, N. C. Moore, C Lin, K. McKee, and A.B. McDermott for their assistance with neutralization assessments on the 208-strain panel, and J. Baalwa, D. Ellenberger, F. Gao, B. Hahn, K. Hong, J. Kim, F. McCutchan, D. Montefiori, L. Morris, E. Sanders-Buell, G. Shaw, R. Swanstrom, M. Thomson,S. Tovanabutra, C. Williamson, and L. Zhang for contributing the HIV-1 envelope plasmids used in this neutralization panel. We thank the DHVI Viral Sequencing Analysis and Duke University Genomic and Computational Biology Cores for BCR sequencing. Cryo-EM data were collected at the Shared Materials Instrumentation Facility at Duke University and National Center for CryoEM Access and Training and the Simons Electron Microscopy Center located at the New York Structural Biology Center, supported by the NIH Common Fund Transformative High Resolution Cryo-Electron Microscopy program (U24 GM129539) and by grants from the Simons Foundation (SF349247). Funding support: P01-AI131251 (G.M.S); UM1-AI100645 and UM1-AI44371 (B.F.H); R01-AI145687 and Translating Duke Health Initiative (P.A.); R37-AI140897, UM1-AI169633 and Duke School of Medicine Whitehead Scholarship (W.B.W); R01-AI120801 (K.O.S.); Duke CFAR. This study utilized the computational resources offered by Duke Research Computing (http://rc.duke.edu).

## Author contributions

W.B.W, G.M.S, and B.F.H. conceived and/or designed the studies, evaluated data, and co-wrote the paper. H.L., C.Z., S.W., W.D., Y.P, J.C., K.Wa, and E.E.G. performed viral Env sequencing and/or analyses. B.K. and B.Ha. provided support for viral Env evolution studies. P.A., S.G., J.L., K.Man. and R.J.E. performed and/or analyzed data from all structural studies. M.C., M.M., T.E., and B.H. performed BCR repertoire analyses and/or functional characterization of mAbs. Y.C., S.R., A.J.C., A.N.S., L.M., and M.P.H. supported Ig and/or transcriptome sequencing and analyses. K.W., M.B. and H.K. provided bioinformatic and computational biology support. N.D-R., J.M., C.J., K.O.S. and H.L. conducted and analyzed high-throughput neutralization assays. M.J.v-G., R.W.S. and K.O.S provided key reagents for functional assays. G.M.S. and H.L. designed SHIV infection of RMs. All authors edited and/or reviewed the manuscript.

## Competing interests

The authors have no competing interests.

## Resource availability

*<u>Lead Contact</u>*. Further information and requests for resources and reagents should be directed to and will be fulfilled by the lead contact, Wilton B. Williams. *<u>Materials Availability.</u>* The data presented in this manuscript, and research materials used in this study are available from Duke University upon request and subsequent execution of an appropriate materials transfer agreement. *<u>Data and Code Availability.</u>* The variable heavy and light chain gene sequences for recombinant DH1030 mAbs were provided in the manuscript (**Table S5**). Cryo-EM maps and models of DH1030.1 Fab bound to BG505 Env were deposited to the EMDB and PDB with accession numbers EMD-27628 and 8DP1, respectively. Cryo-EM maps and models of DH1030.1 Fab bound to CH848 Env were deposited to the EMDB and PDB with accession numbers EMD-27624 and 8DOW, respectively. The crystal structure of the DH1030.1 Fab was deposited to the PDB under accession code 8DKF.

## LIST OF SUPPLEMENTAL MATERIALS

**Figure S1.**
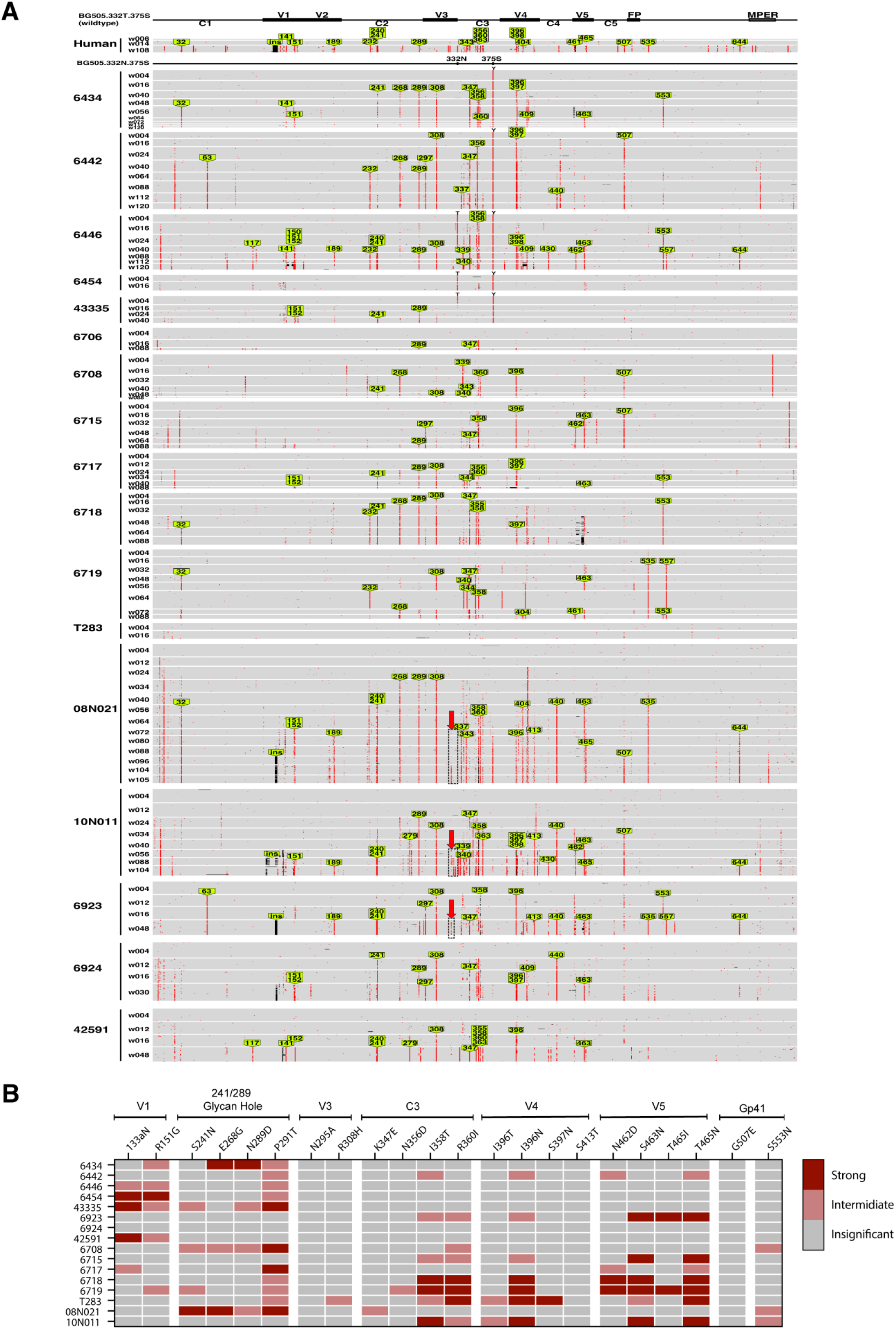
Conserved Env evolutionary patterns between the the HIV-1 infected human subject BG505 and SHIV infected RMs. **(A)** Pixel plots of gp140 SGS of human BG505 and 17 SHIV-BG505 infected RMs. Green arrows indicate conserved mutation sites with >50% TF loss and present in at least two monkeys. The black dotted boxes indicate the V3 glycan bnAb epitope region. **(B)** Plasma neutralization profiles against BG505 virus envelope carrying conserved mutations. Table shows the responses of plasma samples at 40 wpi to the mutants in different regions of the envelope. The changes in ID50s of the mutants relative to the BG505.T332N parental are shown here as relative ID50s (RID50s=ID50mutant/ID50parent). Strong, intermediate or insignificant changes in neutralization resistance are indicated in dark red, light red and grey, respectively.

**Figure S2.**
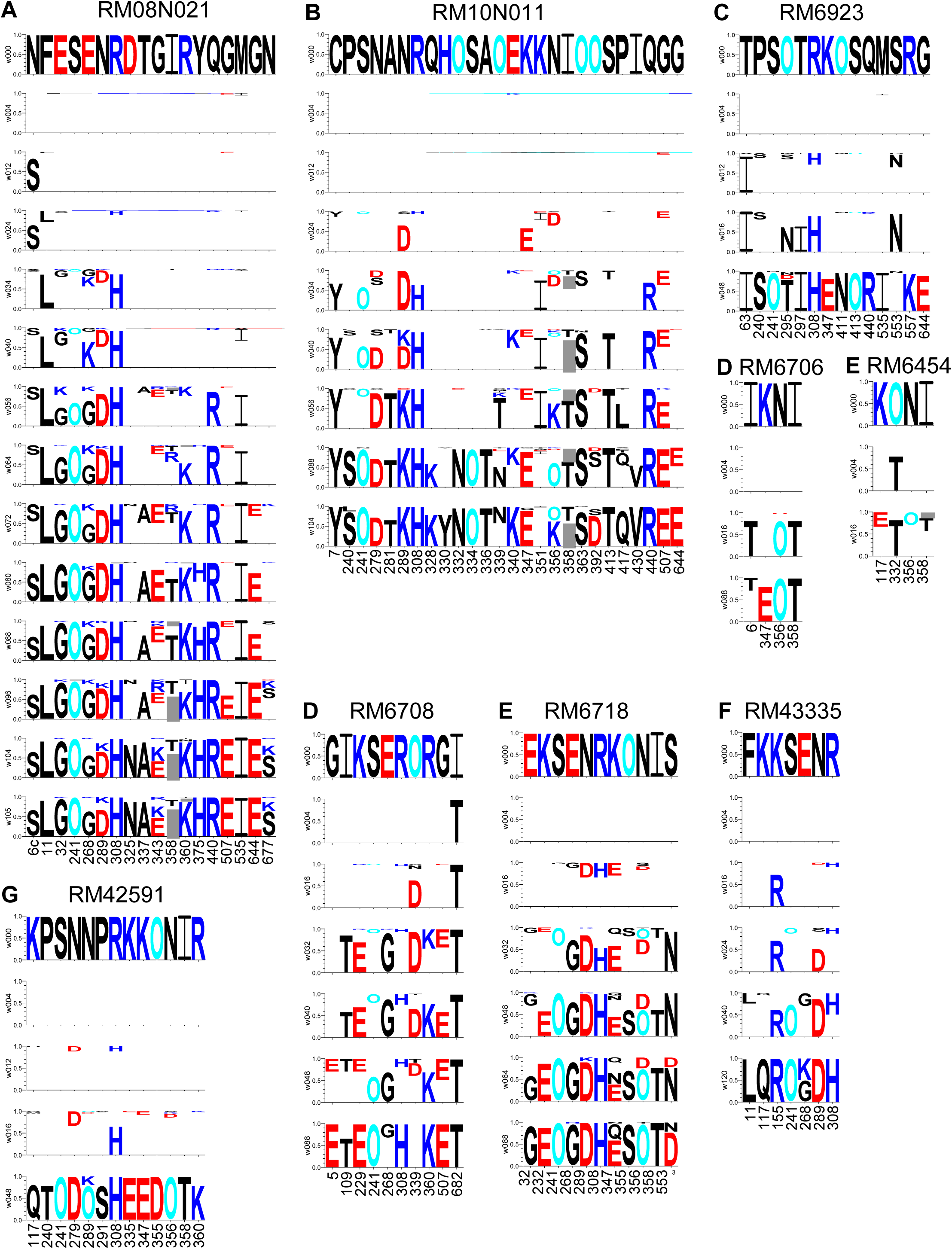
LASSIE selected sites in SHIV BG505 infected RMs (part 1). Mutations are shown as logos for each time point with the height of the mutation proportional to its frequency at a given time point. TF amino acids blanked out for longitudinal time points. O: potential N-linked glycosylation site. A threshold of 80% loss of the TF amino acid was used for LASSIE. The y-axis labels show the time points indicated as weeks post infection (e.g. w004 = 4 weeks post infection).

**Figure S3.**
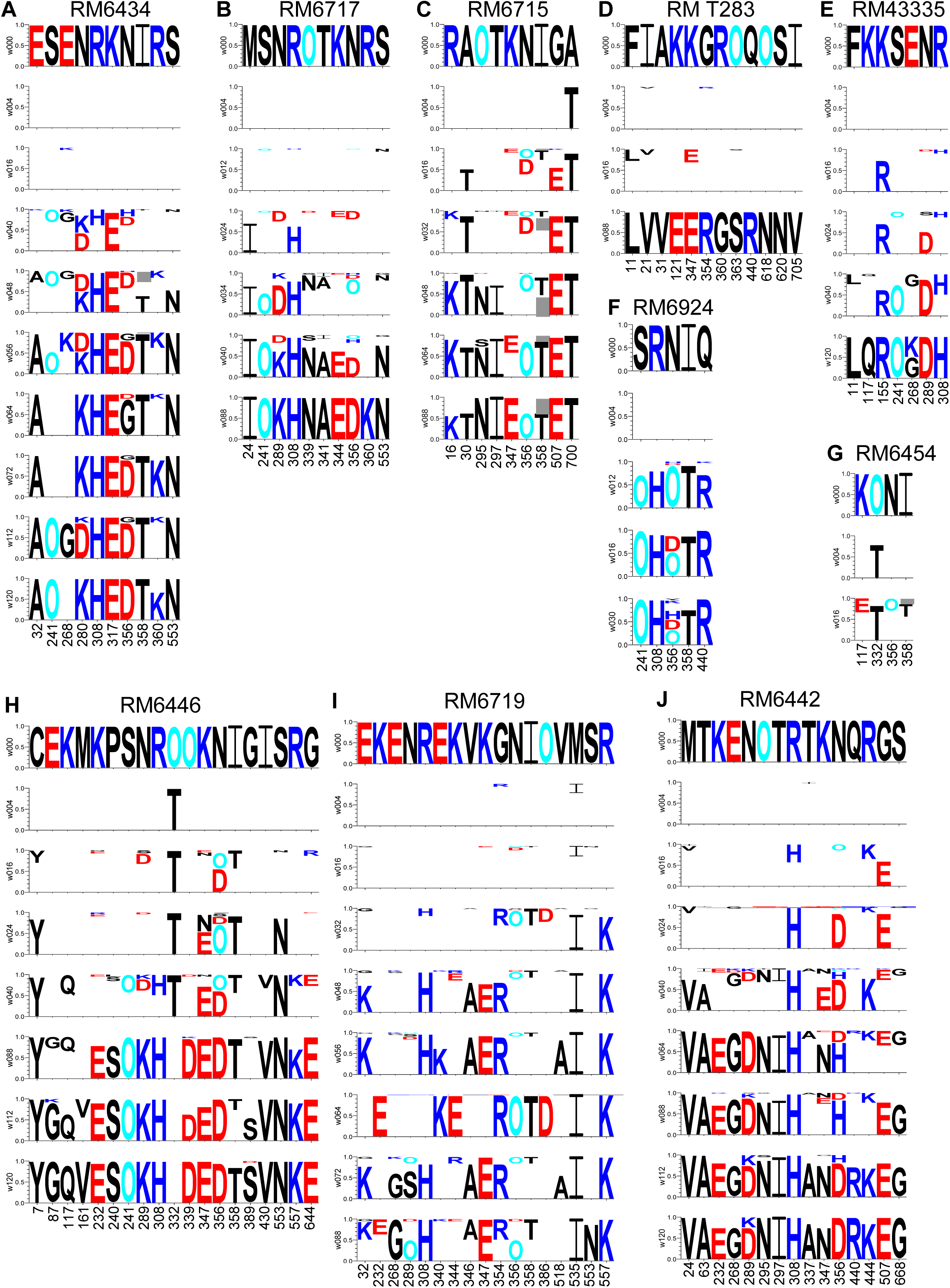
LASSIE selected sites in SHIV BG505 infected RMs (part 2). Same as Figure S2, but for the remaining RMs.

**Figure S4.**
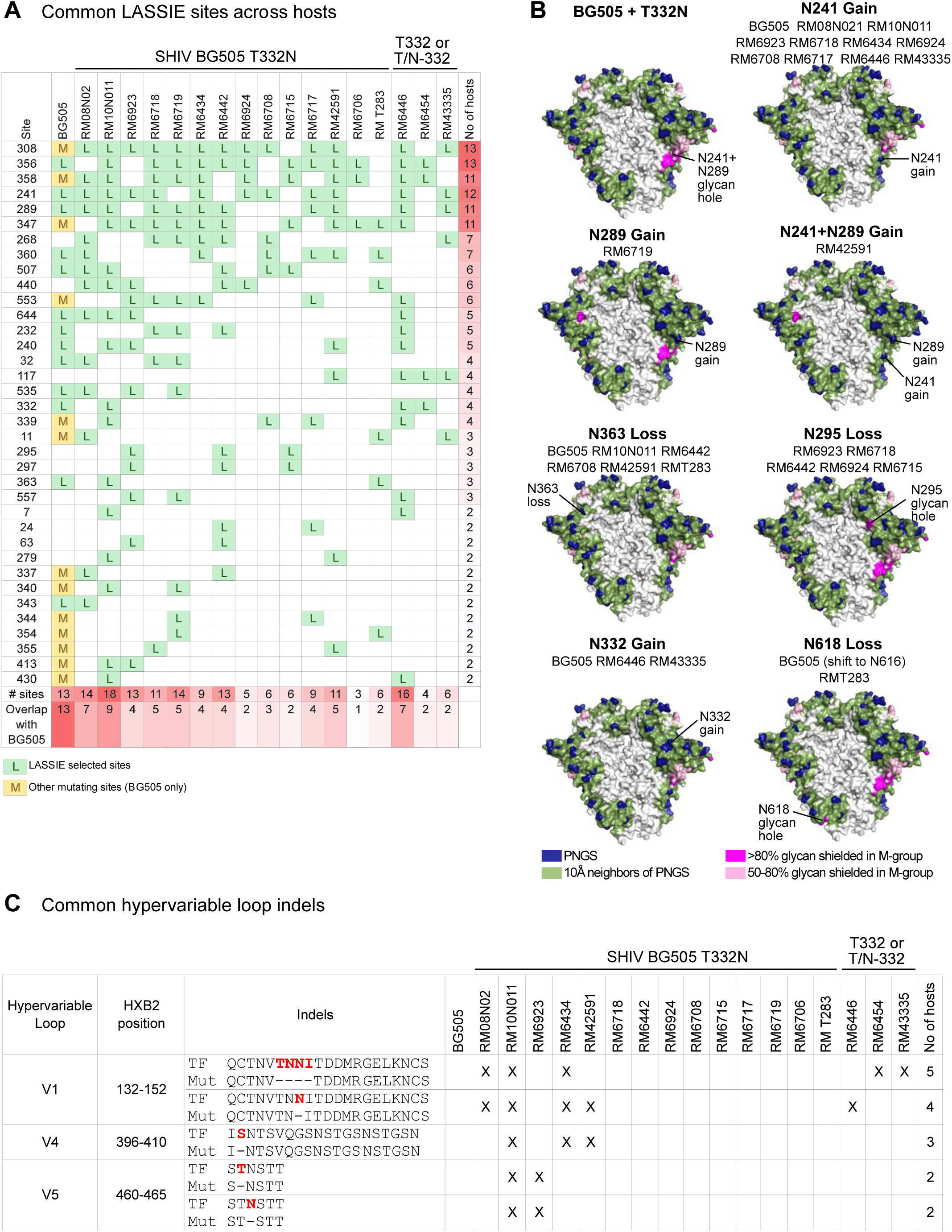
Common features of Env evolution across hosts. **(A)** Table of all LASSIE selected sites that were shared between 2 or more hosts. “L” indicates a LASSIE site under selection in the particular host. For BG505, “M” indicates that the site was mutating but not to the threshold of 80% loss of the TF form required by LASSIE. Last column shows the number of hosts in which each site was selected by LASSIE that was used to order the sites from the most commonly selected site to the least (note sites selected in only a single host are not shown). **(B)** Common patterns of glycan shield evolution. Representative patterns are shown with the hosts in which those patterns occur listed below the subpanel headings. The glycan shields shown in each subpanel are using time-point consensus glycans and hypervariable loop lengths from a single time point from a single host that showed the given pattern. **(C)** Common patterns of hypervariable loop indels across the hosts. For each host, hypervariable loops from Env sequences from the first year were manually aligned and analyzed for repeated indel patterns across hosts. All hypervariable loop indel patterns that were repeated across 2 or more hosts are shown. “X” indicates that the deletion was found in that host.

**Figure S5.**
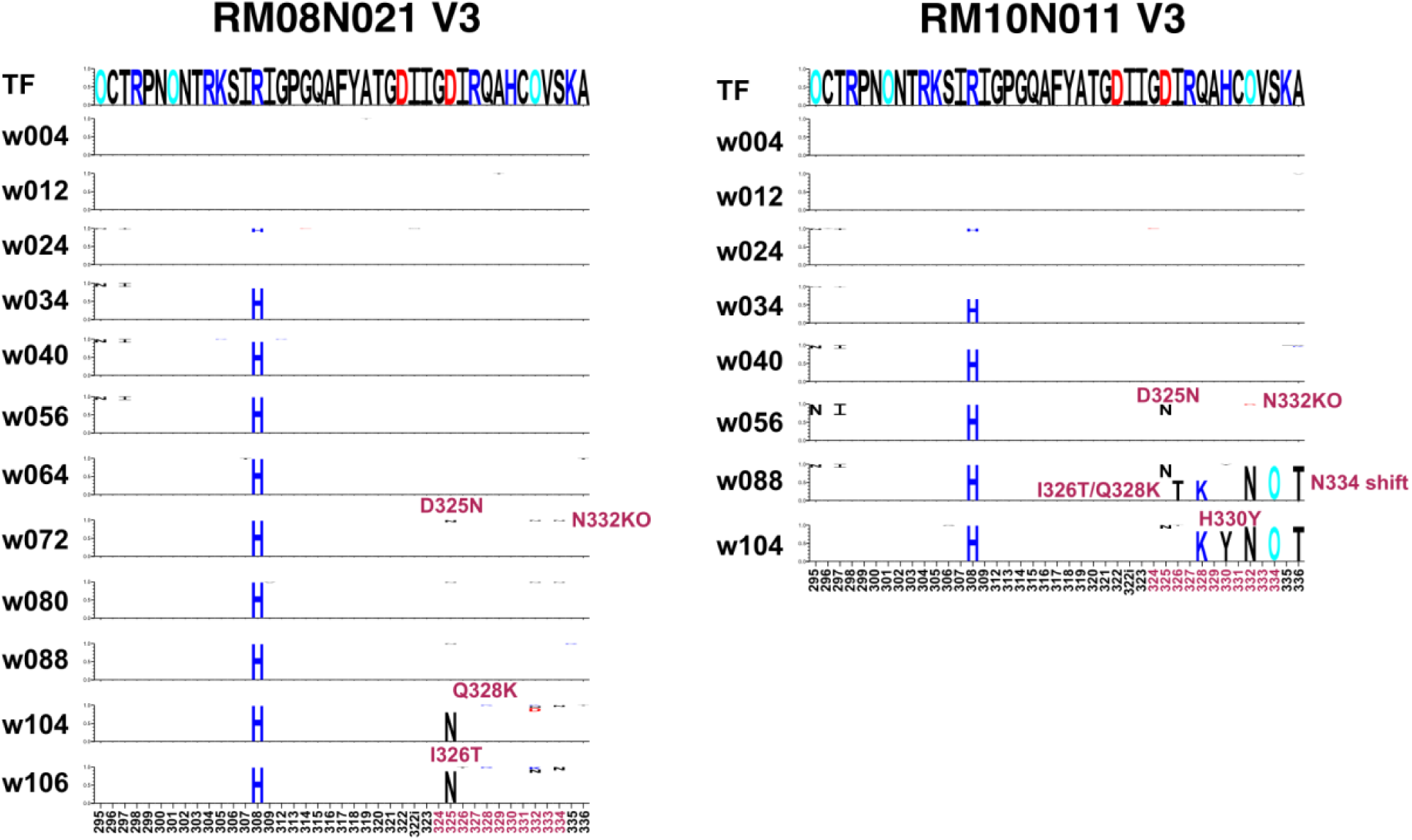
Mutations in the V3-glycan dependent bNAb epitope. Logo plots of sequence evolution in V3 region in macaques RM08N021 (left panel) and RM10N011 (right panel). The V3-glycan bnAb epitope residues are shown in red.

**Figure S6.**
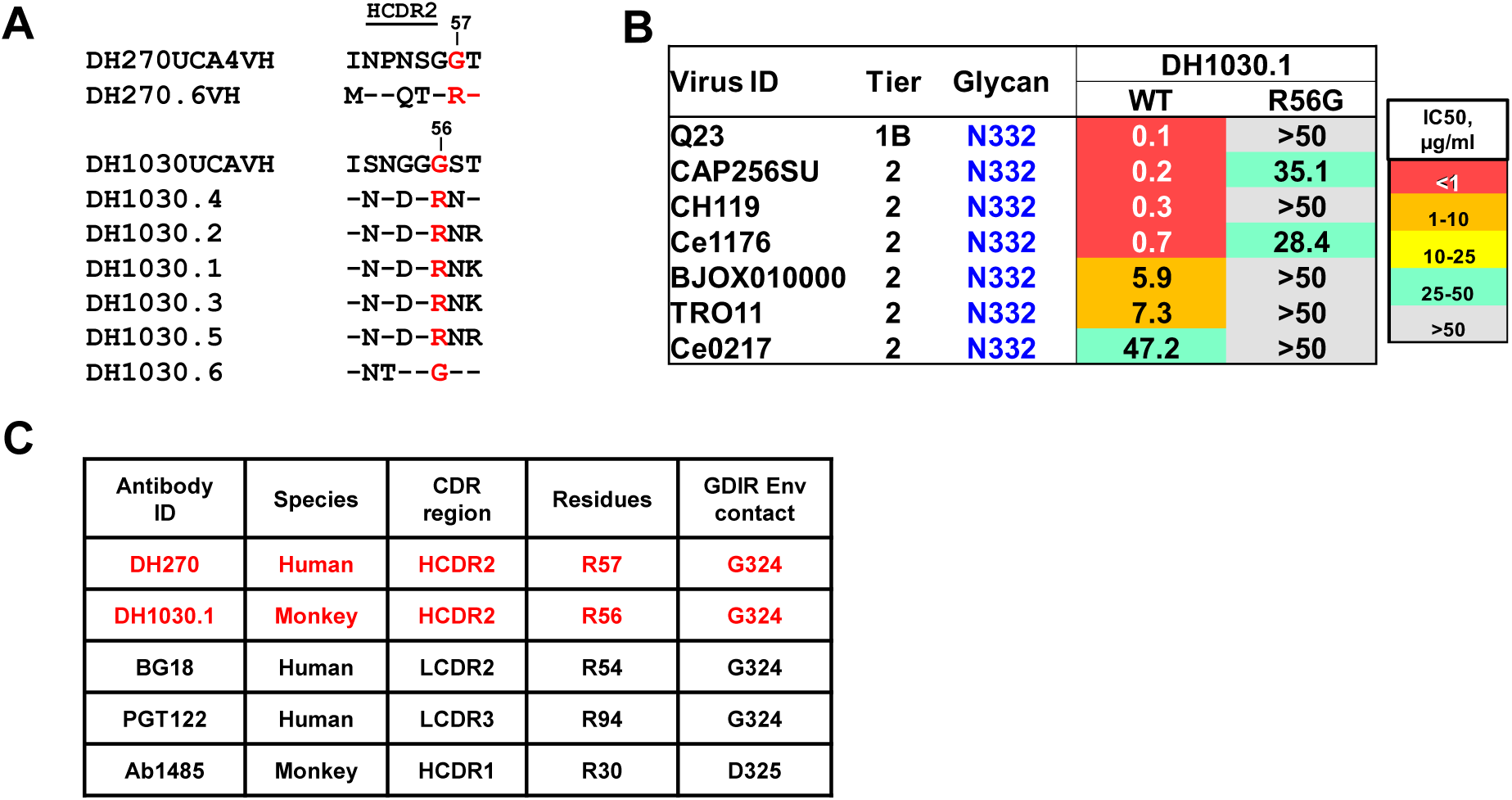
Comparison of human and macaque V3-glycan bNAbs. **(A)** Amino acid alignment of the HCDR2 for representative human-DH270, and macaque-DH1030 V3-glycan bnAb lineage members. Env positions of G56R in the macaque and G57R in the human heavy chain genes are indicated in red font. **(B)** Neutralization titers of wild-type DH1030.1 and a mutant DH1030.1 with R56G-revertant mutation in the heavy chain gene against a panel of heterologous HIV-1 strains bearing the Env N332 glycan. Neutralization was measured in TZM-bl cells and reported in IC50 (µg/ml). **(C)** List of human and macaque V3-glycan bnAbs with arginine residues that have been postulated or shown to mediate Env contact via the ^324^GDIR^327^–motif on Env. These residues have been tested for key interactions with Env in both DH270 and DH1030.

**Figure S7.**
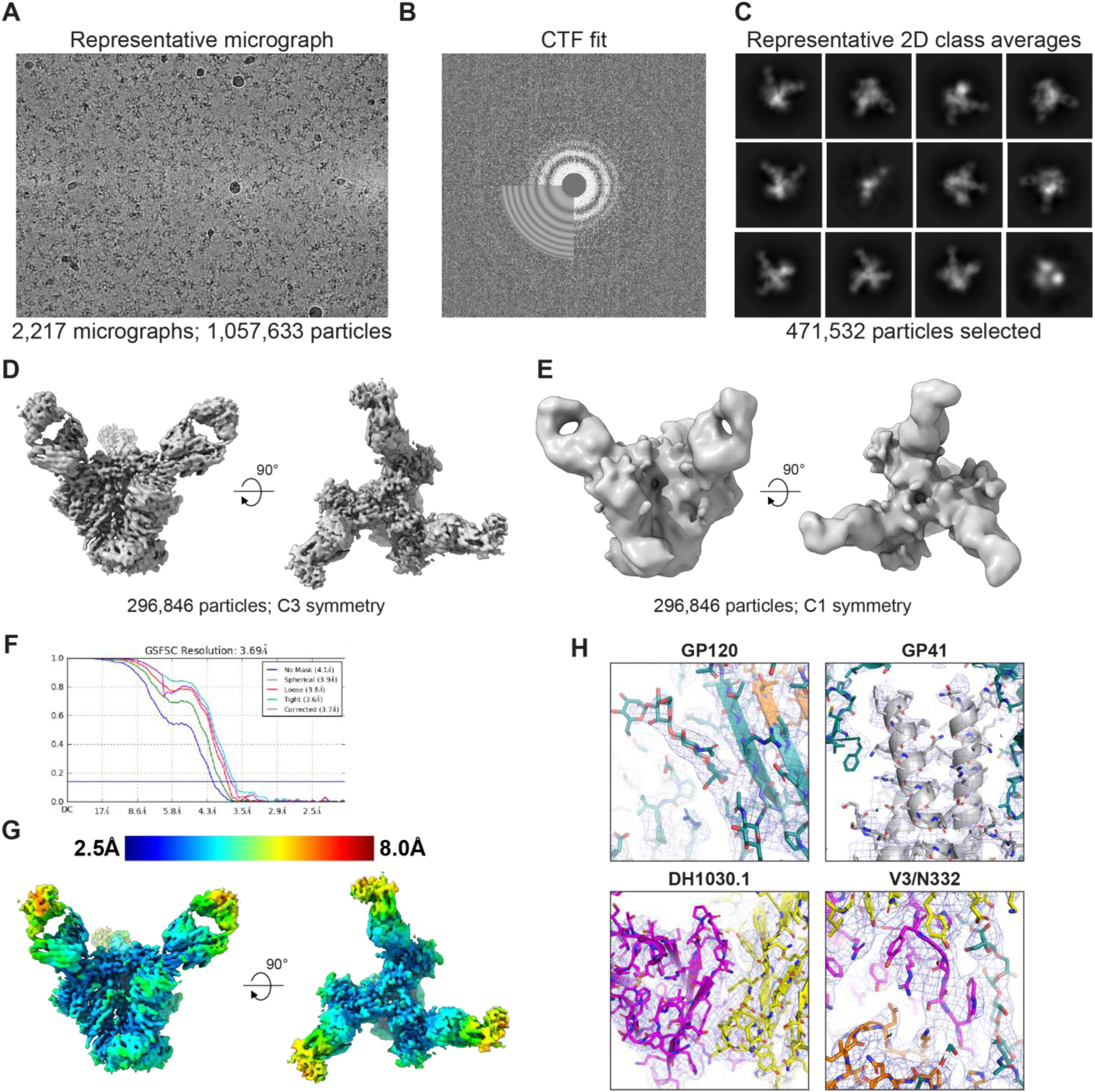
Cryo-EM data processing for the complex between the clade C Env CH848 10.17 DS.SOSIP_DT and DH1030.1 Fab. **(A)** Representative cryo-EM micrograph. **(B)** Cryo-EM CTF fit. **(C)** Representative 2D class averages from cryo-EM dataset. **(D)** Refined map. (E) *Ab initio* reconstruction. **(F)** Fourier shell correlation curve. **(G)** Refined cryo-EM maps colored by local resolution. **(H)** Zoom-in images showing the GP120, GP41, DH1030.1 LC-HC interface and GP120-V3 loop and glycan 332 regions in the structures. The cryo-EM map is shown as a mesh surface and the fitted model is in cartoon representation, with residues shown as sticks.

**Figure S8.**
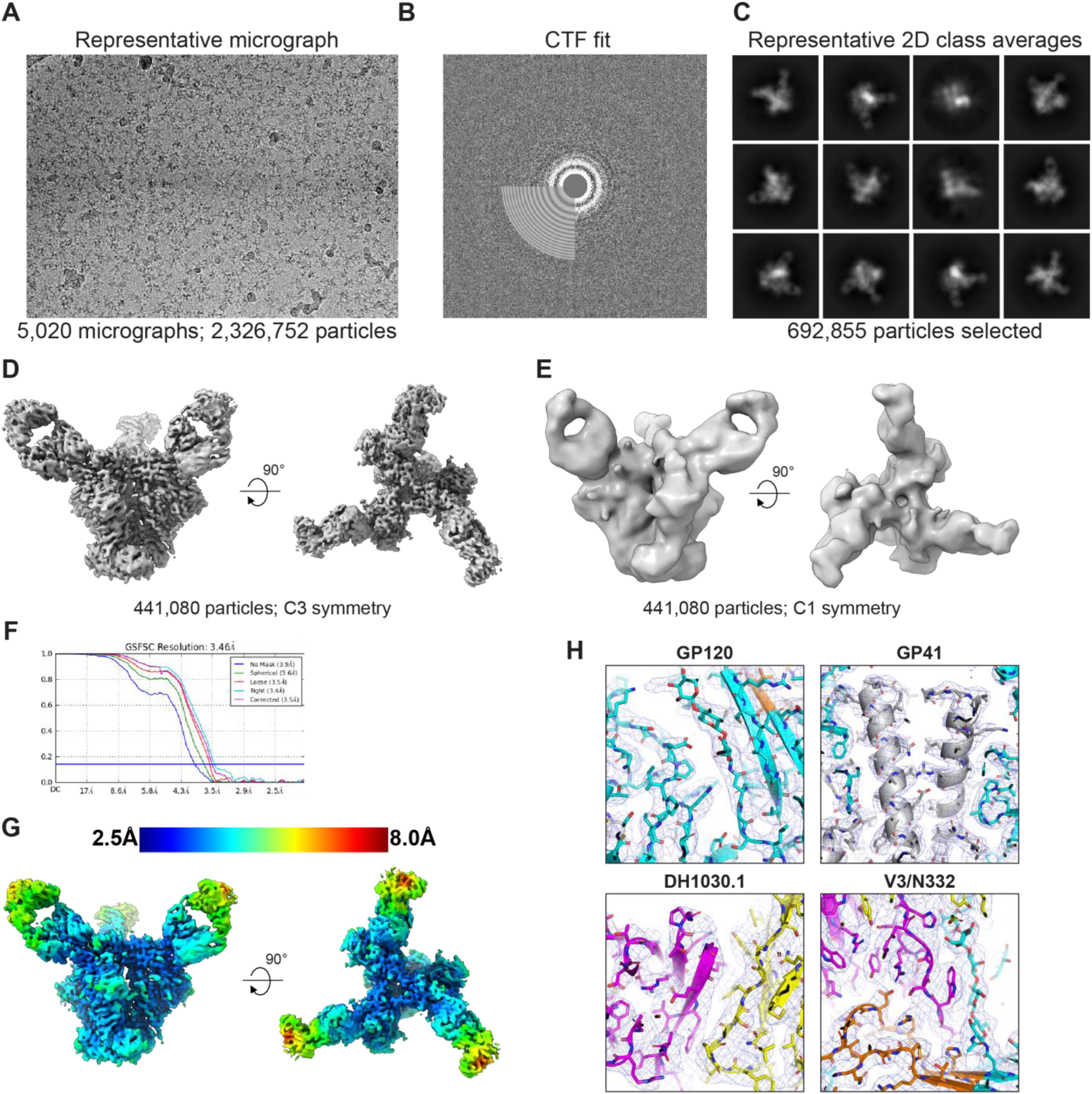
Cryo-EM data processing for the complex between the clade A Env BG505.T332N SOSIP and DH1030.1 Fab. **(A)** Representative cryo-EM micrograph. **(B)** Cryo-EM CTF fit. **(C)** Representative 2D class averages from cryo-EM dataset. (**D**) Refined map. (E) *Ab initio* reconstruction. **(F)** Fourier shell correlation curve. **(G)** Refined cryo-EM maps colored by local resolution. **(H)** Zoom-in images showing the GP120, GP41, DH1030.1 LC-HC interface and GP120-V3 loop and glycan 332 regions in the structures. The cryo-EM map is shown as a mesh surface and the fitted model is in cartoon representation, with residues shown as sticks.

**Figure S9.**
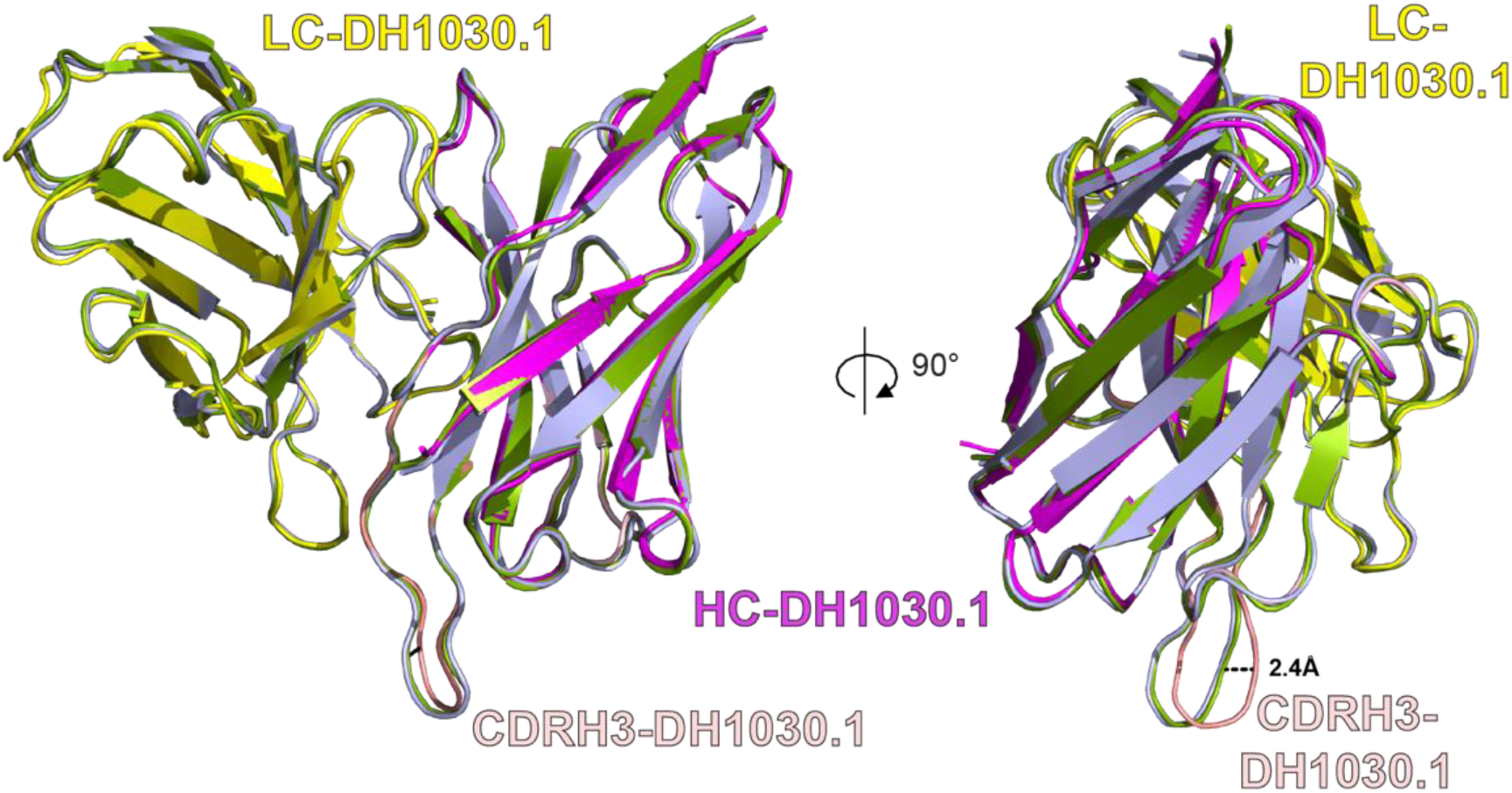
Structural comparison of DH1030.1 Fab variable region in its Apo form (crystal structure) and bound to CH848 10.17 DS.SOSIP_DT or BG505.T332N SOSIP (cryo-EM) forms. DH1030.1 Fab from the crystal structure in its Apo form has its light chain colored yellow and heavy chain magenta. The variable region of DH1030.1 from the complex with BG505 is colored wheat while the Fab from the complex with CH848 is colored pale purple. The RMSD between the crystal structure and complex with BG505 or CH848 were 0.5 and 0.6 Å respectively. The Super sequence-independent alignment in pymol was used to align the structures.

**Figure S10:**
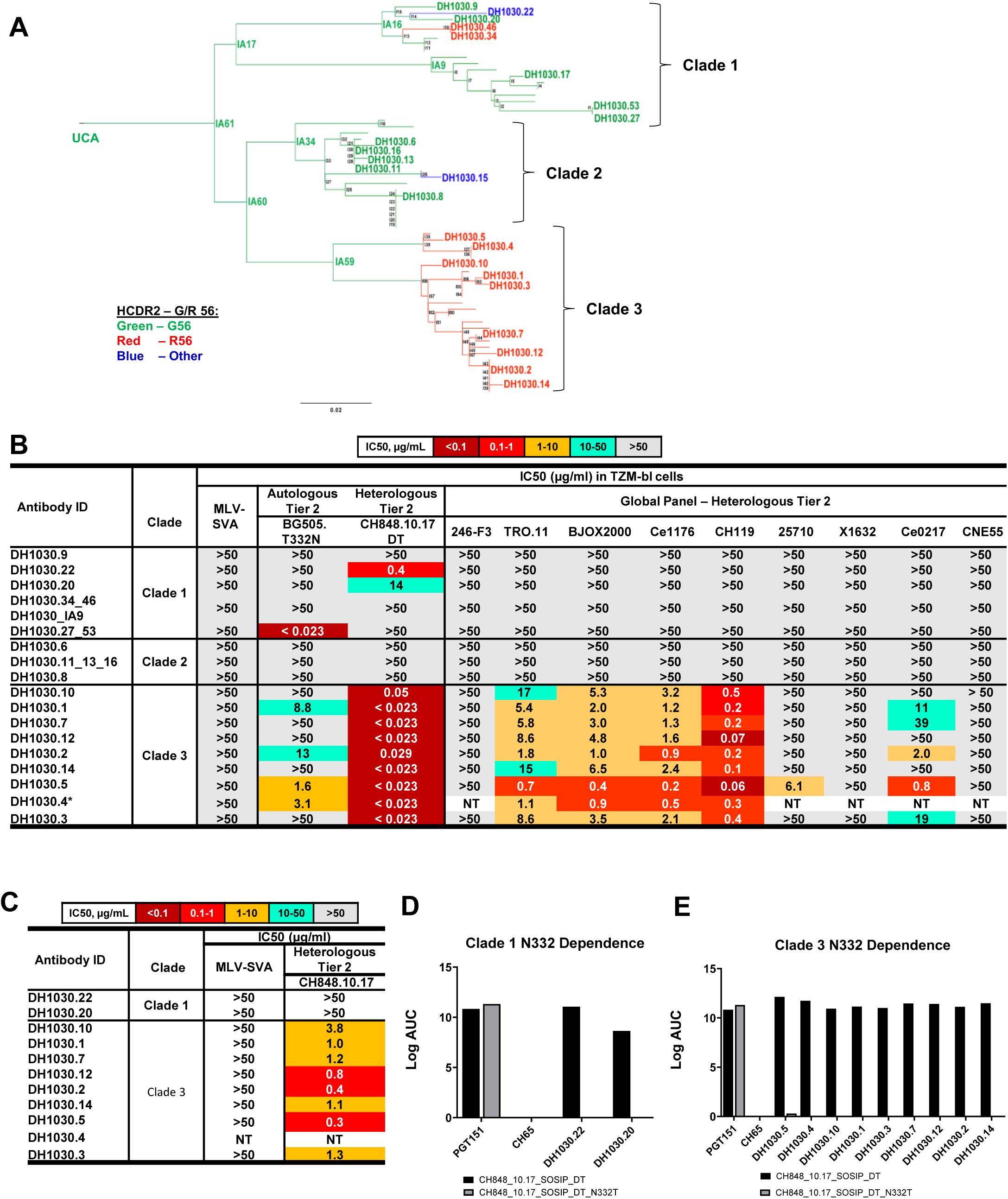
Characteristics of DH1030 mAbs with different genetic signatures. **(A)** A phylogram of the 62-member DH1030 V3 glycan bnAb lineage that was isolated from a SHIV BG505.T332N infected rhesus macaque. The phylogram is divided into three clades based on clustering of antibody sequences; clades 1, 2, and 3. Clade 3 represented the bnAb clade containing Abs that neutralized heterologous tier 2 HIV-1 strains. The lineage members indicated on the phylogram represent 27 recombinant monoclonal antibodies that were functionally characterized. **(B)** Heatmap summarizing the neutralization titers of DH1030 lineage Abs against a panel of HIV-1 strains, including the autologous and heterologous tier 2 HIV-1 strains. Heterologous tier 2 HIV-1 strains tested were from the global panel of HIV-1. CH848 10.17_N133D_N138T (DT) was previously shown to be sensitive for neutralization by precursors and mature Abs of DH270 V3-glycan bnAb lineage. Neutralization titer was reported as IC50 (μg/mL). DH1030.4 had insufficient yield to screen against all strains. **(C)** For DH1030 mAbs that neutralized CH848 10.17 DT, we tested them for neutralization of CH848 10.17 bearing V1 glycans. Heatmap summarizing the neutralization titers of DH1030 lineage Abs against heterologous tier 2 CH848 10.17. Neutralization titer was reported as IC50 (μg/mL). MuLV was used as a non-HIV negative control. **(D and E)** Log AUC graphs depicting Clade 1 and Clade 3 neutralizing mAbs binding against CH848 10.17 DT SOSIP and CH848 10.17 DT SOSIP N332T with PGT151 as a positive binding control and CH65 as a negative binding control.

**Figure S11:**
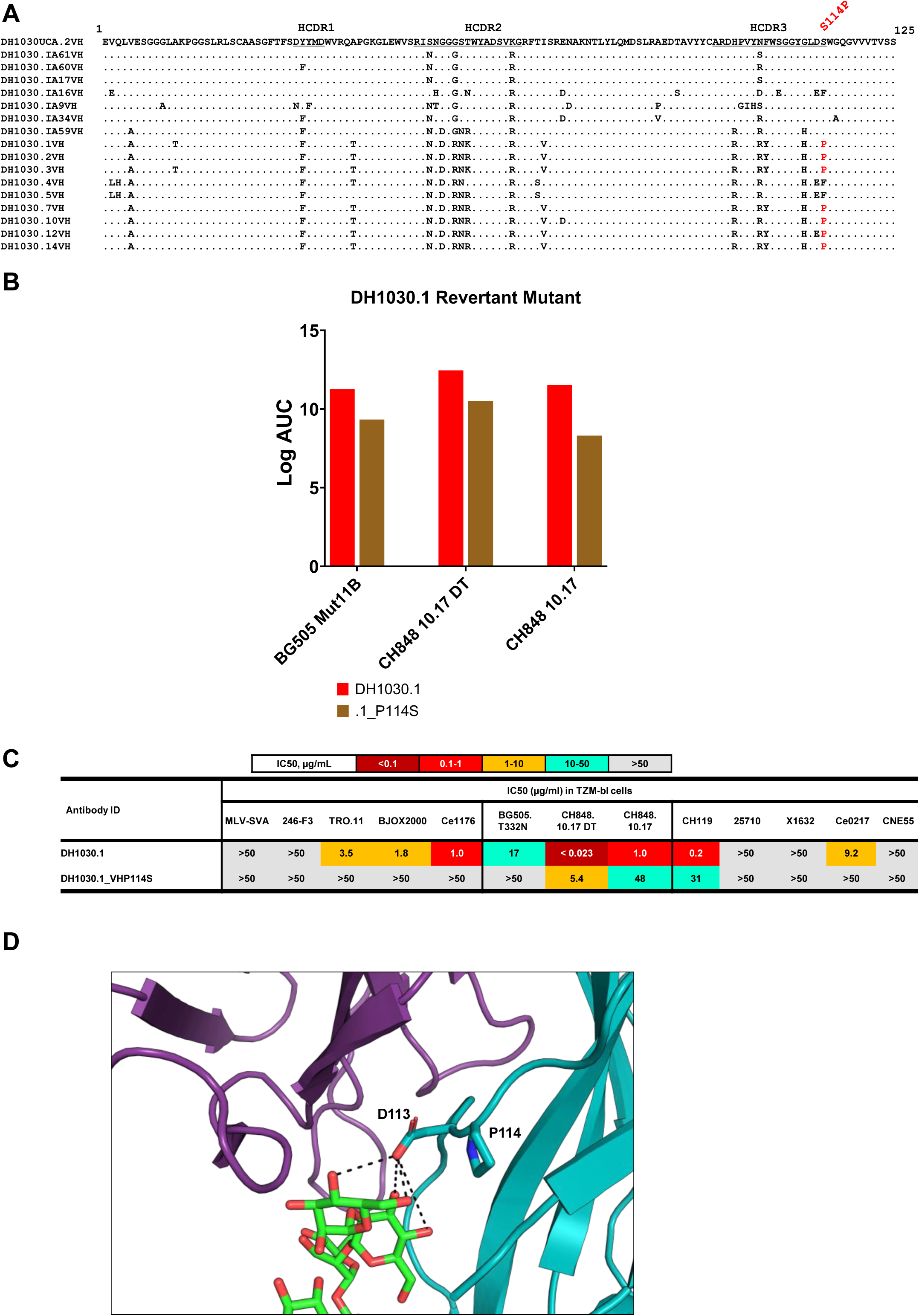
VH mutations associated with Clade 3 bnAb maturation. **(A)** VH alignment of the clade 3 DH1030 bnAbs relative to the putative UCA. Proline residue in the HCDR3 at position 114 is shown as a mutation from serine in the putative UCA. Sequences shown include the putative UCA and intermediate Abs for different clades of the phylogram. S114P was only found in the clade 3 bnAbs. Amino acid positions numbering positions from 1-125 indicated. **(B)** Bar graphs depicting mAb binding titers to recombinant HIV-1 Envs. Binding was measured in ELISA and reported as Log AUC titers. **(C)** Heatmap of IC50 (μg/mL) values for neutralization of DH1030 wild-type and mutant mAbs tested against heterologous HIV-1 strains from the global panel, autologous HIV-1 strain, and a V3-glycan bnAb precursor sensitive virus CH848 10.17 DT, in TZM-bl cells. **(D)** PyMol model of Cryo-EM representation of DH1030.1 Fab in complex with the autologous HIV-1 BG505.T332N SOSIP trimer. The heavy chain is in cyan and the light chain is in purple, with the N332-glycan on the SOSIP represented with green sticks. This representation depicts the D113 and P114 (blue region) residues on the heavy chain, with the D113 forming potential hydrogen bonds (H-bonds) with the N332-glycan; here P114 residue introduces a bend in the heavy chain that brings D113 residue within H-bonding distance with N332 glycan.

**Figure S12.**
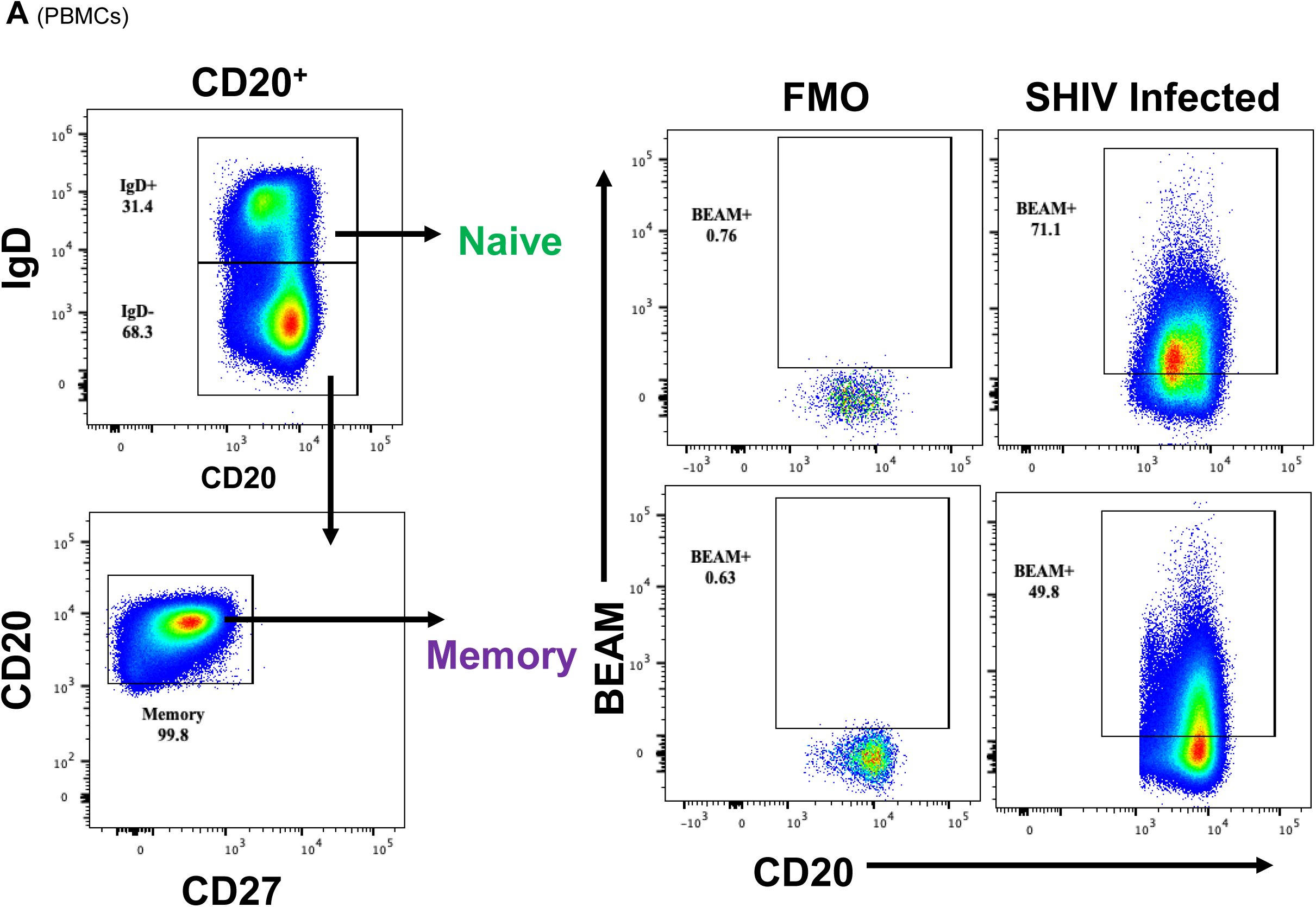
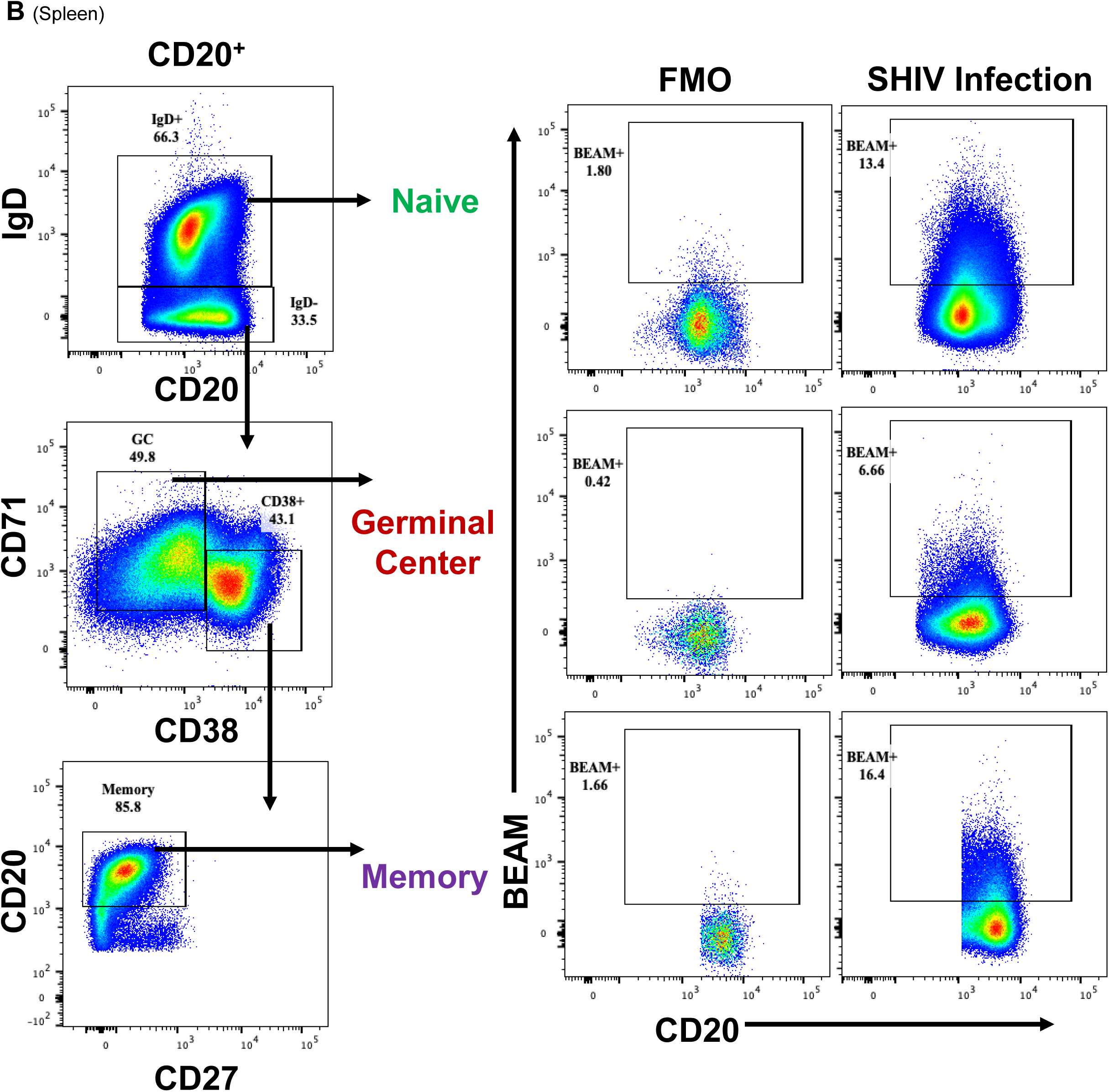
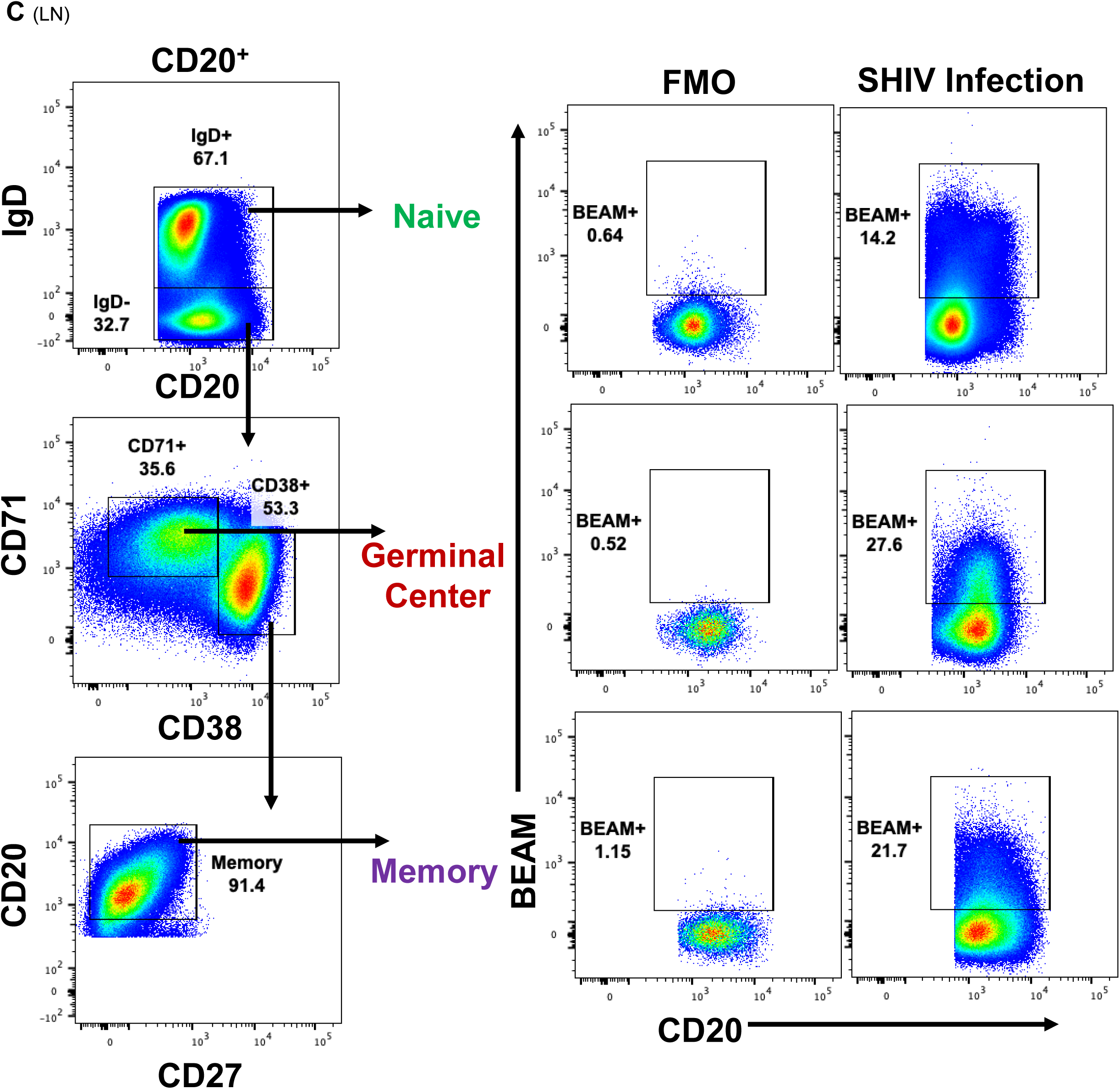
B cell phenotypes in blood and tissues. Flow cytometry gating used to define different B cell subsets in PBMCs **(A)**, spleen **(B)** and LN **(C)**. We studied naïve and memory B cell subsets in all samples. GC B cells were studied in LN and spleen. For the FMOs, we showed flow cytometry gating of cells from SHIV-infected RM 08N021 that did not have antigens added unlike the cells used for BEAM (column shown as SHIV). We observed BEAM+ naïve B cells in all compartments that we hypothesize are due to avidity effects by IgM B cells given a high frequency of IgM BEAM+ B cells from our experiences using BEAM to interrogate antigen-reactive B cells. Using the BEAM conjugate lacking an antigen as a negative control, we were able to subtract background or non-specific binding due to these avid IgM responses, thus our output for BEAM+ IgG B cells are predominantly high affinity binders.

**Figure S13.**
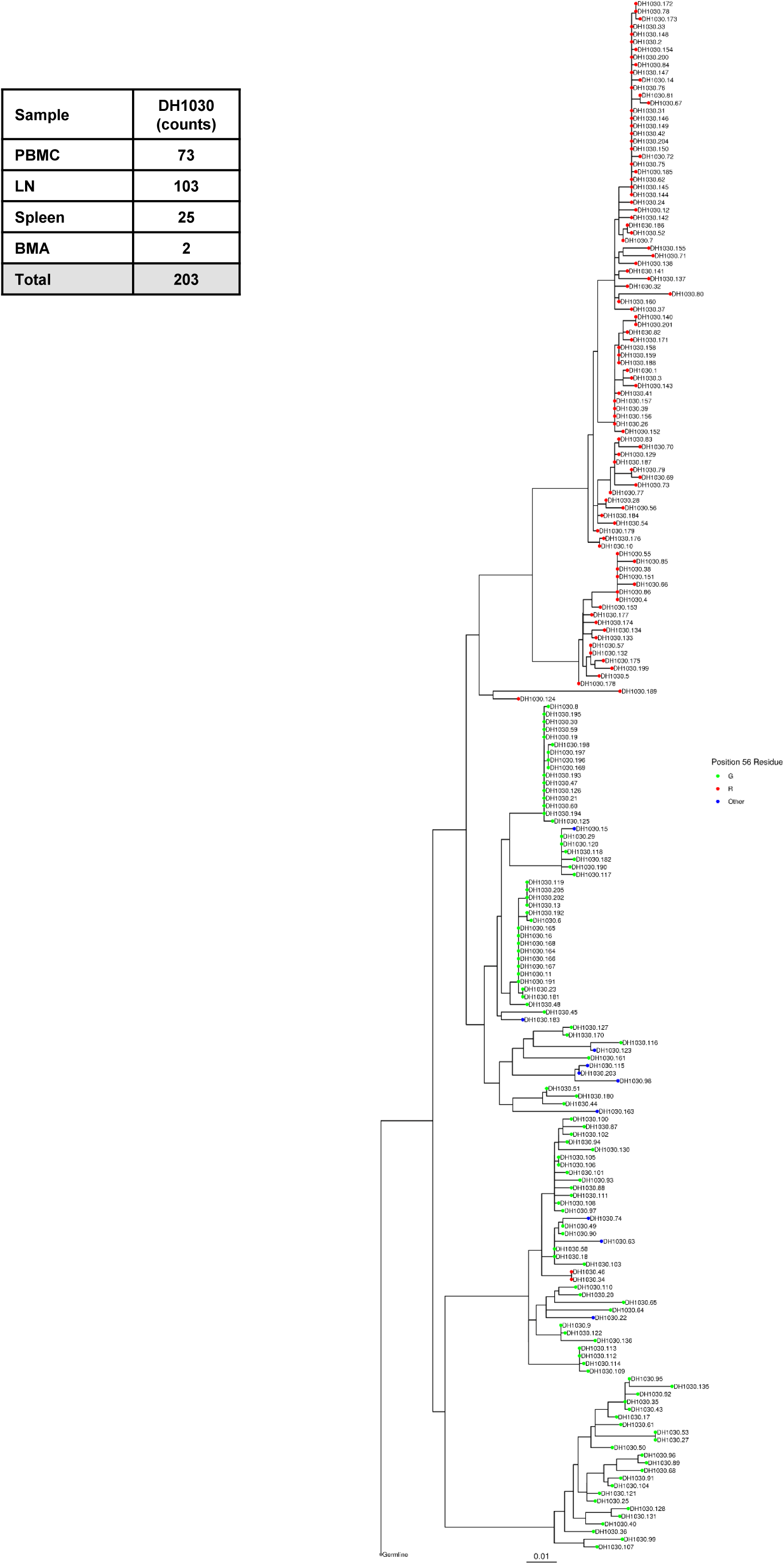
Phylogram of DH1030 lineage B cells. **(A)** Phylogenetic tree of DH1030 lineage members color coded by HCDR2 position 56; glycine, G (green); arginine, R (red); or other (blue) amino acid. Table insert indicates tissue origin for total DH1030 lineage members (n=203) isolated across multiple assays.

**Figure S14.**
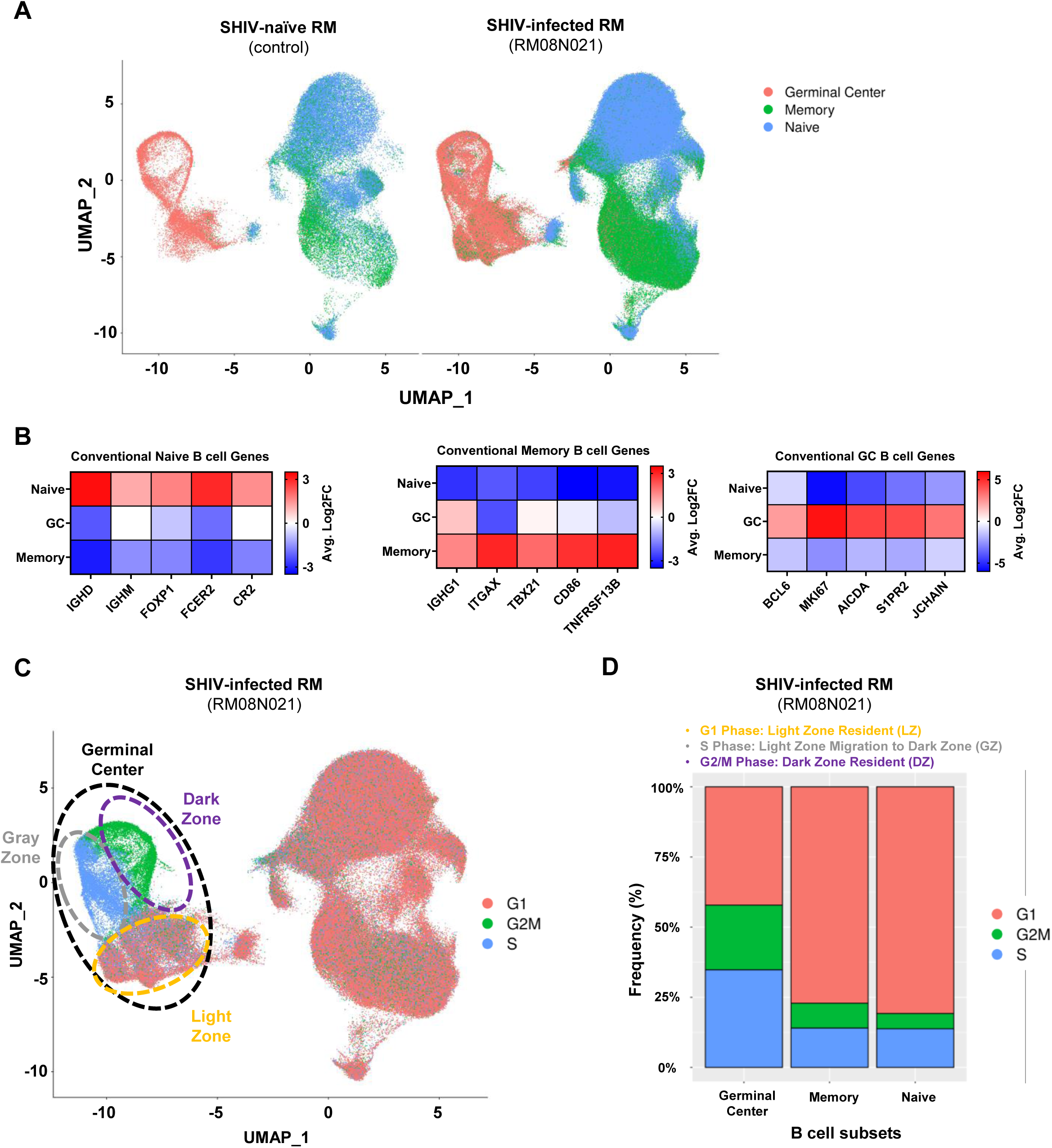
Transcriptional profiling of flow-sorted B cells. scRNA-seq of flow-sorted naïve, germinal center, and memory B cells from PBMC, lymph node, and spleen in RM 08N021 via PheBar-seq. **(A)** Integrated UMAPs of SHIV-naïve RM or SHIV-infected RM 08N021 color coding B cells sorted from naïve (blue), germinal center (pink), or memory (green) populations. UMAPs reflect integrated LN, spleen, and PBMC sorted B cells. **(B)** GEX profiling by average Log_2_ fold change (FC) for distinct lineage defining naïve, germinal center, and memory B cell genes. **(C)** Integrated UMAP of B cells in G1 (light red), G2M (green), and S (blue) phase of cell cycle among sorted B cells from LN, spleen and PBMCs. Dashed lines highlight clusters based on sort phenotype and GEX profiling in agreement with **(D)** bar graph showing frequency of flow-sorted naïve, germinal center, and memory B cells in G1, G2M, and S phase of cell cycle.

**Figure S15.**
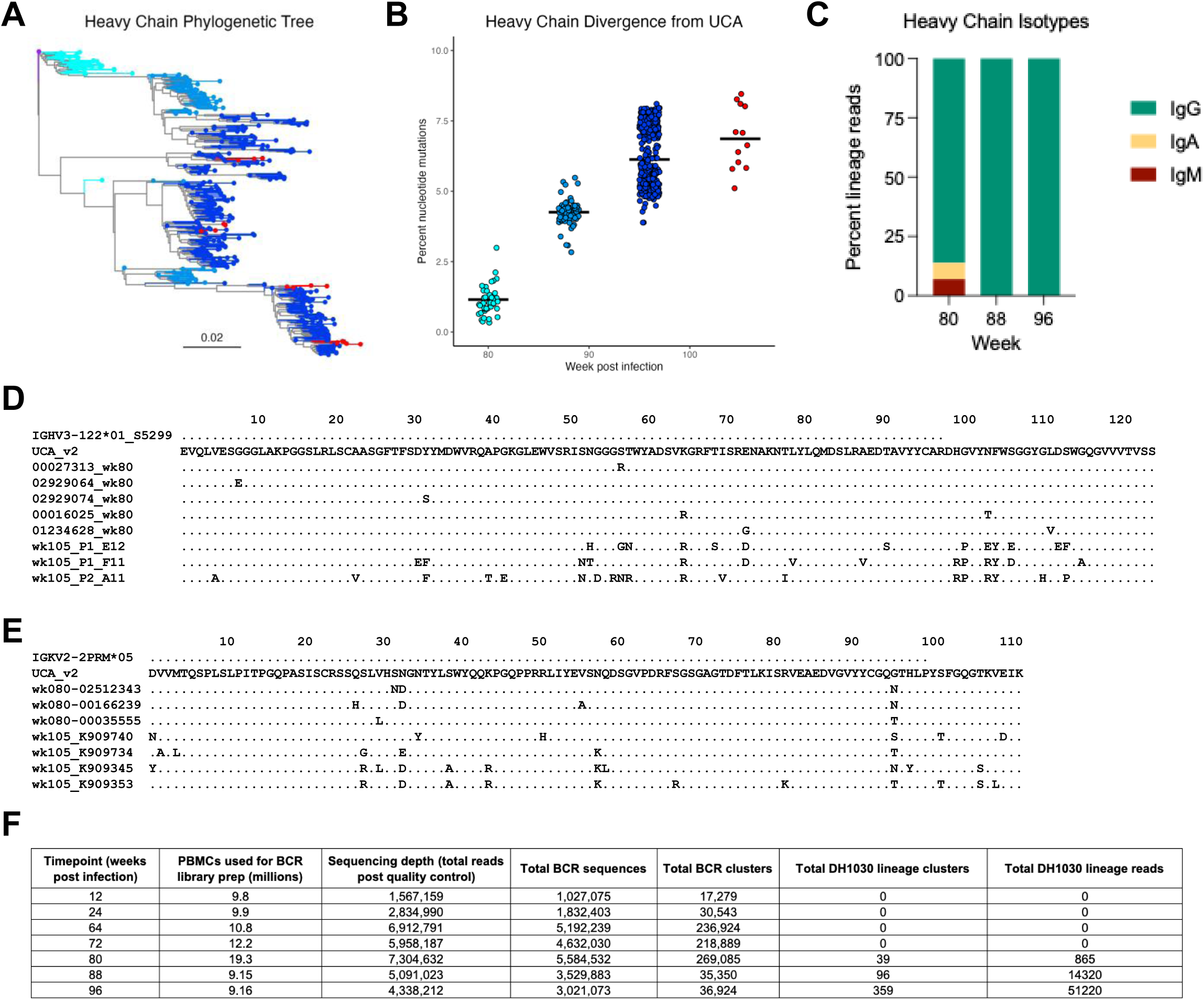
Lineage tracing supports priming between week 72 and 80 post infection. Next generation sequencing (NGS) of B cell receptor (BCR) sequences from peripheral blood mononuclear cells (PBMC) identifies DH1030 bnAb lineage members at weeks 80, 88, and 96 but not weeks 12, 24, 64, and 72. (**A**) Heavy chain phylogenetic tree of bnAb lineage rooted on the assigned V gene (purple), with unique BCR sequences from week 80 (cyan), 88 (light blue), and 96 (dark blue) NGS along with representative monoclonal antibodies from week 105 (red). (**B**) Nucleotide divergence from the inferred unmutated common ancestor (UCA) over time. Each point represents a unique heavy chain NGS sequence or monoclonal antibody sequence and each black line represents the within timepoint mean. (**C**) Percentages of DH1030 lineage heavy chain NGS reads assigned to each antibody isotype by timepoint. (**D, E**) Alignments of representative early and late DH1030 lineage members with their assigned heavy or light germline V gene. (**F**) Sequencing depth for each timepoint. Total BCR sequences includes duplicate and non-functional sequences, while total BCR clusters represents predicted functional antibody sequences deduplicated at a 99% identity. No clusters of the DH1030 lineage members were found between weeks 12 and 72.

**Table S1.**
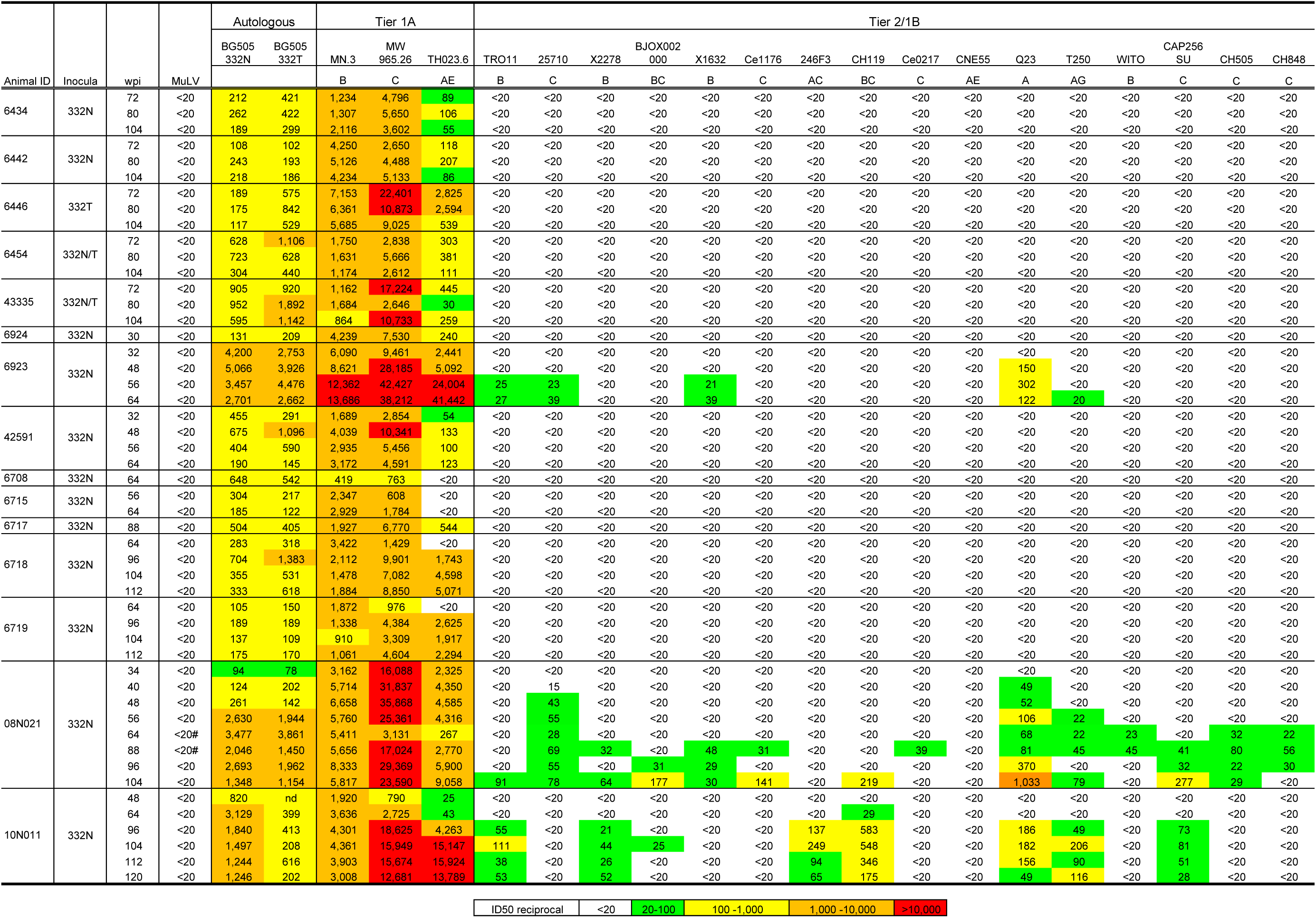
Autologous and heterologous neutralization by plasma from SHIV infected rhesus macaques.

**Table S2:**
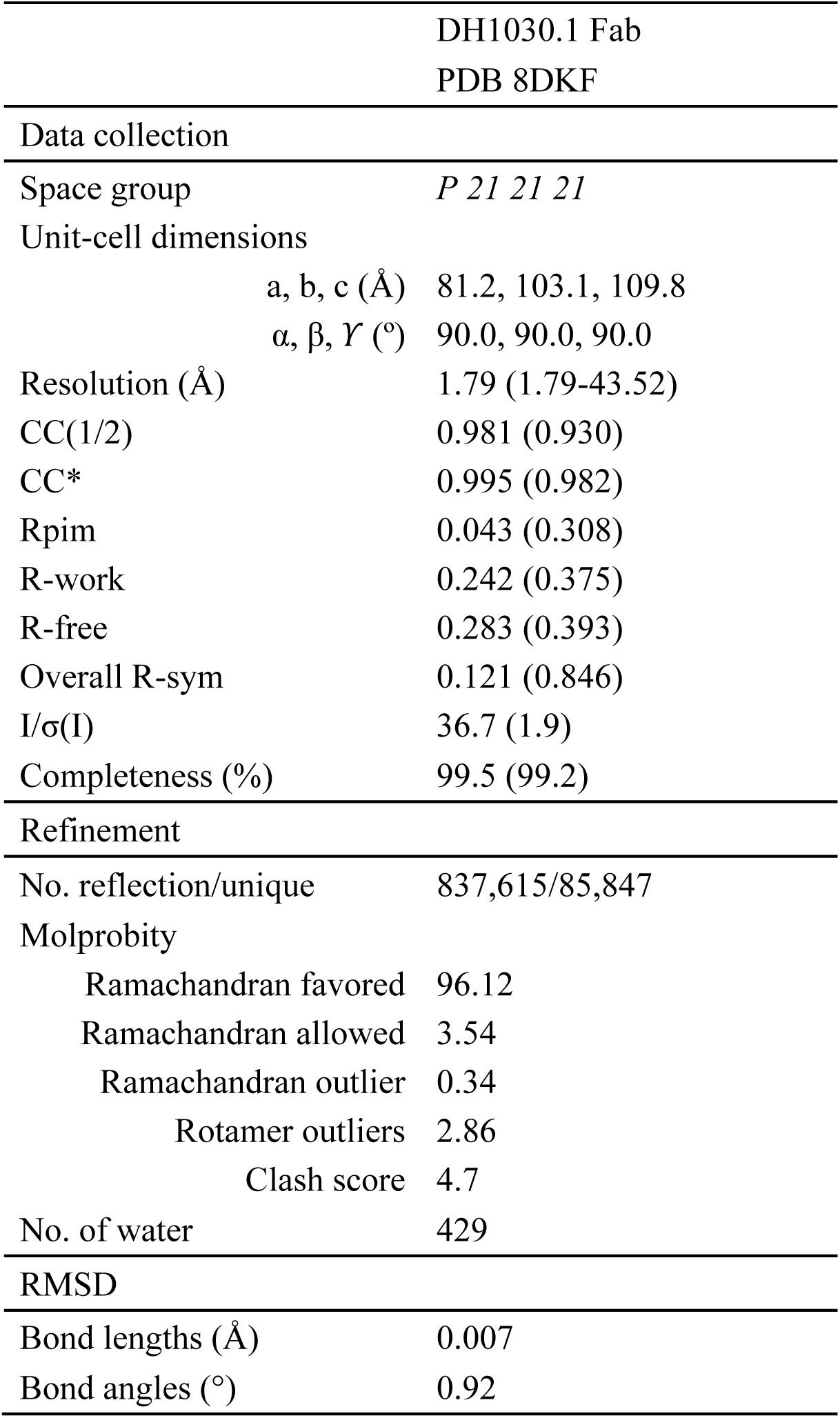
X-ray data collection and refinement statistics. Data collection and refinement statistics for DH1030.1 crystal structure.

**Table S3:**
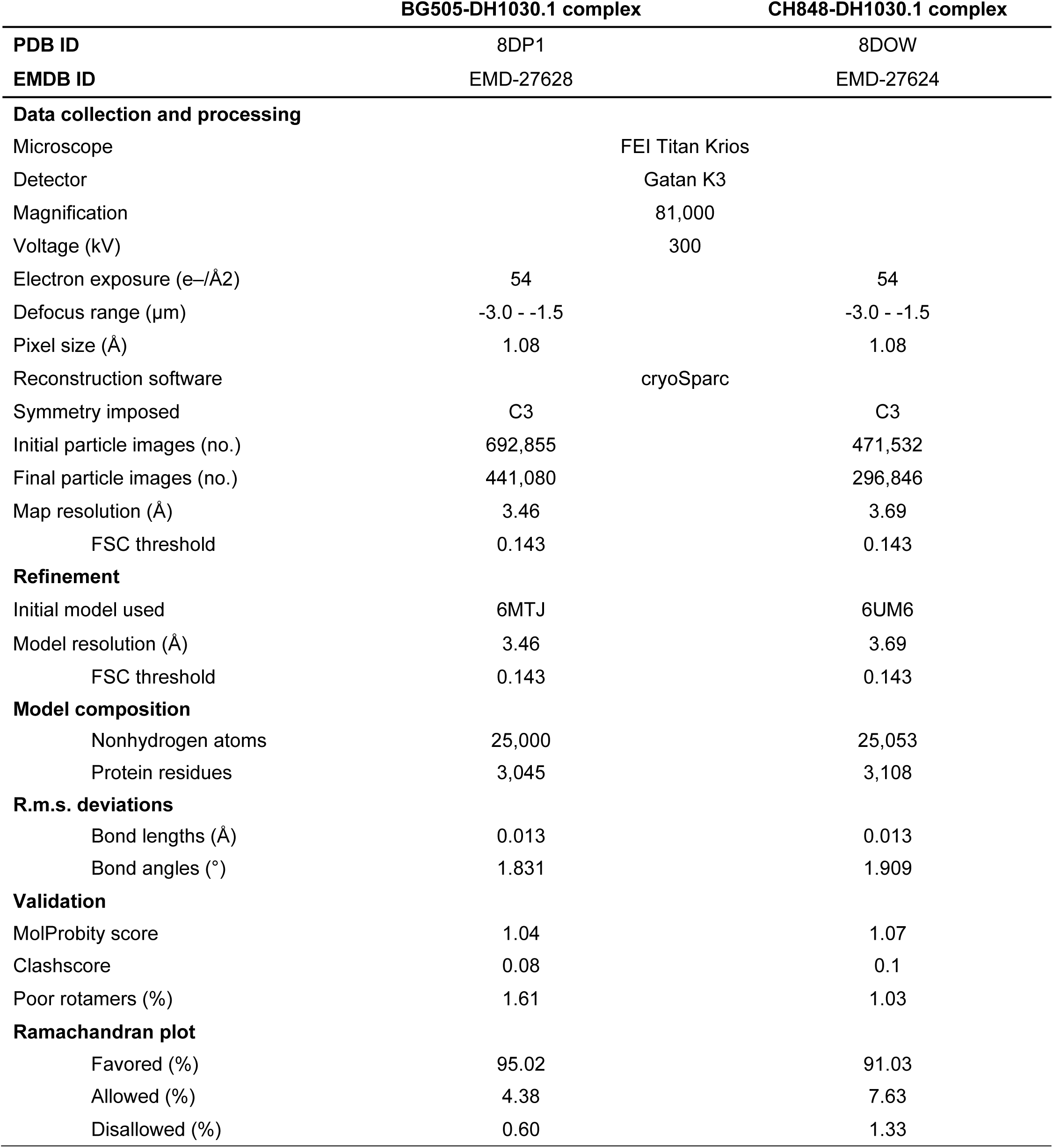
Cryo-EM Data Collection and Refinement Statistics. Cryo-EM data collection and refinements statistics for the complexes between DH1030.1 and the clade C Env CH848 10.17 DS.SOSIP_DT or clade A Env BG505.T332N SOSIP.

**Table S4.**
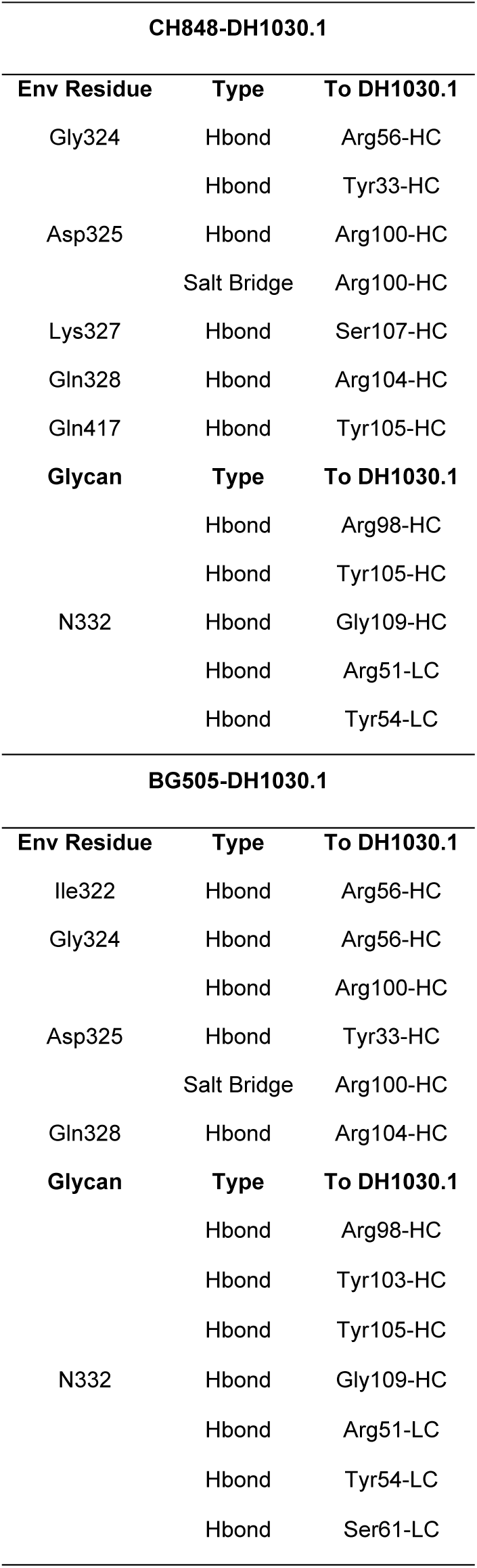
Contacts observed between DH1030.1 Fab and the CH848 10.17 DS.SOSIP_DT or BG505.T332N SOSIP. The PDBePISA tool (Proteins, Interfaces, Structures and Assemblies) was used to define the contacts.

**Table S5.**
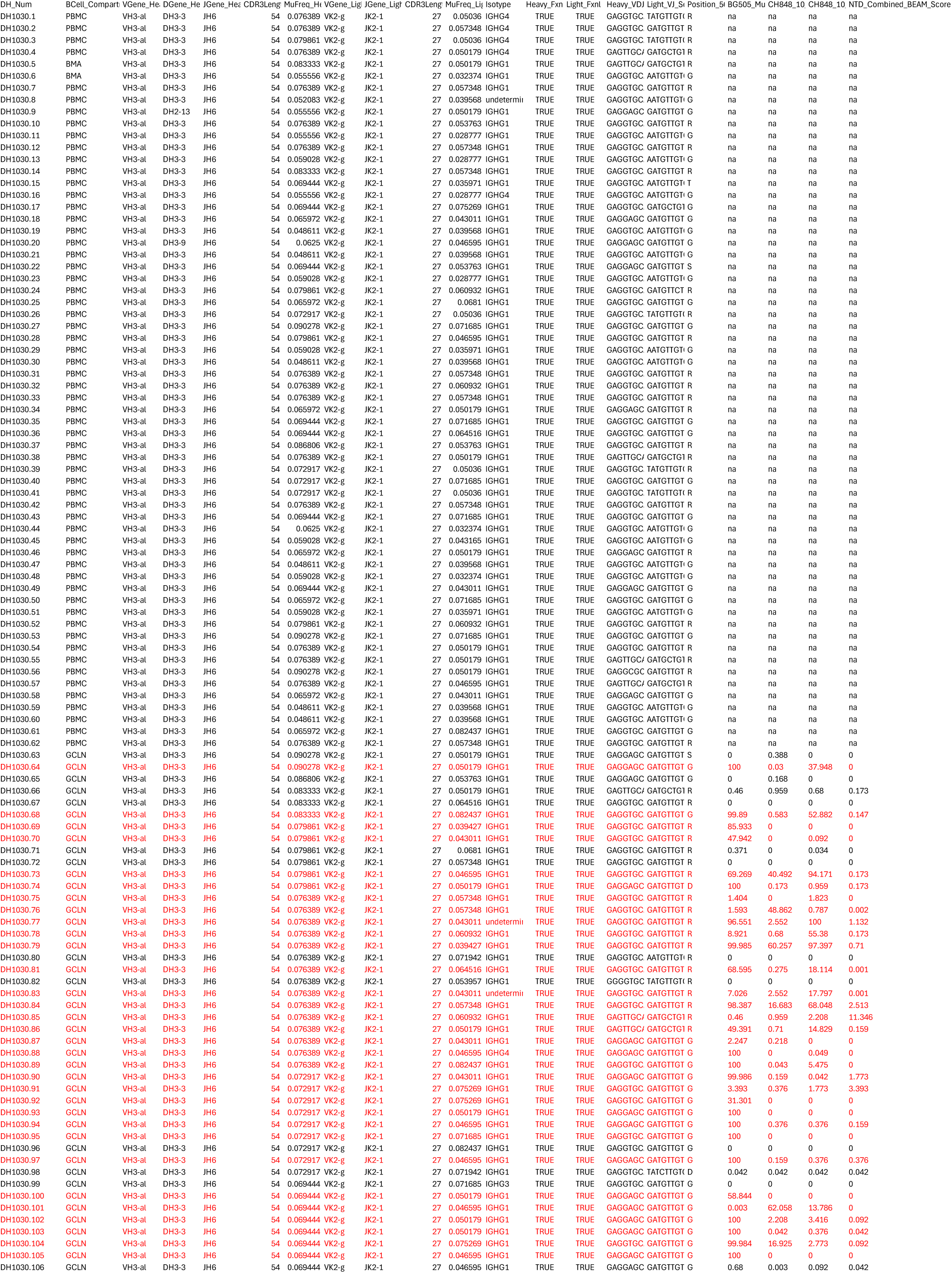
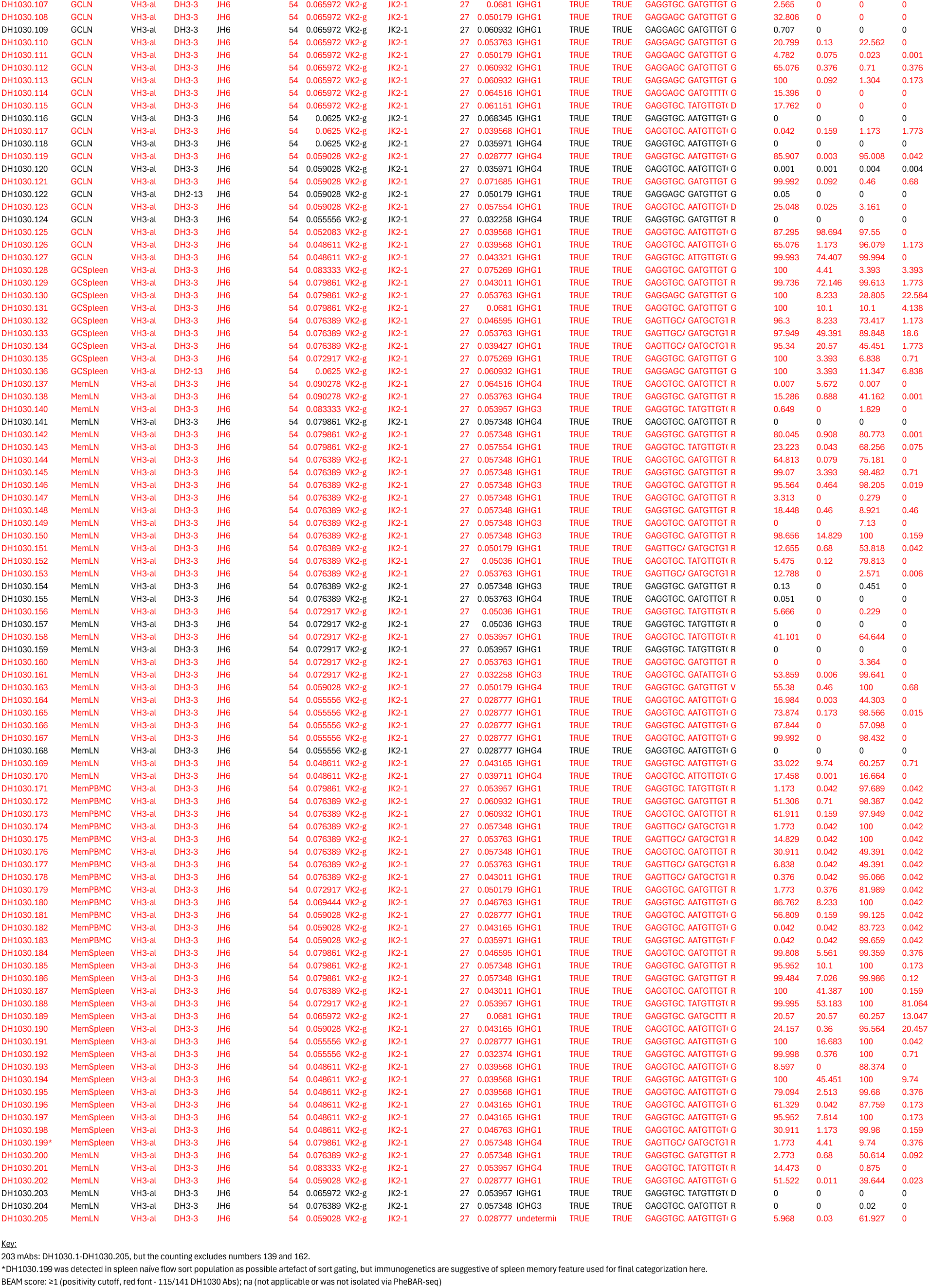
Immunogenetics of clonally-related DH1030 lineage members.

**Table S6.**
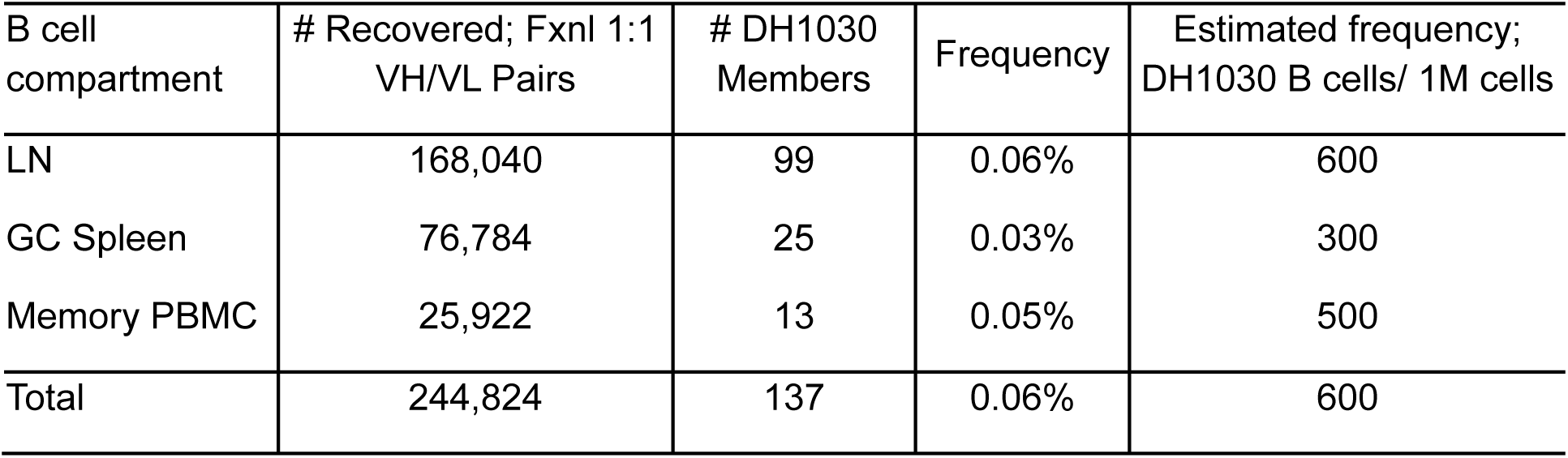
Frequency of DH1030 B cells in blood and tissues. Frequencies of DH1030 B cells among total B cells studied in different tissues and PBMCs via PheBAR-seq. Frequency reported as percent an estimated number of cells per 1 million (M) cells.

## Notes

### Competing Interest Statement

The authors have declared no competing interest.

## REFERENCES

1. Haynes BF, Burton DR, Mascola JR. Multiple roles for HIV broadly neutralizing antibodies. Sci Transl Med. 2019;11(516). doi: 10.1126/scitranslmed.aaz2686. PubMed PMID: 31666399; PMCID: PMC7171597.

2. Chuang GY, Zhou J, Acharya P, Rawi R, Shen CH, Sheng Z, Zhang B, Zhou T, Bailer RT, Dandey VP, Doria-Rose NA, Louder MK, McKee K, Mascola JR, Shapiro L, Kwong PD. Structural Survey of Broadly Neutralizing Antibodies Targeting the HIV-1 Env Trimer Delineates Epitope Categories and Characteristics of Recognition. Structure. 2019;27(1):196–206.e6. Epub 2018/11/21. doi: 10.1016/j.str.2018.10.007. PubMed PMID: 30471922; PMCID: PMC6664815.

3. Caillat C, Guilligay D, Sulbaran G, Weissenhorn W. Neutralizing Antibodies Targeting HIV-1 gp41. Viruses. 2020;12(11). Epub 2020/10/23. doi: 10.3390/v12111210. PubMed PMID: 33114242; PMCID: PMC7690876.

4. Skelly A, Gristick H, Li H, Gavor E, Connell A, Kreider E, Marchitto L, Hogarty M, Newby M, Allen J, Liu W, West Jr. A, Ayyanathan K, Campion M, Winters K, Gordon C, Osbaldeston R, Akeley M, Li Y, Singh A, Cruickshank K, Park Y, Zhao C, Li X, Amereh K, Van Itallie E, Carey J, Albertus A, DeLaitsch A, Keeffe J, Lituchy M, Walsh A, Morris D, Habib R, Bibollet-Ruche F, Mishra N, Avillion G, Koranda N, Plante S, Martella C, Lora J, Wang E, Lewis M, Martin M, Nussenzweig M, Seaman M, Irvine D, Wiehe K, Haynes B, Wagh K, Korber B, Andrabi R, Crispin M, Weissman D, Bjorkman P, Hahn B, Shaw G. Consistent Induction of Broadly Neutralizing HIV Antibodies by a Novel Two-Step Mechanism Informs Immunogen Design. bioRxiv preprint. 2025. doi: 10.1101/2025.10.06.680687.

5. Kong L, Lee JH, Doores KJ, Murin CD, Julien JP, McBride R, Liu Y, Marozsan A, Cupo A, Klasse PJ, Hoffenberg S, Caulfield M, King CR, Hua Y, Le KM, Khayat R, Deller MC, Clayton T, Tien H, Feizi T, Sanders RW, Paulson JC, Moore JP, Stanfield RL, Burton DR, Ward AB, Wilson IA. Supersite of immune vulnerability on the glycosylated face of HIV-1 envelope glycoprotein gp120. Nat Struct Mol Biol. 2013;20(7):796–803. doi: 10.1038/nsmb.2594. PubMed PMID: 23708606; PMCID: PMC3823233.

6. Sok D, Doores KJ, Briney B, Le KM, Saye-Francisco KL, Ramos A, Kulp DW, Julien JP, Menis S, Wickramasinghe L, Seaman MS, Schief WR, Wilson IA, Poignard P, Burton DR. Promiscuous glycan site recognition by antibodies to the high-mannose patch of gp120 broadens neutralization of HIV. Sci Transl Med. 2014;6(236):236ra63. doi: 10.1126/scitranslmed.3008104. PubMed PMID: 24828077; PMCID: PMC4095976.

7. Pejchal R, Doores KJ, Walker LM, Khayat R, Huang PS, Wang SK, Stanfield RL, Julien JP, Ramos A, Crispin M, Depetris R, Katpally U, Marozsan A, Cupo A, Maloveste S, Liu Y, McBride R, Ito Y, Sanders RW, Ogohara C, Paulson JC, Feizi T, Scanlan CN, Wong CH, Moore JP, Olson WC, Ward AB, Poignard P, Schief WR, Burton DR, Wilson IA. A potent and broad neutralizing antibody recognizes and penetrates the HIV glycan shield. Science. 2011;334(6059):1097–103. doi: 10.1126/science.1213256. PubMed PMID: 21998254; PMCID: PMC3280215.

8. Bonsignori M, Kreider EF, Fera D, Meyerhoff RR, Bradley T, Wiehe K, Alam SM, Aussedat B, Walkowicz WE, Hwang KK, Saunders KO, Zhang R, Gladden MA, Monroe A, Kumar A, Xia SM, Cooper M, Louder MK, McKee K, Bailer RT, Pier BW, Jette CA, Kelsoe G, Williams WB, Morris L, Kappes J, Wagh K, Kamanga G, Cohen MS, Hraber PT, Montefiori DC, Trama A, Liao HX, Kepler TB, Moody MA, Gao F, Danishefsky SJ, Mascola JR, Shaw GM, Hahn BH, Harrison SC, Korber BT, Haynes BF. Staged induction of HIV-1 glycan-dependent broadly neutralizing antibodies. Sci Transl Med. 2017;9(381). doi: 10.1126/scitranslmed.aai7514. PubMed PMID: 28298420.

9. Walker LM, Simek MD, Priddy F, Gach JS, Wagner D, Zwick MB, Phogat SK, Poignard P, Burton DR. A limited number of antibody specificities mediate broad and potent serum neutralization in selected HIV-1 infected individuals. PLoS Pathog. 2010;6(8):e1001028. Epub 2010/08/05. doi: 10.1371/journal.ppat.1001028. PubMed PMID: 20700449; PMCID: PMC2916884.

10. Walker LM, Phogat SK, Chan-Hui PY, Wagner D, Phung P, Goss JL, Wrin T, Simek MD, Fling S, Mitcham JL, Lehrman JK, Priddy FH, Olsen OA, Frey SM, Hammond PW, Kaminsky S, Zamb T, Moyle M, Koff WC, Poignard P, Burton DR, Investigators PGP. Broad and potent neutralizing antibodies from an African donor reveal a new HIV-1 vaccine target. Science. 2009;326(5950):285–9. doi: 1178746 [pii] 10.1126/science.1178746. PubMed PMID: 19729618.

11. Bonsignori M, Hwang KK, Chen X, Tsao CY, Morris L, Gray E, Marshall DJ, Crump JA, Kapiga SH, Sam NE, Sinangil F, Pancera M, Yongping Y, Zhang B, Zhu J, Kwong PD, O’Dell S, Mascola JR, Wu L, Nabel GJ, Phogat S, Seaman MS, Whitesides JF, Moody MA, Kelsoe G, Yang X, Sodroski J, Shaw GM, Montefiori DC, Kepler TB, Tomaras GD, Alam SM, Liao HX, Haynes BF. Analysis of a clonal lineage of HIV-1 envelope V2/V3 conformational epitope-specific broadly neutralizing antibodies and their inferred unmutated common ancestors. J Virol. 2011;85(19):9998–10009. doi: 10.1128/JVI.05045-11. PubMed PMID: 21795340; PMCID: PMC3196428.

12. Roark RS, Li H, Williams WB, Chug H, Mason RD, Gorman J, Wang S, Lee FH, Rando J, Bonsignori M, Hwang KK, Saunders KO, Wiehe K, Moody MA, Hraber PT, Wagh K, Giorgi EE, Russell RM, Bibollet-Ruche F, Liu W, Connell J, Smith AG, DeVoto J, Murphy AI, Smith J, Ding W, Zhao C, Chohan N, Okumura M, Rosario C, Ding Y, Lindemuth E, Bauer AM, Bar KJ, Ambrozak D, Chao CW, Chuang GY, Geng H, Lin BC, Louder MK, Nguyen R, Zhang B, Lewis MG, Raymond D, Doria-Rose NA, Schramm CA, Douek DC, Roederer M, Kepler TB, Kelsoe G, Mascola JR, Kwong PD, Korber BT, Harrison SC, Haynes BF, Hahn BH, Shaw GM. Recapitulation of HIV-1 Env-antibody coevolution in macaques leading to neutralization breadth. Science. 2020. Epub 2020/11/19. doi: 10.1126/science.abd2638. PubMed PMID: 33214287.

13. Wang Z, Merkenschlager J, Chen ST, Oliveira TY, Ramos V, Gordon KM, Yao KH, Jankovic M, Nussenzweig M, Escolano A. Isolation of single HIV-1 Envelope specific B cells and antibody cloning from immunized rhesus macaques. J Immunol Methods. 2020;478:112734. Epub 20191219. doi: 10.1016/j.jim.2019.112734. PubMed PMID: 31866284; PMCID: PMC6961706.

14. Wiehe K, Bradley T, Meyerhoff RR, Hart C, Williams WB, Easterhoff D, Faison WJ, Kepler TB, Saunders KO, Alam SM, Bonsignori M, Haynes BF. Functional Relevance of Improbable Antibody Mutations for HIV Broadly Neutralizing Antibody Development. Cell Host Microbe. 2018;23(6):759–65.e6. Epub 2018/05/31. doi: 10.1016/j.chom.2018.04.018. PubMed PMID: 29861171; PMCID: PMC6002614.

15. Saunders KO, Wiehe K, Tian M, Acharya P, Bradley T, Alam SM, Go EP, Scearce R, Sutherland L, Henderson R, Hsu AL, Borgnia MJ, Chen H, Lu X, Wu NR, Watts B, Jiang C, Easterhoff D, Cheng HL, McGovern K, Waddicor P, Chapdelaine-Williams A, Eaton A, Zhang J, Rountree W, Verkoczy L, Tomai M, Lewis MG, Desaire HR, Edwards RJ, Cain DW, Bonsignori M, Montefiori D, Alt FW, Haynes BF. Targeted selection of HIV-specific antibody mutations by engineering B cell maturation. Science. 2019;366(6470). doi: 10.1126/science.aay7199. PubMed PMID: 31806786.

16. Henderson R, Zhou Y, Stalls V, Wiehe K, Saunders KO, Wagh K, Anasti K, Barr M, Parks R, Alam SM, Korber B, Haynes BF, Bartesaghi A, Acharya P. Structural basis for breadth development in the HIV-1 V3-glycan targeting DH270 antibody clonal lineage. Nat Commun. 2023;14(1):2782. Epub 20230515. doi: 10.1038/s41467-023-38108-1. PubMed PMID: 37188681; PMCID: PMC10184639.

17. Wiehe K, Saunders KO, Stalls V, Cain DW, Venkatayogi S, Martin Beem JS, Berry M, Evangelous T, Henderson R, Hora B, Xia SM, Jiang C, Newman A, Bowman C, Lu X, Bryan ME, Bal J, Sanzone A, Chen H, Eaton A, Tomai MA, Fox CB, Tam YK, Barbosa C, Bonsignori M, Muramatsu H, Alam SM, Montefiori DC, Williams WB, Pardi N, Tian M, Weissman D, Alt FW, Acharya P, Haynes BF. Mutation-guided vaccine design: A process for developing boosting immunogens for HIV broadly neutralizing antibody induction. Cell Host Microbe. 2024;32(5):693–709.e7. Epub 20240425. doi: 10.1016/j.chom.2024.04.006. PubMed PMID: 38670093.

18. Mu Z, Wiehe K, Saunders KO, Henderson R, Cain DW, Parks R, Martik D, Mansouri K, Edwards RJ, Newman A, Lu X, Xia SM, Eaton A, Bonsignori M, Montefiori D, Han Q, Venkatayogi S, Evangelous T, Wang Y, Rountree W, Korber B, Wagh K, Tam Y, Barbosa C, Alam SM, Williams WB, Tian M, Alt FW, Pardi N, Weissman D, Haynes BF. mRNA-encoded HIV-1 Env trimer ferritin nanoparticles induce monoclonal antibodies that neutralize heterologous HIV-1 isolates in mice. Cell Rep. 2022;38(11):110514. doi: 10.1016/j.celrep.2022.110514. PubMed PMID: 35294883; PMCID: PMC8922439.

19. Steichen JM, Lin YC, Havenar-Daughton C, Pecetta S, Ozorowski G, Willis JR, Toy L, Sok D, Liguori A, Kratochvil S, Torres JL, Kalyuzhniy O, Melzi E, Kulp DW, Raemisch S, Hu X, Bernard SM, Georgeson E, Phelps N, Adachi Y, Kubitz M, Landais E, Umotoy J, Robinson A, Briney B, Wilson IA, Burton DR, Ward AB, Crotty S, Batista FD, Schief WR. A generalized HIV vaccine design strategy for priming of broadly neutralizing antibody responses. Science. 2019;366(6470). Epub 2019/10/31. doi: 10.1126/science.aax4380. PubMed PMID: 31672916; PMCID: PMC7092357.

20. Escolano A, Steichen JM, Dosenovic P, Kulp DW, Golijanin J, Sok D, Freund NT, Gitlin AD, Oliveira T, Araki T, Lowe S, Chen ST, Heinemann J, Yao KH, Georgeson E, Saye-Francisco KL, Gazumyan A, Adachi Y, Kubitz M, Burton DR, Schief WR, Nussenzweig MC. Sequential Immunization Elicits Broadly Neutralizing Anti-HIV-1 Antibodies in Ig Knockin Mice. Cell. 2016;166(6):1445–58.e12. doi: 10.1016/j.cell.2016.07.030. PubMed PMID: 27610569; PMCID: PMC5019122.

21. Escolano A, Gristick HB, Abernathy ME, Merkenschlager J, Gautam R, Oliveira TY, Pai J, West AP, Barnes CO, Cohen AA, Wang H, Golijanin J, Yost D, Keeffe JR, Wang Z, Zhao P, Yao KH, Bauer J, Nogueira L, Gao H, Voll AV, Montefiori DC, Seaman MS, Gazumyan A, Silva M, McGuire AT, Stamatatos L, Irvine DJ, Wells L, Martin MA, Bjorkman PJ, Nussenzweig MC. Immunization expands B cells specific to HIV-1 V3 glycan in mice and macaques. Nature. 2019;570(7762):468–73. Epub 2019/05/29. doi: 10.1038/s41586-019-1250-z. PubMed PMID: 31142836; PMCID: PMC6657810.

22. Havenar-Daughton C, Abbott RK, Schief WR, Crotty S. When designing vaccines, consider the starting material: the human B cell repertoire. Curr Opin Immunol. 2018;53:209–16. Epub 2018/09/03. doi: 10.1016/j.coi.2018.08.002. PubMed PMID: 30190230; PMCID: PMC6148213.

23. Steichen JM, Kulp DW, Tokatlian T, Escolano A, Dosenovic P, Stanfield RL, McCoy LE, Ozorowski G, Hu X, Kalyuzhniy O, Briney B, Schiffner T, Garces F, Freund NT, Gitlin AD, Menis S, Georgeson E, Kubitz M, Adachi Y, Jones M, Mutafyan AA, Yun DS, Mayer CT, Ward AB, Burton DR, Wilson IA, Irvine DJ, Nussenzweig MC, Schief WR. HIV Vaccine Design to Target Germline Precursors of Glycan-Dependent Broadly Neutralizing Antibodies. Immunity. 2016;45(3):483–96. Epub 2016/09/08. doi: 10.1016/j.immuni.2016.08.016. PubMed PMID: 27617678; PMCID: PMC5040827.

24. Li H, Wang S, Kong R, Ding W, Lee FH, Parker Z, Kim E, Learn GH, Hahn P, Policicchio B, Brocca-Cofano E, Deleage C, Hao X, Chuang GY, Gorman J, Gardner M, Lewis MG, Hatziioannou T, Santra S, Apetrei C, Pandrea I, Alam SM, Liao HX, Shen X, Tomaras GD, Farzan M, Chertova E, Keele BF, Estes JD, Lifson JD, Doms RW, Montefiori DC, Haynes BF, Sodroski JG, Kwong PD, Hahn BH, Shaw GM. Envelope residue 375 substitutions in simian-human immunodeficiency viruses enhance CD4 binding and replication in rhesus macaques. Proceedings of the National Academy of Sciences of the United States of America. 2016;113(24):E3413-22. Epub 2016/06/02. doi: 10.1073/pnas.1606636113. PubMed PMID: 27247400; PMCID: PMC4914158.

25. Doria-Rose NA, Schramm CA, Gorman J, Moore PL, Bhiman JN, DeKosky BJ, Ernandes MJ, Georgiev IS, Kim HJ, Pancera M, Staupe RP, Altae-Tran HR, Bailer RT, Crooks ET, Cupo A, Druz A, Garrett NJ, Hoi KH, Kong R, Louder MK, Longo NS, McKee K, Nonyane M, O’Dell S, Roark RS, Rudicell RS, Schmidt SD, Sheward DJ, Soto C, Wibmer CK, Yang Y, Zhang Z, Mullikin JC, Binley JM, Sanders RW, Wilson IA, Moore JP, Ward AB, Georgiou G, Williamson C, Abdool Karim SS, Morris L, Kwong PD, Shapiro L, Mascola JR, Program NCS. Developmental pathway for potent V1V2-directed HIV-neutralizing antibodies. Nature. 2014;509(7498):55–62. doi: 10.1038/nature13036. PubMed PMID: 24590074; PMCID: PMC4395007.

26. Havenar-Daughton C, Lee JH, Crotty S. Tfh cells and HIV bnAbs, an immunodominance model of the HIV neutralizing antibody generation problem. Immunol Rev. 2017;275(1):49–61. doi: 10.1111/imr.12512. PubMed PMID: 28133798.

27. Finney J, Yeh CH, Kelsoe G, Kuraoka M. Germinal center responses to complex antigens. Immunol Rev. 2018;284(1):42–50. doi: 10.1111/imr.12661. PubMed PMID: 29944756; PMCID: PMC6023402.

28. Mesin L, Ersching J, Victora GD. Germinal Center B Cell Dynamics. Immunity. 2016;45(3):471–82. doi: 10.1016/j.immuni.2016.09.001. PubMed PMID: 27653600; PMCID: PMC5123673.

29. Viant C, Weymar GHJ, Escolano A, Chen S, Hartweger H, Cipolla M, Gazumyan A, Nussenzweig MC. Antibody Affinity Shapes the Choice between Memory and Germinal Center B Cell Fates. Cell. 2020;183(5):1298–311.e11. Epub 2020/10/29. doi: 10.1016/j.cell.2020.09.063. PubMed PMID: 33125897; PMCID: PMC7722471.

30. Nakagawa R, Calado DP. Positive Selection in the Light Zone of Germinal Centers. Front Immunol. 2021;12:661678. Epub 20210331. doi: 10.3389/fimmu.2021.661678. PubMed PMID: 33868314; PMCID: PMC8044421.

31. Suan D, Kräutler NJ, Maag JLV, Butt D, Bourne K, Hermes JR, Avery DT, Young C, Statham A, Elliott M, Dinger ME, Basten A, Tangye SG, Brink R. CCR6 Defines Memory B Cell Precursors in Mouse and Human Germinal Centers, Revealing Light-Zone Location and Predominant Low Antigen Affinity. Immunity. 2017;47(6):1142–53.e4. doi: 10.1016/j.immuni.2017.11.022. PubMed PMID: 29262350.

32. De Silva NS, Klein U. Dynamics of B cells in germinal centres. Nat Rev Immunol. 2015;15(3):137–48. Epub 20150206. doi: 10.1038/nri3804. PubMed PMID: 25656706; PMCID: PMC4399774.

33. Young C, Brink R. The unique biology of germinal center B cells. Immunity. 2021;54(8):1652–64. doi: 10.1016/j.immuni.2021.07.015. PubMed PMID: 34380063.

34. Bonsignori M, Liao HX, Gao F, Williams WB, Alam SM, Montefiori DC, Haynes BF. Antibody-virus co-evolution in HIV infection: paths for HIV vaccine development. Immunol Rev. 2017;275(1):145–60. doi: 10.1111/imr.12509. PubMed PMID: 28133802; PMCID: PMC5302796.

35. Haynes BF, Wiehe K, Borrrow P, Saunders KO, Korber B, Wagh K, McMichael AJ, Kelsoe G, Hahn BH, Alt F, Shaw GM. Strategies for HIV-1 vaccines that induce broadly neutralizing antibodies. Nat Rev Immunol. 2022. Epub 20220812. doi: 10.1038/s41577-022-00753-w. PubMed PMID: 35962033; PMCID: PMC9372928.

36. van Schooten J, van Haaren MM, Li H, McCoy LE, Havenar-Daughton C, Cottrell CA, Burger JA, van der Woude P, Helgers LC, Tomris I, Labranche CC, Montefiori DC, Ward AB, Burton DR, Moore JP, Sanders RW, Crotty S, Shaw GM, van Gils MJ. Antibody responses induced by SHIV infection are more focused than those induced by soluble native HIV-1 envelope trimers in non-human primates. PLoS Pathog. 2021;17(8):e1009736. Epub 20210825. doi: 10.1371/journal.ppat.1009736. PubMed PMID: 34432859; PMCID: PMC8423243.

37. Wei X, Ghosh SK, Taylor ME, Johnson VA, Emini EA, Deutsch P, Lifson JD, Bonhoeffer S, Nowak MA, Hahn BH. Viral dynamics in human immunodeficiency virus type 1 infection. Nature. 1995;373(6510):117–22. doi: 10.1038/373117a0. PubMed PMID: 7529365.

38. Ho DD, Neumann AU, Perelson AS, Chen W, Leonard JM, Markowitz M. Rapid turnover of plasma virions and CD4 lymphocytes in HIV-1 infection. Nature. 1995;373(6510):123–6. doi: 10.1038/373123a0. PubMed PMID: 7816094.

39. Markowitz M, Louie M, Hurley A, Sun E, Di Mascio M, Perelson AS, Ho DD. A novel antiviral intervention results in more accurate assessment of human immunodeficiency virus type 1 replication dynamics and T-cell decay in vivo. J Virol. 2003;77(8):5037–8. doi: 10.1128/jvi.77.8.5037-5038.2003. PubMed PMID: 12663814; PMCID: PMC152136.

40. Wei X, Decker JM, Wang S, Hui H, Kappes JC, Wu X, Salazar-Gonzalez JF, Salazar MG, Kilby JM, Saag MS, Komarova NL, Nowak MA, Hahn BH, Kwong PD, Shaw GM. Antibody neutralization and escape by HIV-1. Nature. 2003;422(6929):307–12. doi: 10.1038/nature01470. PubMed PMID: 12646921.

41. Habib R, Roark R, Li H, Connell A, Hogarty M, Wagh K, Wang S, Marchitto L, Skelly A, Carey J, Sowers K, Ayyanathan K, Plante S, Bibollet-Ruche F, Park Y, Agostino C, Singh A, Martella C, Lewis E, Lora J, Ding W, Campion M, Zhao C, Liu W, Li Y, Li X, Liang B, Chowdhury R, Amereh K, Van Itallie E, Sheng Z, Ghosh A, Bar K, Williams W, Wiehe K, Saunders K, Edwards R, Cain D, Lewis M, Batista F, Burton D, Andrabi R, Kulp D, Haynes B, Korber B, Shapiro L, Kwong P, Hahn B, Shaw G. Env-antibody coevolution identifies B cell priming as the principal bottleneck to HIV-1 V2 apex broadly neutralizing antibody development. bioRxiv preprint. 2025. doi: 10.1101/2025.05.03.652068.

42. Salazar-Gonzalez JF, Bailes E, Pham KT, Salazar MG, Guffey MB, Keele BF, Derdeyn CA, Farmer P, Hunter E, Allen S, Manigart O, Mulenga J, Anderson JA, Swanstrom R, Haynes BF, Athreya GS, Korber BT, Sharp PM, Shaw GM, Hahn BH. Deciphering human immunodeficiency virus type 1 transmission and early envelope diversification by single-genome amplification and sequencing. J Virol. 2008;82(8):3952–70. doi: JVI.02660-07 [pii] 10.1128/JVI.02660-07. PubMed PMID: 18256145; PMCID: PMC2293010.

43. Keele BF, Giorgi EE, Salazar-Gonzalez JF, Decker JM, Pham KT, Salazar MG, Sun C, Grayson T, Wang S, Li H, Wei X, Jiang C, Kirchherr JL, Gao F, Anderson JA, Ping LH, Swanstrom R, Tomaras GD, Blattner WA, Goepfert PA, Kilby JM, Saag MS, Delwart EL, Busch MP, Cohen MS, Montefiori DC, Haynes BF, Gaschen B, Athreya GS, Lee HY, Wood N, Seoighe C, Perelson AS, Bhattacharya T, Korber BT, Hahn BH, Shaw GM. Identification and characterization of transmitted and early founder virus envelopes in primary HIV-1 infection. Proc Natl Acad Sci U S A. 2008;105(21):7552–7. Epub 2008/05/19. doi: 10.1073/pnas.0802203105. PubMed PMID: 18490657; PMCID: PMC2387184.

44. Hraber P, Korber B, Wagh K, Giorgi EE, Bhattacharya T, Gnanakaran S, Lapedes AS, Learn GH, Kreider EF, Li Y, Shaw GM, Hahn BH, Montefiori DC, Alam SM, Bonsignori M, Moody MA, Liao HX, Gao F, Haynes BF. Longitudinal Antigenic Sequences and Sites from Intra-Host Evolution (LASSIE) Identifies Immune-Selected HIV Variants. Viruses. 2015;7(10):5443–75. Epub 20151021. doi: 10.3390/v7102881. PubMed PMID: 26506369; PMCID: PMC4632389.

45. Klasse PJ, Ketas TJ, Cottrell CA, Ozorowski G, Debnath G, Camara D, Francomano E, Pugach P, Ringe RP, LaBranche CC, van Gils MJ, Bricault CA, Barouch DH, Crotty S, Silvestri G, Kasturi S, Pulendran B, Wilson IA, Montefiori DC, Sanders RW, Ward AB, Moore JP. Epitopes for neutralizing antibodies induced by HIV-1 envelope glycoprotein BG505 SOSIP trimers in rabbits and macaques. PLoS Pathog. 2018;14(2):e1006913. Epub 20180223. doi: 10.1371/journal.ppat.1006913. PubMed PMID: 29474444; PMCID: PMC5841823.

46. Wagh K, Kreider EF, Li Y, Barbian HJ, Learn GH, Giorgi E, Hraber PT, Decker TG, Smith AG, Gondim MV, Gillis L, Wandzilak J, Chuang GY, Rawi R, Cai F, Pellegrino P, Williams I, Overbaugh J, Gao F, Kwong PD, Haynes BF, Shaw GM, Borrow P, Seaman MS, Hahn BH, Korber B. Completeness of HIV-1 Envelope Glycan Shield at Transmission Determines Neutralization Breadth. Cell Rep. 2018;25(4):893–908.e7. doi: 10.1016/j.celrep.2018.09.087. PubMed PMID: 30355496; PMCID: PMC6426304.

47. Williams WB, Meyerhoff RR, Edwards RJ, Li H, Manne K, Nicely NI, Henderson R, Zhou Y, Janowska K, Mansouri K, Gobeil S, Evangelous T, Hora B, Berry M, Abuahmad AY, Sprenz J, Deyton M, Stalls V, Kopp M, Hsu AL, Borgnia MJ, Stewart-Jones GBE, Lee MS, Bronkema N, Moody MA, Wiehe K, Bradley T, Alam SM, Parks RJ, Foulger A, Oguin T, Sempowski GD, Bonsignori M, LaBranche CC, Montefiori DC, Seaman M, Santra S, Perfect J, Francica JR, Lynn GM, Aussedat B, Walkowicz WE, Laga R, Kelsoe G, Saunders KO, Fera D, Kwong PD, Seder RA, Bartesaghi A, Shaw GM, Acharya P, Haynes BF. Fab-dimerized glycan-reactive antibodies are a structural category of natural antibodies. Cell. 2021. Epub 2021/05/18. doi: 10.1016/j.cell.2021.04.042. PubMed PMID: 34019795; PMCID: PMC8135257.

48. Liao HX, Levesque MC, Nagel A, Dixon A, Zhang R, Walter E, Parks R, Whitesides J, Marshall DJ, Hwang KK, Yang Y, Chen X, Gao F, Munshaw S, Kepler TB, Denny T, Moody MA, Haynes BF. High-throughput isolation of immunoglobulin genes from single human B cells and expression as monoclonal antibodies. J Virol Methods. 2009;158(1-2):171–9. doi: 10.1016/j.jviromet.2009.02.014. PubMed PMID: 19428587; PMCID: PMC2805188.

49. Fan Q, Nelson CS, Bialas KM, Chiuppesi F, Amos J, Gurley TC, Marshall DJ, Eudailey J, Heimsath H, Himes J, Deshpande A, Walter MR, Wussow F, Diamond DJ, Barry PA, Moody MA, Kaur A, Permar SR. Plasmablast Response to Primary Rhesus Cytomegalovirus (CMV) Infection in a Monkey Model of Congenital CMV Transmission. Clin Vaccine Immunol. 2017;24(5). Epub 20170505. doi: 10.1128/CVI.00510-16. PubMed PMID: 28298291; PMCID: PMC5424243.

50. Sok D, Burton DR. Recent progress in broadly neutralizing antibodies to HIV. Nat Immunol. 2018;19(11):1179–88. Epub 20181017. doi: 10.1038/s41590-018-0235-7. PubMed PMID: 30333615; PMCID: PMC6440471.

51. Setliff I, Shiakolas AR, Pilewski KA, Murji AA, Mapengo RE, Janowska K, Richardson S, Oosthuysen C, Raju N, Ronsard L, Kanekiyo M, Qin JS, Kramer KJ, Greenplate AR, McDonnell WJ, Graham BS, Connors M, Lingwood D, Acharya P, Morris L, Georgiev IS. High-Throughput Mapping of B Cell Receptor Sequences to Antigen Specificity. Cell. 2019;179(7):1636–46.e15. Epub 2019/11/28. doi: 10.1016/j.cell.2019.11.003. PubMed PMID: 31787378; PMCID: PMC7158953.

52. deCamp A, Hraber P, Bailer RT, Seaman MS, Ochsenbauer C, Kappes J, Gottardo R, Edlefsen P, Self S, Tang H, Greene K, Gao H, Daniell X, Sarzotti-Kelsoe M, Gorny MK, Zolla-Pazner S, LaBranche CC, Mascola JR, Korber BT, Montefiori DC. Global panel of HIV-1 Env reference strains for standardized assessments of vaccine-elicited neutralizing antibodies. J Virol. 2014;88(5):2489–507. Epub 2013/12/18. doi: 10.1128/JVI.02853-13. PubMed PMID: 24352443; PMCID: PMC3958090.

53. Holmes S, Li H, Shen X, Martin M, Tuck R, Chen Y, Giorgi EE, Kirshner HF, Berry M, Van Italie E, Venkatayogi S, Martin Beem JS, Edwards RJ, Mansouri K, Singh A, Kuykendall C, Gurley T, Anthony Moody M, DeNayer N, Demarco T, Denny TN, Wang Y, Evangelous TD, Clinton JT, Hora B, Wagh K, Seaman MS, Saunders KO, Solomotis N, Misamore J, Lewis MG, Wiehe K, Montefiori DC, Shaw GM, Williams WB. Neonatal immunity associated with heterologous HIV-1 neutralizing antibody induction in SHIV-infected Rhesus Macaques. Nat Commun. 2024;15(1):10302. Epub 20241127. doi: 10.1038/s41467-024-54753-6. PubMed PMID: 39604409; PMCID: PMC11603298.

54. Macosko EZ, Basu A, Satija R, Nemesh J, Shekhar K, Goldman M, Tirosh I, Bialas AR, Kamitaki N, Martersteck EM, Trombetta JJ, Weitz DA, Sanes JR, Shalek AK, Regev A, McCarroll SA. Highly Parallel Genome-wide Expression Profiling of Individual Cells Using Nanoliter Droplets. Cell. 2015;161(5):1202–14. doi: 10.1016/j.cell.2015.05.002. PubMed PMID: 26000488; PMCID: PMC4481139.

55. Satija R, Farrell JA, Gennert D, Schier AF, Regev A. Spatial reconstruction of single-cell gene expression data. Nat Biotechnol. 2015;33(5):495–502. Epub 2015/04/14. doi: 10.1038/nbt.3192. PubMed PMID: 25867923; PMCID: PMC4430369.

56. Stuart T, Butler A, Hoffman P, Hafemeister C, Papalexi E, Mauck WM, Hao Y, Stoeckius M, Smibert P, Satija R. Comprehensive Integration of Single-Cell Data. Cell. 2019;177(7):1888–902.e21. Epub 20190606. doi: 10.1016/j.cell.2019.05.031. PubMed PMID: 31178118; PMCID: PMC6687398.

57. Axelsson S, Magnuson A, Lange A, Alshamari A, Hörnquist EH, Hultgren O. A combination of the activation marker CD86 and the immune checkpoint marker B and T lymphocyte attenuator (BTLA) indicates a putative permissive activation state of B cell subtypes in healthy blood donors independent of age and sex. BMC Immunol. 2020;21(1):14. Epub 20200320. doi: 10.1186/s12865-020-00343-2. PubMed PMID: 32197584; PMCID: PMC7082969.

58. Syeda MZ, Hong T, Huang C, Huang W, Mu Q. B cell memory: from generation to reactivation: a multipronged defense wall against pathogens. Cell Death Discov. 2024;10(1):117. Epub 20240307. doi: 10.1038/s41420-024-01889-5. PubMed PMID: 38453885; PMCID: PMC10920759.

59. Knox JJ, Buggert M, Kardava L, Seaton KE, Eller MA, Canaday DH, Robb ML, Ostrowski MA, Deeks SG, Slifka MK, Tomaras GD, Moir S, Moody MA, Betts MR. T-bet+ B cells are induced by human viral infections and dominate the HIV gp140 response. JCI Insight. 2017;2(8). Epub 20170420. doi: 10.1172/jci.insight.92943. PubMed PMID: 28422752; PMCID: PMC5396521.

60. Pae J, Ersching J, Castro TBR, Schips M, Mesin L, Allon SJ, Ordovas-Montanes J, Mlynarczyk C, Melnick A, Efeyan A, Shalek AK, Meyer-Hermann M, Victora GD. Cyclin D3 drives inertial cell cycling in dark zone germinal center B cells. J Exp Med. 2021;218(4). doi: 10.1084/jem.20201699. PubMed PMID: 33332554; PMCID: PMC7754672.

61. Kennedy DE, Okoreeh MK, Maienschein-Cline M, Ai J, Veselits M, McLean KC, Dhungana Y, Wang H, Peng J, Chi H, Mandal M, Clark MR. Novel specialized cell state and spatial compartments within the germinal center. Nat Immunol. 2020;21(6):660–70. Epub 20200427. doi: 10.1038/s41590-020-0660-2. PubMed PMID: 32341509; PMCID: PMC7255947.

62. Schramm CA, Sheng Z, Zhang Z, Mascola JR, Kwong PD, Shapiro L. SONAR: A High-Throughput Pipeline for Inferring Antibody Ontogenies from Longitudinal Sequencing of B Cell Transcripts. Front Immunol. 2016;7:372. Epub 20160921. doi: 10.3389/fimmu.2016.00372. PubMed PMID: 27708645; PMCID: PMC5030719.

63. Corcoran MM, Phad GE, Vázquez Bernat, Stahl-Hennig C, Sumida N, Persson MA, Martin M, Karlsson Hedestam GB. Production of individualized V gene databases reveals high levels of immunoglobulin genetic diversity. Nat Commun. 2016;7:13642. Epub 2016/12/20. doi: 10.1038/ncomms13642. PubMed PMID: 27995928; PMCID: PMC5187446.

64. Williams WB, Wiehe K, Saunders KO, Haynes BF. Strategies for induction of HIV-1 envelope-reactive broadly neutralizing antibodies. J Int AIDS Soc. 2021;24 Suppl 7:e25831. doi: 10.1002/jia2.25831. PubMed PMID: 34806332; PMCID: PMC8606870.

65. Barnes CO, Gristick HB, Freund NT, Escolano A, Lyubimov AY, Hartweger H, West AP, Cohen AE, Nussenzweig MC, Bjorkman PJ. Structural characterization of a highly-potent V3- glycan broadly neutralizing antibody bound to natively-glycosylated HIV-1 envelope. Nat Commun. 2018;9(1):1251. Epub 2018/03/28. doi: 10.1038/s41467-018-03632-y. PubMed PMID: 29593217; PMCID: PMC5871869.

66. Wang Z, Barnes CO, Gautam R, Cetrulo Lorenzi JC, Mayer CT, Oliveira TY, Ramos V, Cipolla M, Gordon KM, Gristick HB, West AP, Nishimura Y, Raina H, Seaman MS, Gazumyan A, Martin M, Bjorkman PJ, Nussenzweig MC, Escolano A. A broadly neutralizing macaque monoclonal antibody against the HIV-1 V3-Glycan patch. Elife. 2020;9. Epub 20201021. doi: 10.7554/eLife.61991. PubMed PMID: 33084569; PMCID: PMC7577740.

67. Cirelli KM, Carnathan DG, Nogal B, Martin JT, Rodriguez OL, Upadhyay AA, Enemuo CA, Gebru EH, Choe Y, Viviano F, Nakao C, Pauthner MG, Reiss S, Cottrell CA, Smith ML, Bastidas R, Gibson W, Wolabaugh AN, Melo MB, Cossette B, Kumar V, Patel NB, Tokatlian T, Menis S, Kulp DW, Burton DR, Murrell B, Schief WR, Bosinger SE, Ward AB, Watson CT, Silvestri G, Irvine DJ, Crotty S. Slow Delivery Immunization Enhances HIV Neutralizing Antibody and Germinal Center Responses via Modulation of Immunodominance. Cell. 2020;180(1):206. doi: 10.1016/j.cell.2019.12.027. PubMed PMID: 31923396; PMCID: PMC7009795.

68. Havenar-Daughton C, Carnathan DG, Boopathy AV, Upadhyay AA, Murrell B, Reiss SM, Enemuo CA, Gebru EH, Choe Y, Dhadvai P, Viviano F, Kaushik K, Bhiman JN, Briney B, Burton DR, Bosinger SE, Schief WR, Irvine DJ, Silvestri G, Crotty S. Rapid Germinal Center and Antibody Responses in Non-human Primates after a Single Nanoparticle Vaccine Immunization. Cell Rep. 2019;29(7):1756–66.e8. doi: 10.1016/j.celrep.2019.10.008. PubMed PMID: 31722194; PMCID: PMC6905039.

69. McCoy LE, van Gils MJ, Ozorowski G, Messmer T, Briney B, Voss JE, Kulp DW, Macauley MS, Sok D, Pauthner M, Menis S, Cottrell CA, Torres JL, Hsueh J, Schief WR, Wilson IA, Ward AB, Sanders RW, Burton DR. Holes in the Glycan Shield of the Native HIV Envelope Are a Target of Trimer-Elicited Neutralizing Antibodies. Cell Rep. 2016;16(9):2327–38. doi: 10.1016/j.celrep.2016.07.074. PubMed PMID: 27545891; PMCID: PMC5007210.

70. Pauthner M, Havenar-Daughton C, Sok D, Nkolola JP, Bastidas R, Boopathy AV, Carnathan DG, Chandrashekar A, Cirelli KM, Cottrell CA, Eroshkin AM, Guenaga J, Kaushik K, Kulp DW, Liu J, McCoy LE, Oom AL, Ozorowski G, Post KW, Sharma SK, Steichen JM, de Taeye SW, Tokatlian T, Torrents de la Peña A, Butera ST, LaBranche CC, Montefiori DC, Silvestri G, Wilson IA, Irvine DJ, Sanders RW, Schief WR, Ward AB, Wyatt RT, Barouch DH, Crotty S, Burton DR. Elicitation of Robust Tier 2 Neutralizing Antibody Responses in Nonhuman Primates by HIV Envelope Trimer Immunization Using Optimized Approaches. Immunity. 2017;46(6):1073–88.e6. doi: 10.1016/j.immuni.2017.05.007. PubMed PMID: 28636956; PMCID: PMC5483234.

71. Pauthner MG, Nkolola JP, Havenar-Daughton C, Murrell B, Reiss SM, Bastidas R, Prévost J, Nedellec R, von Bredow B, Abbink P, Cottrell CA, Kulp DW, Tokatlian T, Nogal B, Bianchi M, Li H, Lee JH, Butera ST, Evans DT, Hangartner L, Finzi A, Wilson IA, Wyatt RT, Irvine DJ, Schief WR, Ward AB, Sanders RW, Crotty S, Shaw GM, Barouch DH, Burton DR. Vaccine-Induced Protection from Homologous Tier 2 SHIV Challenge in Nonhuman Primates Depends on Serum-Neutralizing Antibody Titers. Immunity. 2019;50(1):241–52.e6. Epub 2018/12/11. doi: 10.1016/j.immuni.2018.11.011. PubMed PMID: 30552025; PMCID: PMC6335502.

72. Yang YR, McCoy LE, van Gils MJ, Andrabi R, Turner HL, Yuan M, Cottrell CA, Ozorowski G, Voss J, Pauthner M, Polveroni TM, Messmer T, Wilson IA, Sanders RW, Burton DR, Ward AB. Autologous Antibody Responses to an HIV Envelope Glycan Hole Are Not Easily Broadened in Rabbits. J Virol. 2020;94(7). Epub 2020/03/17. doi: 10.1128/JVI.01861-19. PubMed PMID: 31941772; PMCID: PMC7081899.

73. Nelson AN, Shen X, Vekatayogi S, Zhang S, Ozorowski G, Dennis M, Sewall LM, Milligan E, Davis D, Cross KA, Chen Y, van Schooten J, Eudailey J, Isaac J, Memon S, Weinbaum C, Gao H, Stanfield-Oakley S, Byrd A, Chutkan S, Berendam S, Cronin K, Yasmeen A, Alam S, LaBranche CC, Rogers K, Shirreff L, Cupo A, Derking R, Villinger F, Klasse PJ, Ferrari G, Williams WB, Hudgens MG, Ward AB, Montefiori DC, Van Rompay KKA, Wiehe K, Moore JP, Sanders RW, De Paris K, Permar SR. Immunization with germ line-targeting SOSIP trimers elicits broadly neutralizing antibody precursors in infant macaques. Sci Immunol. 2024;9(98):eadm7097. Epub 20240830. doi: 10.1126/sciimmunol.adm7097. PubMed PMID: 39213340.

74. Haynes BF, Saunders KO, Hahn BH, Wiehe K, Baden LR, Shaw GM. Status of HIV vaccine development: progress and promise. J Int AIDS Soc. 2025;28(5):e26489. doi: 10.1002/jia2.26489. PubMed PMID: 40375607; PMCID: PMC12081836.

75. Haynes BF, Wiehe K, Alam SM, Weissman D, Saunders KO. Progress with induction of HIV broadly neutralizing antibodies in the Duke Consortia for HIV/AIDS Vaccine Development. Curr Opin HIV AIDS. 2023;18(6):300–8. Epub 20230925. doi: 10.1097/COH.0000000000000820. PubMed PMID: 37751363; PMCID: PMC10552807.

76. Saunders KO, Edwards RJ, Tilahun K, Manne K, Lu X, Cain DW, Wiehe K, Williams WB, Mansouri K, Hernandez GE, Sutherland L, Scearce R, Parks R, Barr M, DeMarco T, Eater CM, Eaton A, Morton G, Mildenberg B, Wang Y, Rountree RW, Tomai MA, Fox CB, Moody MA, Alam SM, Santra S, Lewis MG, Denny TN, Shaw GM, Montefiori DC, Acharya P, Haynes BF. Stabilized HIV-1 envelope immunization induces neutralizing antibodies to the CD4bs and protects macaques against mucosal infection. Sci Transl Med. 2022;14(661):eabo5598. Epub 20220907. doi: 10.1126/scitranslmed.abo5598. PubMed PMID: 36070369.

77. Saunders KO, Counts J, Thakur B, Stalls V, Edwards R, Manne K, Lu X, Mansouri K, Chen Y, Parks R, Barr M, Sutherland L, Bal J, Havill N, Chen H, Machiele E, Jamieson N, Hora B, Kopp M, Janowska K, Anasti K, Jiang C, Van Itallie E, Venkatayogi S, Eaton A, Henderson R, Barbosa C, Alam SM, Santra S, Weissman D, Moody MA, Cain DW, Tam YK, Lewis M, Williams WB, Wiehe K, Montefiori DC, Acharya P, Haynes BF. Vaccine induction of CD4-mimicking HIV-1 broadly neutralizing antibody precursors in macaques. Cell. 2024;187(1):79–94.e24. doi: 10.1016/j.cell.2023.12.002. PubMed PMID: 38181743.

78. Kumar S, Moral-Sánchez ID, Singh S, Newby ML, Allen JD, Bijl TPL, Vaghani Y, Jing L, Lodha R, Ortlund EA, Crispin M, Patel A, Sanders RW, Luthra K. The Design and Immunogenicity of an HIV-1 Clade C Pediatric Envelope Glycoprotein Stabilized by Multiple Platforms. Vaccines (Basel). 2025;13(2). Epub 20250122. doi: 10.3390/vaccines13020110. PubMed PMID: 40006657; PMCID: PMC11860714.

79. Li D, Edwards RJ, Manne K, Martinez DR, Schäfer A, Alam SM, Wiehe K, Lu X, Parks R, Sutherland LL, Oguin TH, McDanal C, Perez LG, Mansouri K, Gobeil SMC, Janowska K, Stalls V, Kopp M, Cai F, Lee E, Foulger A, Hernandez GE, Sanzone A, Tilahun K, Jiang C, Tse LV, Bock KW, Minai M, Nagata BM, Cronin K, Gee-Lai V, Deyton M, Barr M, Von Holle T, Macintyre AN, Stover E, Feldman J, Hauser BM, Caradonna TM, Scobey TD, Rountree W, Wang Y, Moody MA, Cain DW, DeMarco CT, Denny TN, Woods CW, Petzold EW, Schmidt AG, Teng IT, Zhou T, Kwong PD, Mascola JR, Graham BS, Moore IN, Seder R, Andersen H, Lewis MG, Montefiori DC, Sempowski GD, Baric RS, Acharya P, Haynes BF, Saunders KO. In vitro and in vivo functions of SARS-CoV-2 infection-enhancing and neutralizing antibodies. Cell. 2021;184(16):4203–19.e32. Epub 20210618. doi: 10.1016/j.cell.2021.06.021. PubMed PMID: 34242577; PMCID: PMC8232969.

80. Williams WB, Alam SM, Ofek G, Erdmann N, Montefiori DC, Seaman MS, Wagh K, Korber B, Edwards RJ, Mansouri K, Eaton A, Cain DW, Martin M, Hwang J, Arus-Altuz A, Lu X, Cai F, Jamieson N, Parks R, Barr M, Foulger A, Anasti K, Patel P, Sammour S, Parsons RJ, Huang X, Lindenberger J, Fetics S, Janowska K, Niyongabo A, Janus BM, Astavans A, Fox CB, Mohanty I, Evangelous T, Chen Y, Berry M, Kirshner H, Van Itallie E, Saunders KO, Wiehe K, Cohen KW, McElrath MJ, Corey L, Acharya P, Walsh SR, Baden LR, Haynes BF. Vaccine induction of heterologous HIV-1-neutralizing antibody B cell lineages in humans. Cell. 2024;187(12):2919–34.e20. Epub 20240517. doi: 10.1016/j.cell.2024.04.033. PubMed PMID: 38761800.

81. Kepler TB, Munshaw S, Wiehe K, Zhang R, Yu JS, Woods CW, Denny TN, Tomaras GD, Alam SM, Moody MA, Kelsoe G, Liao HX, Haynes BF. Reconstructing a B-Cell Clonal Lineage. II. Mutation, Selection, and Affinity Maturation. Front Immunol. 2014;5:170. doi: 10.3389/fimmu.2014.00170. PubMed PMID: 24795717; PMCID: PMC4001017.

82. Ramesh A, Darko S, Hua A, Overman G, Ransier A, Francica JR, Trama A, Tomaras GD, Haynes BF, Douek DC, Kepler TB. Structure and Diversity of the Rhesus Macaque Immunoglobulin Loci through Multiple. Front Immunol. 2017;8:1407. Epub 2017/10/27. doi: 10.3389/fimmu.2017.01407. PubMed PMID: 29163486; PMCID: PMC5663730.

83. Stuart T, Butler A, Hoffman P, Hafemeister C, Papalexi E, Mauck WM, 3rd, Hao Y, Stoeckius M, Smibert P, Satija R. Comprehensive Integration of Single-Cell Data. Cell. 2019;177(7):1888–902 e21. Epub 2019/06/11. doi: 10.1016/j.cell.2019.05.031. PubMed PMID: 31178118; PMCID: PMC6687398.

84. McDavid A, Finak G, Chattopadyay PK, Dominguez M, Lamoreaux L, Ma SS, Roederer M, Gottardo R. Data exploration, quality control and testing in single-cell qPCR-based gene expression experiments. Bioinformatics. 2013;29(4):461–7. Epub 2012/12/26. doi: 10.1093/bioinformatics/bts714. PubMed PMID: 23267174; PMCID: PMC3570210.

85. Montefiori DC. Measuring HIV neutralization in a luciferase reporter gene assay. Methods Mol Biol. 2009;485:395–405. doi: 10.1007/978-1-59745-170-3_26. PubMed PMID: 19020839.

86. Punjani A, Rubinstein JL, Fleet DJ, Brubaker MA. cryoSPARC: algorithms for rapid unsupervised cryo-EM structure determination. Nat Methods. 2017;14(3):290–6. Epub 20170206. doi: 10.1038/nmeth.4169. PubMed PMID: 28165473.

87. Goddard TD, Huang CC, Meng EC, Pettersen EF, Couch GS, Morris JH, Ferrin TE. UCSF ChimeraX: Meeting modern challenges in visualization and analysis. Protein Sci. 2018;27(1):14–25. Epub 2017/09/06. doi: 10.1002/pro.3235. PubMed PMID: 28710774; PMCID: PMC5734306.

88. Emsley P, Lohkamp B, Scott WG, Cowtan K. Features and development of Coot. Acta Crystallogr D Biol Crystallogr. 2010;66(Pt 4):486–501. Epub 2010/03/24. doi: 10.1107/S0907444910007493. PubMed PMID: 20383002; PMCID: PMC2852313.

89. Afonine PV, Poon BK, Read RJ, Sobolev OV, Terwilliger TC, Urzhumtsev A, Adams PD. Real-space refinement in PHENIX for cryo-EM and crystallography. Acta Crystallogr D Struct Biol. 2018;74(Pt 6):531–44. Epub 20180530. doi: 10.1107/S2059798318006551. PubMed PMID: 29872004; PMCID: PMC6096492.

90. Otwinowski Z, Minor W. Processing of X-ray diffraction data collected in oscillation mode. Methods Enzymol. 1997;276:307–26. PubMed PMID: 27754618.

91. McCoy AJ, Grosse-Kunstleve RW, Adams PD, Winn MD, Storoni LC, Read RJ. Phaser crystallographic software. J Appl Crystallogr. 2007;40(Pt 4):658–74. Epub 20070713. doi: 10.1107/S0021889807021206. PubMed PMID: 19461840; PMCID: PMC2483472.

92. Adams PD, Afonine PV, Bunkóczi G, Chen VB, Davis IW, Echols N, Headd JJ, Hung LW, Kapral GJ, Grosse-Kunstleve RW, McCoy AJ, Moriarty NW, Oeffner R, Read RJ, Richardson DC, Richardson JS, Terwilliger TC, Zwart PH. PHENIX: a comprehensive Python-based system for macromolecular structure solution. Acta Crystallogr D Biol Crystallogr. 2010;66(Pt 2):213–21. doi: 10.1107/S0907444909052925. PubMed PMID: 20124702; PMCID: PMC2815670.

93. Liebschner D, Afonine PV, Baker ML, Bunkóczi G, Chen VB, Croll TI, Hintze B, Hung LW, Jain S, McCoy AJ, Moriarty NW, Oeffner RD, Poon BK, Prisant MG, Read RJ, Richardson JS, Richardson DC, Sammito MD, Sobolev OV, Stockwell DH, Terwilliger TC, Urzhumtsev AG, Videau LL, Williams CJ, Adams PD. Macromolecular structure determination using X-rays, neutrons and electrons: recent developments in Phenix. Acta Crystallogr D Struct Biol. 2019;75(Pt 10):861–77. Epub 20191002. doi: 10.1107/S2059798319011471. PubMed PMID: 31588918; PMCID: PMC6778852.

94. Krissinel E, Henrick K. Inference of macromolecular assemblies from crystalline state. J Mol Biol. 2007;372(3):774–97. Epub 20070513. doi: 10.1016/j.jmb.2007.05.022. PubMed PMID: 17681537.

